# Chromatin interaction analysis with updated ChIA-PET Tool (V3)

**DOI:** 10.1101/627257

**Authors:** Guoliang Li, Tongkai Sun, Huidan Chang, Liuyang Cai, Ping Hong, Qiangwei Zhou

## Abstract

Understanding chromatin interactions is important since they create chromosome conformation and link the cis- and trans-regulatory elements to their target genes for transcriptional regulation. Chromatin Interaction Analysis with Paired-End Tag (ChIA-PET) sequencing is a genome-wide high-throughput technology that detects chromatin interactions associated with a specific protein of interest. Previously we developed ChIA-PET Tool in 2010 for ChIA-PET data analysis. Here we present the updated version of ChIA-PET Tool (V3), is a computational package to process the next-generation sequence data generated from ChIA-PET experiments. It processes the short-read data and long-read ChIA-PET data with multithreading and generates the statistics of results in a HTML file. In this paper, we provide a detailed demonstration of the design of ChIA-PET Tool V3 and how to install it and analyze a specific ChIA-PET data set with it. At present, other ChIA-PET data analysis tools have developed including ChiaSig, MICC, Mango and ChIA-PET2 and so on. We compared our tool with other tools using the same public data set in the same machine. Most of peaks detected by ChIA-PET Tool V3 overlap with those from other tools. There is higher enrichment for significant chromatin interactions of ChIA-PET Tool V3 in APA plot. ChIA-PET Tool V3 is open source and is available at GitHub (https://github.com/GuoliangLi-HZAU/ChIA-PET_Tool_V3/).

## 1. Introduction

In eukaryotic cells, the genomes are packaged into the micron-sized nucleus with chromatin as the basic unit. Such packing genomes into three-dimensional structure with chromatin interactions is important for DNA replication, repair, gene transcription, and other biological functions. Due to the significance of chromatin interactions in biology in general and transcription regulation in particular, National Institute of Health (NIH) initiated NIH Common Fund: 4D Nucleome (4DN) Program to study the three-dimensional genome structure and its spatiotemporal dynamics. In 4DN program, chromatin interaction is an essential part and Chromatin Interaction Analysis with Paired-End Tag (ChIA-PET) sequencing method [1] is one of the high-throughput methods to generate chromatin interaction data with next-generation sequencing technology.

ChIA-PET method is a derivative of Chromosome Conformation Capture (3C) method [2], and is a genome-wide, high-throughput, high-resolution method to detect chromatin interactions associated with a specific protein of interest, and has been used in a number of applications. The ChIA-PET wet-lab experiment is based on the idea that the proximal DNA fragments from the same cross-linked molecular complexes can be ligated together [3] and it comprises a few basic steps (Figure 1, adapted from Li et al., 2010): cross-linking of the molecules inside the nucleus, shearing the chromatin, precipitating molecules with some antibody of a protein of interest to pull down DNAs associated with the protein, dividing the samples into two aliquots and adding different linkers to two aliquots for linker ligation, combining the two aliquots for proximal ligation of DNAs from the individual molecules, de-cross-linking the proteins from DNAs, digesting the DNAs with enzyme MmeI, Polymerase Chain Reaction (PCR) amplification, and purifying the final sample for next-generation sequencing. The DNA constructs can be sequenced with the high-throughput DNA sequencing facilities in paired-end mode, to generate tens to hundreds of millions of 2×36bp paired-end tag (PET) sequences, or can be sequenced as single reads with read length more than 78bp.

**Figure 1.**
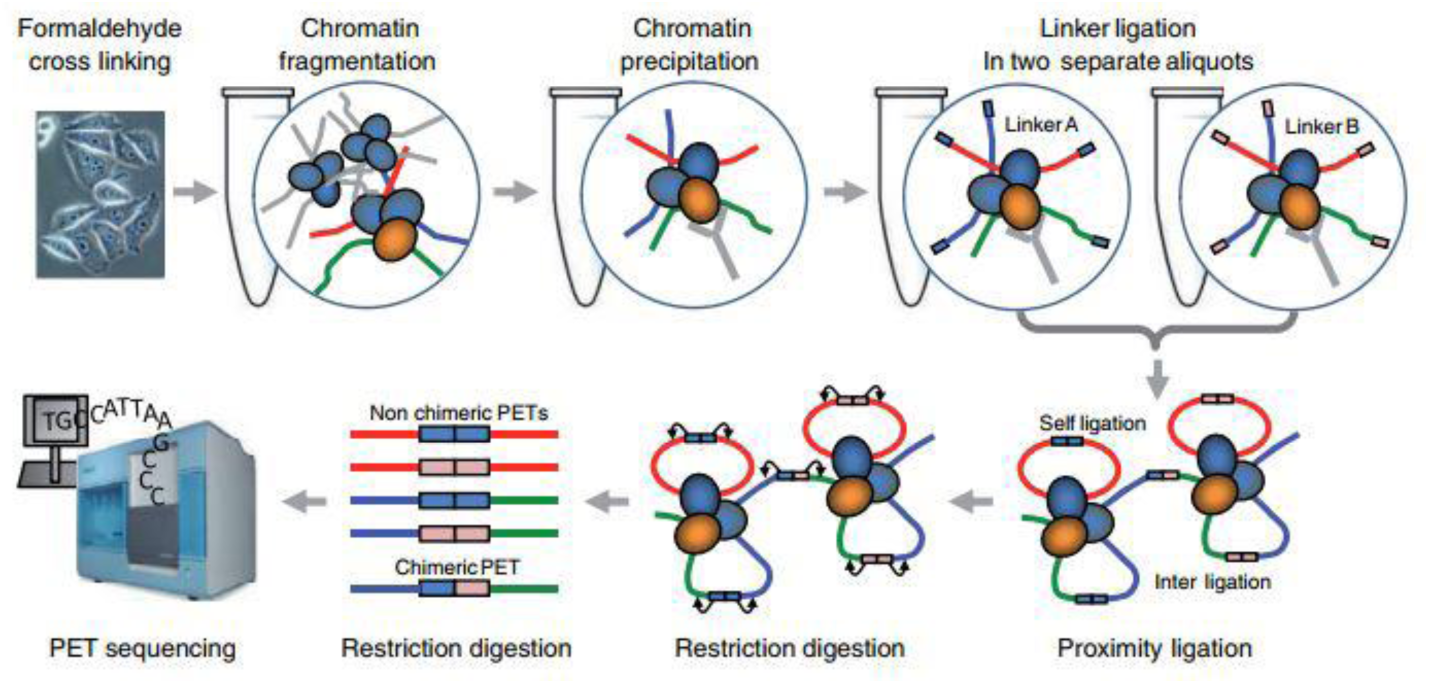
Schematic of ChIA-PET experiment steps, adapted from Li et al., 2010.

In 2015, Tang et al. improved the original method by developing long-read ChIA-PET [4]. The long reads facilitate higher mapping confidence and base pair coverage. The long-read ChIA-PET experiment includes following steps (Figure 2, adapted from Tang et al., 2015): millions of cells were fixed in PBS buffer. Next, formaldehyde was added to cross-link the cells and then neutralized. The cross-linked cells were lysed by cell lysis buffer and nuclear lysis buffer. Chromatin was subjected to fragmentation with an average length of 300 bp by sonication. The specific antibody was used to enrich chromatin fragments. After performing the end-repair and A-tailing using T4 DNA polymerase and Klenow enzyme, the Chromatin Immunoprecipitation (ChIP) DNA ends were proximity-ligated by the single biotinylated bridge-linker with the 3’ nucleotide T over-hanging on both strands. Proximity ligation DNA was reverse cross-linked and fragmented and added sequencing adaptors simultaneously by using Tn5 transposase. DNA fragments contained the bridge linker at ligation junctions were captured by Streptividin beads, and used as templates for PCR amplification. These DNA products were paired-end sequencing (2×150 bp) using Illumina Hi-Seq 2500.

**Figure 2.**
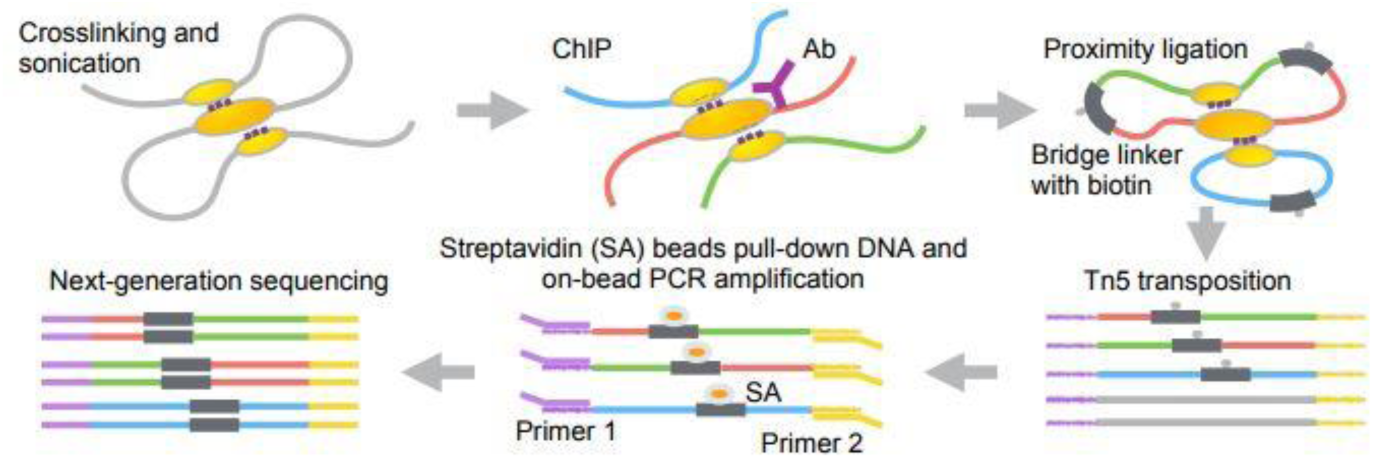
Schematic of long-read ChIA-PET experiment steps, adapted from Tang et al., 2015.

ChIA-PET Tool V3 is a computational package to process the DNA sequence data generated from ChIA-PET experimental method, and it consists of 7 steps (Figure 3): (1) linker filtering, (2) mapping the paired-end reads to a reference genome, (3) purifying the mapped reads, (4) dividing the mapped reads into different categories, (5) peak calling from self-ligation PETs, (6) interaction calling from inter-ligation PETs, and (7) visualizing the results. It was originally published in the journal Genome Biology in 2010. The original ChIA-PET Tool was designed to process the data from 2 × 36bp paired-end sequencing mode. With the advance in sequencing technology, the cost of sequencing is reduced, the amount of data increases at the same time. In the long-read ChIA-PET, the length of the paired-end tags is up to 2 × 250bp [5]. Therefore, we modify ChIA-PET Tool which includes integrating the pipeline of processing short-read data and long-read data, rewriting the shell script of the processing flow with Java, revising the linker filtering with multithreading, adjusting the step of mapping to reduce running time, generating the statistics of the result, and evaluating the quality of the data. In this protocol, we demonstrate how to apply the ChIA-PET Tool V3 to the public ChIA-PET data and illustrate the details and interpretation of the results to facilitate the usage of ChIA-PET Tool V3.

**Figure 3.**
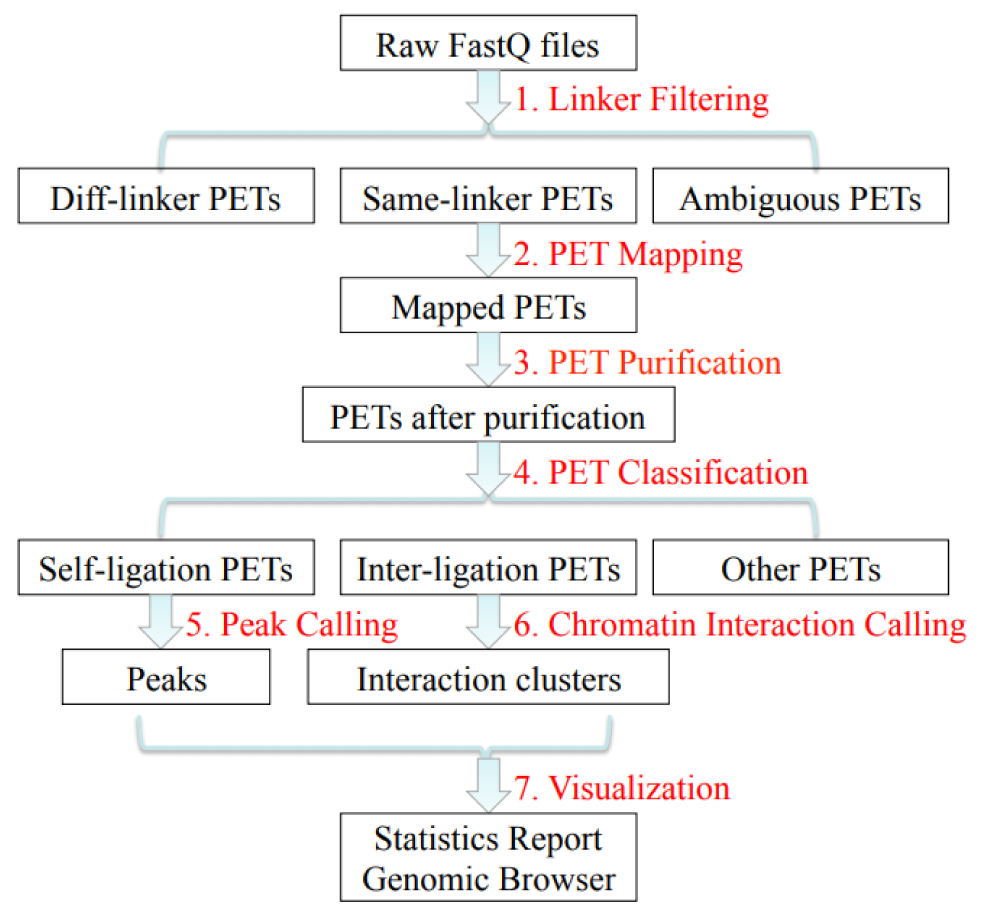
Flowchart of ChIA-PET Tool V3 for ChIA-PET data analysis.

## 2. Materials and Methods

### 2.1. EQUIPMENT

#### 2.1.1. Hardware

Our example of short read ChIA-PET data was run on CentOS release 7.3.1611 with Intel(R) Xeon(R) CPU E5-2630 0 @ 2.30GHz.

Our example of long read ChIA-PET data was run on CentOS release 6.6 with Intel(R) Xeon(R) CPU E5-2620 v3 @ 2.40GHz.

#### 2.1.2. Required supporting software

Java is a popular platform-independent programming language and can be run on any machines with a JVM. BWA [6,7] is used to map ChIA-PET sequencing reads to a reference genome. SAMtools [8] is used to convert the alignment output from SAM format to BAM format. BEDTools [9] is required to convert the files from BAM format to BED format. R environment and its packages are used to compute the p-values in peak calling and interaction calling and generate the graphs for visualization.

Download the supporting software listed as follows:

1. JDK>=1.8 (https://www.oracle.com/technetwork/java/javase/downloads/index.html)
2. BWA (http://bio-bwa.sourceforge.net/)
3. SAMtools (http://samtools.sourceforge.net/)
4. BEDTools (https://bedtools.readthedocs.io/en/latest/)
5. R (https://www.r-project.org/)
6. R package grid (install.packages(“grid”))
7. R package xtable (install.packages(“xtable”))
8. R package RCircos (install.packages(“RCircos”))

Download the ChIA-PET Tool V3 package and unpack it.

#### 2.1.3. Required supporting data

To run ChIA-PET Tool V3, the linker sequence, the genome sequence and its mapping index, the lengths of the individual chromosomes, and cytoband data of the interested genome are required. In this protocol, human reference genome hg19, chromosome size data and cytoband data were downloaded from UCSC website. The random sequences in the genome were removed before further processing. BWA was used to build genome index for mapping sequence reads.

human reference genome:

(ftp://hgdownload.cse.ucsc.edu/goldenPath/hg19/bigZips/chromFa.tar.gz)

chromosome size data:

(ftp://hgdownload.cse.ucsc.edu/goldenPath/hg19/bigZips/hg19.chrom.sizes)

cytoband data:

(http://hgdownload.soe.ucsc.edu/goldenPath/hg19/database/cytoBandIdeo.txt.gz)

#### 2.1.4. Example ChIA-PET data

For test data of short read, ChIA-PET data associated with RNA polymerase II (RNAPII) from human K562 cells was downloaded from NCBI GEO with accession numbers GSM832464 and GSM832465 and dumped to FASTQ format. The raw reads from these two replicates were combined for further processing.

GSM832464 (https://www.ncbi.nlm.nih.gov/geo/query/acc.cgi?acc=GSM832464)

GSM832465 (https://www.ncbi.nlm.nih.gov/geo/query/acc.cgi?acc=GSM832465)

For test data of long read, we use CTCF ChIA-PET data from human GM12878 cells.

GSM1872886 (https://www.ncbi.nlm.nih.gov/geo/query/acc.cgi?acc=GSM1872886)

### 2.2. PROCEDURE

ChIA-PET Tool V3 is an easy-to-use pipeline and we can simply run it using one command line with some options (Appendix A). Users must set the 10 necessary options while other options have default values. Especially, the directories of data should be set properly to make sure that the programs could run smoothly. ChIA-PET Tool V3 will create a folder named by “OUTPUT_PREFIX” in the “OUTPUT_DIRECTORY”. The default value of “OUTPUT_DIRECTORY” is the master folder “ChIA-PET_Tool_V3/”, and the default value of “OUTPUT_PREFIX” is “out”.

ChIA-PET Tool V3 includes 7 steps for data processing (Figure 3). We can use the option “start_step” to select to start with which step. The following sections illustrate the main steps in ChIA-PET Tool V3.

#### 2.2.1. Linker filtering (Timing 7.45 mins)

--output: specifies the directory to store the output data from ChIA-PET Tool V3. Default: ChIA-PET_Tool_V3/output.

--prefix: specifies the prefix of all the output files. Default: out.

--mode: There are two modes for ChIA-PET Tool V3. 0 for short read, 1 for long read.

--fastq1: The path and file name of read1 fastq file in pair-end sequencing.

--fastq2: The path and file name of read2 fastq file in pair-end sequencing.

--linker: linker file.

--minimum_linker_alignment_score: Specifies the allowed minimum alignment score.

--minimum_tag_length: Specifies the minimum tag length. Tag is the sequence after linker removal. This parameter is better to be set above 18bp. Default: 18.

--maximum_tag_length: Specifies the maximum tag length. Default: 1000.

--minSecondBestScoreDiff: Specifies the score difference between the best-aligned and the second-best aligned linkers. Default: 3.

--output_data_with_ambiguous_linker_info: Determines whether to print the linker-ambiguity PETs. Value 0 means not to print these reads to specific files, while Value 1 means to print the PETs with ambiguous linkers. Default: 1.

--thread: the number of threads used in linker filtering and mapping. Default: 1.

ChIA-PET Tool V3 could process short-read data or long-read data by setting parameter “mode”. When the linker sequences and lengths are changed, the parameters should be changed accordingly.

If we select to process short-read data, the start position of the barcode and the length of the barcode would be calculated in the program according to the linker sequences.

By the design, the constructs from ChIA-PET protocol are 78bp, consisting of two tags of 20bp from interacting DNA fragments after MmeI digestion and one full linker of 38bp. With 2 × 36bp paired-end sequencing mode, reads of 36bp will be sequenced from each end of the constructs, which contains 20bp from the DNA fragments and 16bp from the linker in the ideal condition.

In the step of linker filtering, local sequence alignment algorithm is used to align the designed linker sequences to the part of linker on real reads sequences. If the alignment score is higher than a user-defined threshold, the linker is identified. Then, the part of linker on the reads sequences is trimmed and the remaining part is kept for further analysis, which should be at least 18bp.

In the original design, the barcodes in the linker sequences are 2bp (AT or CG) corresponding to half-linkers A and B, and now they are changed to 4bp (TAAG or ATGT) [10] in order to improve the differentiation of barcodes (Figure 4). With the two half-linkers A and B (with different barcodes), we can evaluate the random ligation of DNA fragments from different molecular complexes. In the ChIA-PET experiment, the half-linkers are separately added to two aliquots (Figure 1) for ligation with DNA fragments. Then, the two aliquots are mixed for proximal ligation of DNA fragments within the same molecular complexes. During the step of proximal ligation, DNA fragments from two different molecular complexes in the solution could be ligated together by chance. Such process is called random ligation and the ligation products do not reflect real interactions. Since DNA fragments are ligated with two half-linkers in two aliquots separately, the complete linker constructs consisted of AB or BA are definitely from random ligation. With this idea, we can evaluate the random ligation ratio. The real random ligation ratio is roughly double if the samples in two aliquots are in similar amount.

**Figure 4.**
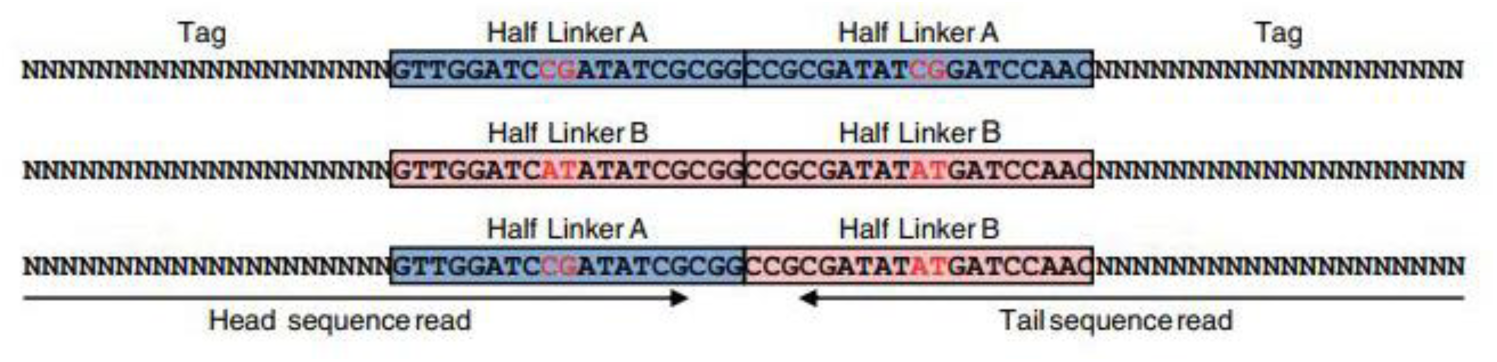
DNA constructs from the experiments in tag-linker-tag format, adapted from Li et al., 2010.

If the linker sequences in the real reads match one of the designed linker sequences with the specified criteria, the PETs will be output to specific FASTQ files with the linker sequences excluded. The main output files from linker filtering program are named as *.m_n.R1.fastq and *.m_n.R2.fastq. For example, the files *.2_1.R1.fastq and *.2_1.R2.fastq are a pair of files with linker B from read1 input file and linker A from read2 input file. The PETs in the files with labels 1_1 or 2_2 are defined as same-linker PETs, and the PETs in the files with labels 1_2 or 2_1 are defined as different-linker PETs.

If we select to process long-read data, the two sequences are from +/- strand of linker without information of barcode. The alignment of linker is similar to that in the short-read process. Especially, we would divide the PETs into different files with the length of paired reads less than or greater than 55bp, so that the accuracy of mapping to the reference genome can be improved. So there would be 8 files named as *.m_n.R1.fastq, *.m_n.R2.fastq, *.m_n.R3.fastq, *.m_n.R4.fastq, *.m_n.R5.fastq, *.m_n.R6.fastq, *.m_n.R7.fastq and *.m_n.R8.fastq in the output files. The R1, R3, R5, R7 are from read1 and R2, R4, R6, R8 are from read2.

In order to speed up the process, ChIA-PET Tool V3 use multithreading in the step of linker filtering. With the Java interface BlockingQueue, one thread reads thousands of PETs from two FASTQ files as a data block and puts it into the queue. And then multiple threads get data blocks from the queue and process them (Figure 5).

**Figure 5.**
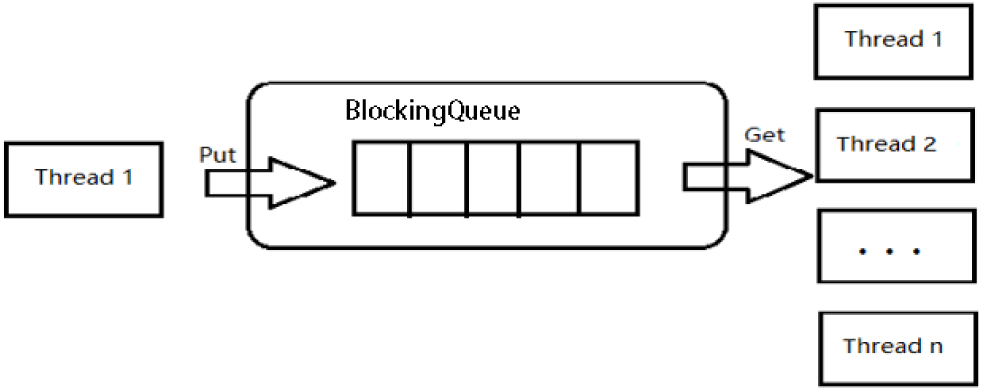
Flowchart of multithreading.

#### 2.2.2. Mapping to genome (Timing 128.75 mins)

--GENOME_INDEX: specifies the path of BWA index file.

--MAPPING_CUTOFF: The mapping threshold to remove the PETs with poor quality. Default: 30.

After linker filtering, the trimmed paired DNA fragments are mapped to a reference genome. A Burrows-Wheeler-transform-based method BatMis [11] was used to generate customized output in SAM format. We assume that the genome index is already built for BWA and the path and the prefix of the index files are specified with variable “GENOME_INDEX”.

For short-read data, the DNA sequences after linker filtering are around 20-21bp, option “aln” in BWA is used for reads mapping and option “samse” is used to convert the mapping results into SAM format. We found that using option “samse” instead of “sampe” could reduce the running time. So we merge the same-linker PETs from pair end. After mapping, we use a Java program to extract the uniquely mapped reads with high mapping quality score and generate BEDPE file with multithreading. The default cutoff for mapping quality score is 30.

For long-read data, the DNA sequences after linker filtering have different lengths, option “aln” are used for PETs length less than 55bp while option “mem” are used for PETs length more than 55bp. After extracting the uniquely mapped reads, SAMtools is used to convert the files from SAM format to BAM format, and BEDTools is used to convert the files from BAM format to BED format.

#### 2.2.3. Cleaning the mapped PETs (Timing 19.65 mins)

--MERGE_DISTANCE: specifies the distance limit to merge the PETs with similar mapping locations. Default: 2.

Different sources of noises exist in ChIA-PET data, including duplicated reads from PCR amplification, variable cutting length from MmeI enzyme, sequencing errors due to the repetitive sequences in the genome, which are considered separately in the step of data purification. Firstly, the reads that are exactly mapped to the same location (Figure A1A) are likely the PCR duplicates, and only one of them is kept for further processing. Secondly, if different PETs have both tags within 2bp at each side (Figure A1B), there are high chances that they are from the same DNA fragment with variable MmeI cutting length or linker filtering cutting. Such PETs are combined as one PET for further processing.

This step consists of two main operations to remove duplicate PETs from amplification and other noises: (1) Merge all PETs with the same mapping locations, probably due to PCR amplification, into one unique PET; (2) Merge all the similar PETs (within ±2bp at the both ends of different PETs) into one unique PET.

#### 2.2.4. Categorization of the PETs (Timing 0.17 mins)

--SELF_LIGATION_CUFOFF: specifies the distance threshold between self-ligation PETs and intra-chromosomal inter-ligation PETs. Default: 8000.

By the ChIA-PET experiment design, the two tags in each paired-end read could come from a single DNA fragment or two different DNA fragments by ligation. In order to identify the chromatin interactions, we should use the PETs from different DNA fragments. We divide the PETs into different categories (Figure 6, adapted from Li et al., 2010): self-ligation PETs (PETs from the single DNA fragments), intra-chromosomal inter-ligation PETs (PETs from two different DNA fragments in the same chromosome), inter-chromosomal inter-ligation PETs (PETs from DNA fragments in two different chromosomes) and others.

**Figure 6.**
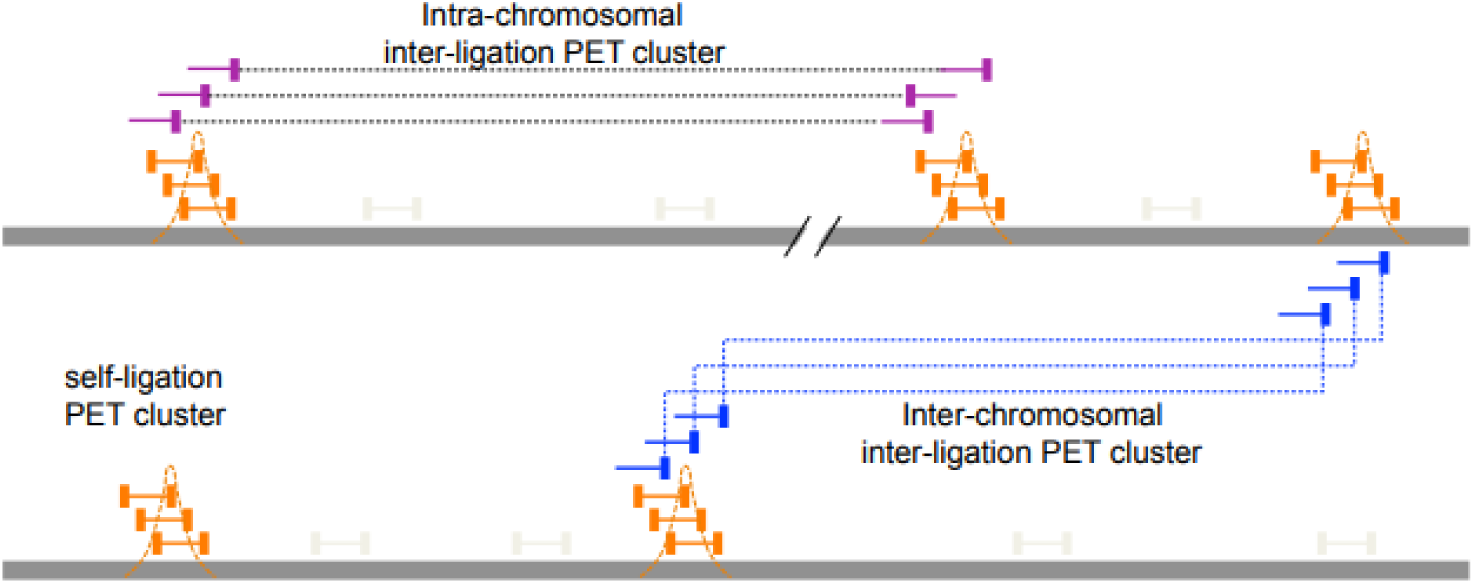
Illustration of the different categories of PETs, adapted from Li et al., 2010.

To separate the self-ligation PETs from intra-chromosomal inter-ligation PETs, we need to determine the genomic distance cutoff between the two tags of the PETs in the same chromosome. By the experiment design and sequencing protocol, the self-ligation PETs are (1) from minus-plus (-/+) chromosome strand composition, (2) with short genomic span and (3) the tag from minus strand should have smaller genomic coordinate. The intra-chromosomal inter-ligation PETs are from all possible strand compositions. By comparing the genomic span distributions for PETs with different strand compositions, it will give a clue of the span threshold for self-ligations. Figure 7 shows that the genomic spans from +/+, +/-, and -/- strand compositions have the similar distributions, while genomic spans from -/+ strand composition have different distributions——there are much more PETs with genomic spans less than 10Kb from -/+ strand composition. Based on the assumption that the intra-chromosomal inter-ligation PETs should be uniformly distributed among different strand compositions, the extra PETs with genomic span less than 10Kb from -/+ strand composition are from self-ligation. Then, the difference of genomic span distributions from -/+ strand composition and the average distribution from other strand compositions is used to determine the span cutoff for self-ligation. The log-log plot of genomic span distribution difference in Figure 8 shows that the cutoff is around 8Kb. For the illustration, we use 8Kb as the default cutoff to classify the PETs into different categories.

**Figure 7.**
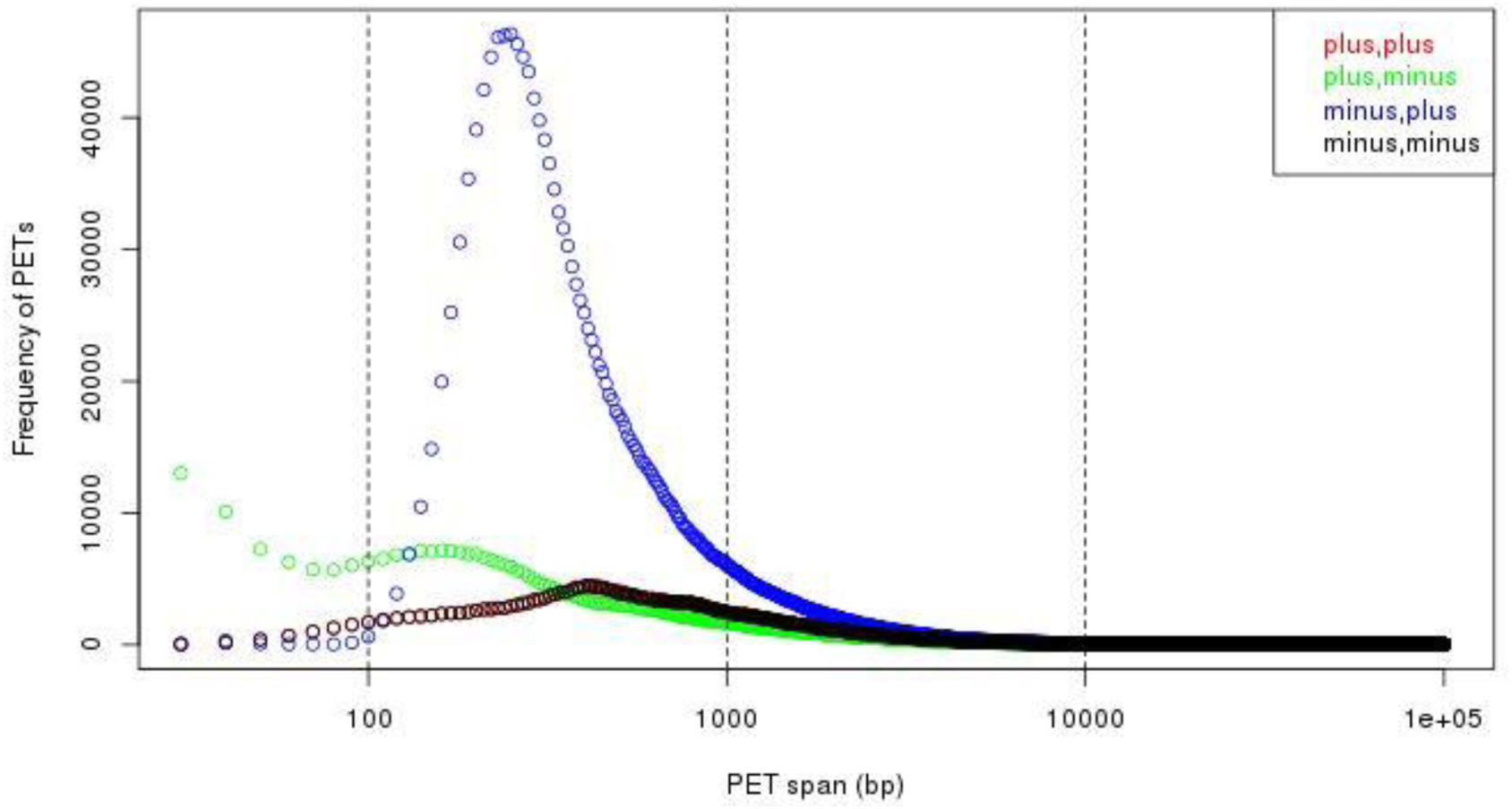
Genomic span distributions from different strand compositions. We can see that there are much more PETs with minus-plus (-/+) strand composition in short span (less than 10Kb, especially less than 1Kb).

**Figure 8.**
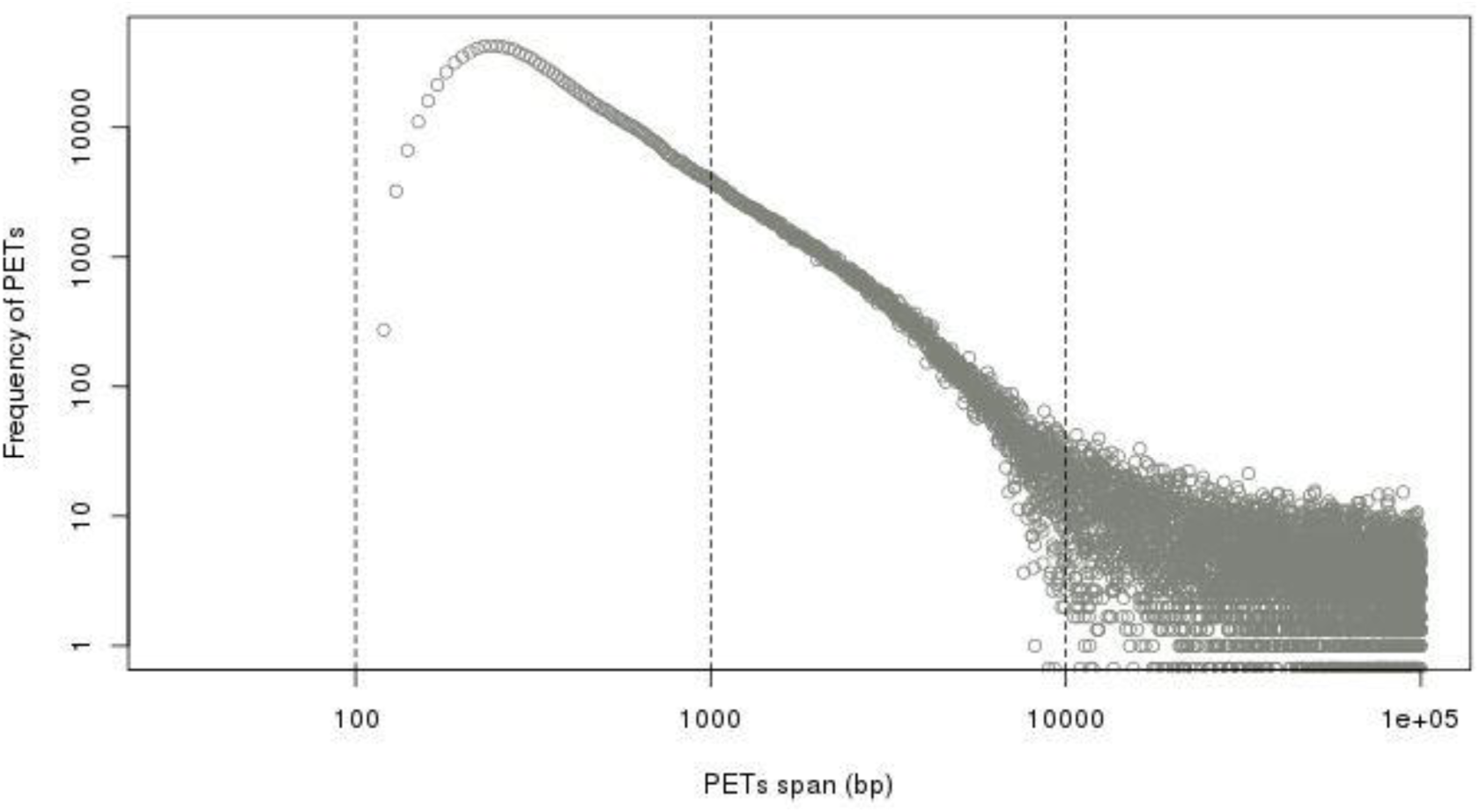
Difference of genomic span distributions from minus-plus (-/+) strand composition and average distribution from other strand compositions in log-log plot. It shows that the self-ligation cutoff is around 8Kb.

#### 2.2.5. Peak calling (Timing 0.50 mins)

--EXTENSION_LENGTH: specifies the extension length from the location of each tag, which is determined by the median span of the self-ligation PETs. Default: 500.

--MIN_COVERAGE_FOR_PEAK: specifies the minimum coverage to define peak regions. Default: 5.

--PEAK_MODE: There are two modes for peak calling. Number 1 is “peak region” mode, which takes all the overlapping PET regions above the coverage threshold as peak regions, and number 2 is “peak summit” mode, which takes the highest coverage of overlapping regions as peak regions. Default: 2.

--MIN_DISTANCE_BETWEEN_PEAK: specifies the minimum distance between two peaks. If the distance of two peak regions is below the threshold, only the one with higher coverage will be kept. Default: 500.

--GENOME_LENGTH: specifies the number of base pairs in the whole genome.

--GENOME_COVERAGE_RATIO: specifies the estimated proportion of the genome covered by the reads. Default: 0.8.

--PVALUE_CUTOFF_PEAK: specifies p-value to filter peaks that are not statistically significant. Default: 0.00001.

Peak calling with self-ligation PETs are similar to the peak calling from ChIP-Seq data with paired-end sequencing mode. The enrichment of the PETs in a genomic region is considered as the potential binding sites of the protein of interest. Different ChIP-Seq peak calling methods, such as MACS [12], can be used. In ChIA-PET Tool V3, a method similar to MACS is provided for peak calling. Overlapping regions of self-ligation PETs are used to define transcription factor binding sites and Poisson distribution is used to calculate the p-values for the peaks.

#### 2.2.6. Interaction calling (Timing 4.35 mins)

--CHROM_SIZE_INFO: specifies the file that contains the length of each chromosome. INPUT_ANCHOR_FILE: a file which contains user-specified anchors for interaction calling. If you don’t have this file, please specify the value of this variable as “null” instead. Default: null.

--PVALUE_CUTOFF_INTERACTION: specifies p-value to filter false positive interactions. Default: 0.05.

Overlapping regions of inter-ligation PETs are used to define interacting regions and hyper-geometric distribution is used to calculate p-value for the interactions. Chromatin interaction calling is based on the overlapping of the extended tags from different PETs. The extension of the tags is based on the fact that the sequencing reads are just part of DNA fragments in the experiment. The tag extension length is determined by the median span of the self-ligation PETs, which is around 500bp. For illustration, we use 500bp as the default parameter. In the current pipeline, the interaction calling does not depend on the given peaks, since some interactions may not be supported by strong peaks at the anchor regions. This is why we changed the interaction calling to the overlap of the extended tags from the PETs. Still, we provide the option for users to call chromatin interactions with any given regions, which can be from ChIP-Seq peak calling, or any genomic regions of interest. The statistical significance of the interaction is assessed by p-value from hyper-geometric model and adjusted for multiple hypothesis testing.

#### 2.2.7. Visualization Report

--CYTOBAND_DATA: specifies the ideogram data used to plot intra-chromosomal peaks and interactions.

--SPECIES: specifies the genome used to plot inter-chromosomal interactions, 1 for human, 2 for mouse and 3 for others. Default: 1.

The results of ChIA-PET data analysis are visualized in two ways: (1) the statistics of the data quality; (2) the list of peaks and interactions. The statistics along the data processing procedure is summarized in a HTML file for users to check the information of the data. The chromosome interaction maps contain interaction information of intra-chromosomal and inter-chromosomal. Maybe the interaction information can not be clearly observed which is affected by the amount of chromosomes and interactions. We use Discuz to achieve the image zooming (Figure 9). Discuz is an image processing toolkit based JavaScript. When you click on a picture, the picture will be displayed in a new window, which can be dragged and zoomed by mouse. The ChIA-PET data (original mapping PETs, peaks and interactions) are converted into BED format for visualization with a genomic browser, such as UCSC browser.

**Figure 9.**
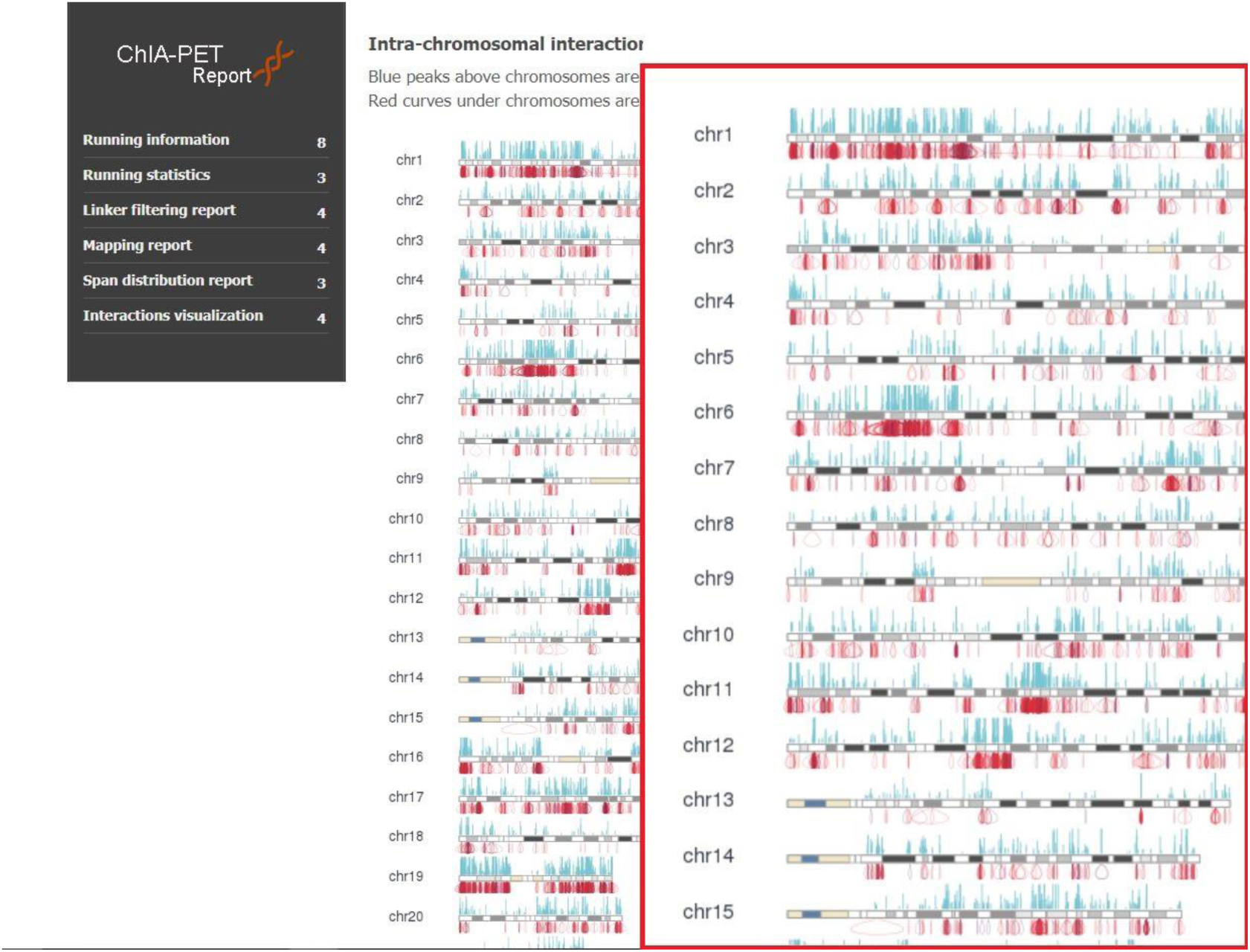
Realization of picture zooming

## 3. Results

### 3.1. Anticipated results

ChIA-PET Tool V3 can process the next-generation sequence data from ChIA-PET experiment to generate enriched binding peaks of the protein of interest and the related chromatin interactions. We demonstrated the application of ChIA-PET Tool V3 with RNAPII-associated ChIA-PET data from human K562 cells as an example.

The results from linker filtering include a few summary statistics. Figure A2A shows the distribution of linker alignment scores. It reflects the distribution of best alignment scores from the designed linker sequences to the reads. We can see that most of the alignment scores are 10, which means that most of the linker sequences in the reads are 10 base pairs. Figure A2B shows the distribution of linker alignment score differences. It reflects the distribution of alignment score differences between the best-aligned linker and the second-best aligned linker. This distribution is used to check how different between the best-aligned linker and the second-best aligned linker, in case there are ambiguities to locate the linker in the reads. Figure A2C shows the distribution of tag lengths. It reflects the distribution of tag lengths after trimming the best-aligned linker sequences from the reads. We can see that most of the tag lengths are 20 or 21bp, which is consistent with enzyme MmeI’s digestion property. Table 1 reports the proportion of each linker combination in the reads. The results show that most (90.73%) of the PETs are composed of same-linker (A_A or B_B). This indicates proper proximity-ligation within individual molecules.

**Table 1.**
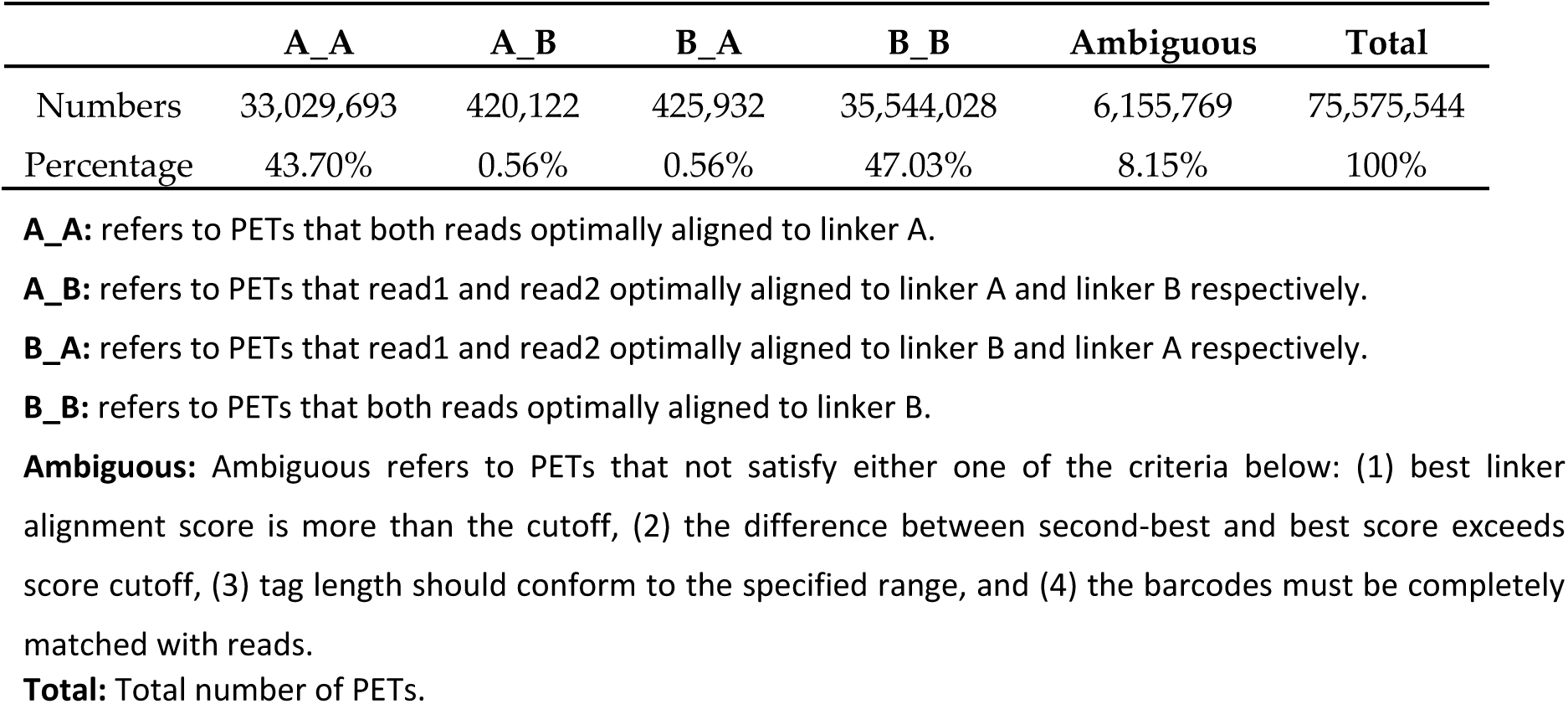
Statistics of linker composition

|  | A_A | A_B | B_A | B_B | Ambiguous | Total |
| --- | --- | --- | --- | --- | --- | --- |
| Numbers | 33,029,693 | 420,122 | 425,932 | 35,544,028 | 6,155,769 | 75,575,544 |
| Percentage | 43.70% | 0.56% | 0.56% | 47.03% | 8.15% | 100% |

**A\_A:** refers to PETs that both reads optimally aligned to linker A.
**A\_B:** refers to PETs that read1 and read2 optimally aligned to linker A and linker B respectively.
**B\_A:** refers to PETs that read1 and read2 optimally aligned to linker B and linker A respectively.
**B\_B:** refers to PETs that both reads optimally aligned to linker B.
**Ambiguous:** Ambiguous refers to PETs that not satisfy either one of the criteria below: (1) best linker alignment score is more than the cutoff, (2) the difference between second-best and best score exceeds score cutoff, (3) tag length should conform to the specified range, and (4) the barcodes must be completely matched with reads.
**Total:** Total number of PETs.

Figure 10 shows a screenshot of binding profile and chromatin interactions from the example data set. Peaks from self-ligation PETs of RNAPII-associated ChIA-PET are mainly enriched around gene promoter regions. The interactions are mainly between gene promoter regions and other regulatory elements, which indicate that RNAPII-associated chromatin interactions are involved in gene transcription regulation. The first 6 lines of the peak output file from Step 5-peak calling are in Table A1. The first 6 lines of the interaction output file from Step 6-interaction calling are in the following Table A2. The statistics of the clusters are summarized in Table 2 and Table 3. We can see that majority of the clusters are intra-chromosomal interactions. Majority of the interactions are spans within 1Mb. Figure A3 shows the chromosome view of binding peaks and intra-chromosomal chromatin interactions. The binding peaks are distributed all over the whole genome. Most of the intra-chromosomal chromatin interactions are within a short span (within 1Mb) and a small portion of chromatin interactions can span a very long distance. Figure A4 shows the circular view of the inter-chromosomal chromatin interactions. Compared to intra-chromosomal chromatin interactions, there are fewer inter-chromosomal chromatin interactions.

**Table 2.**
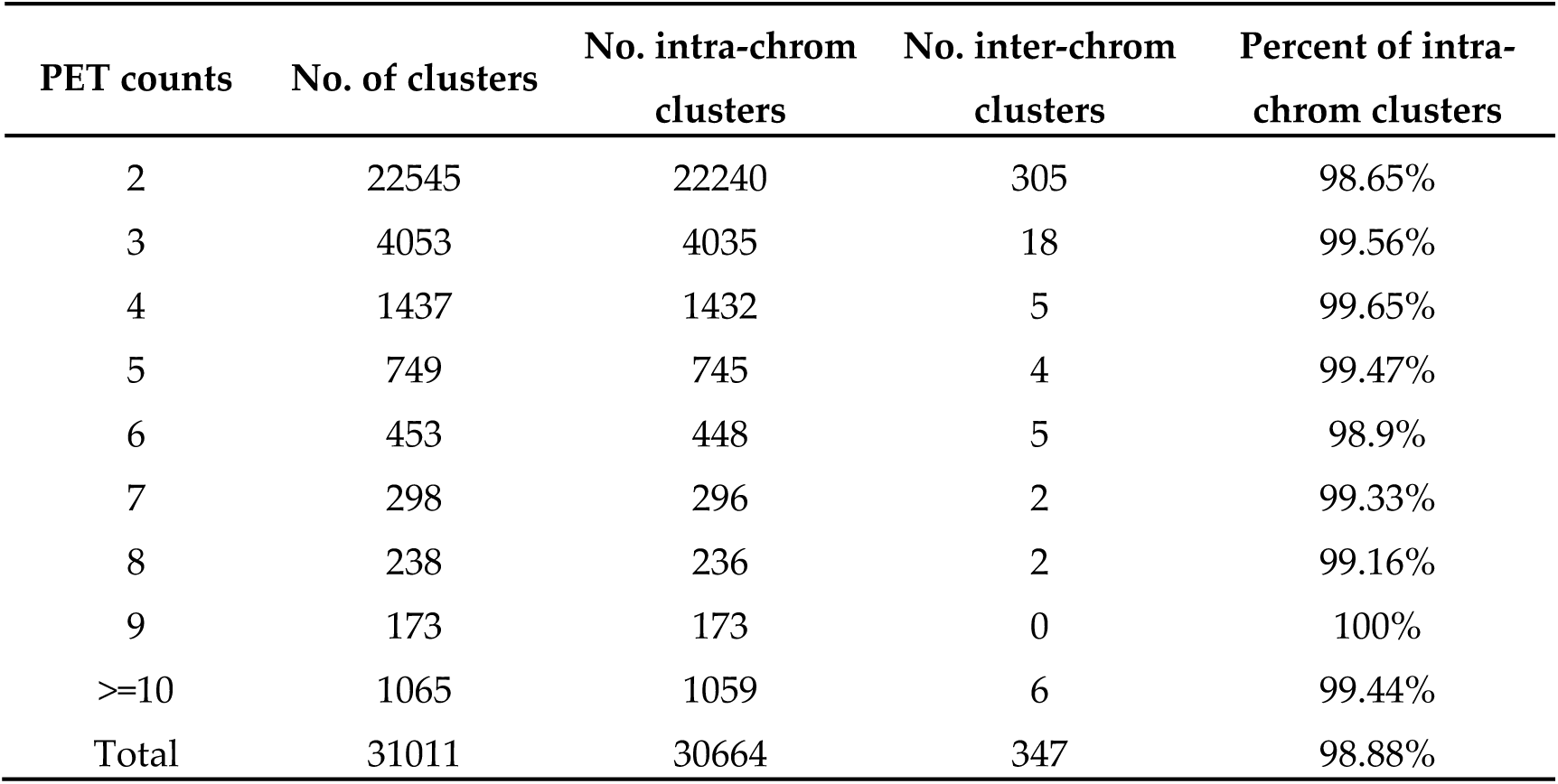
Statistics of interactions with PET counts

**Table 3.**
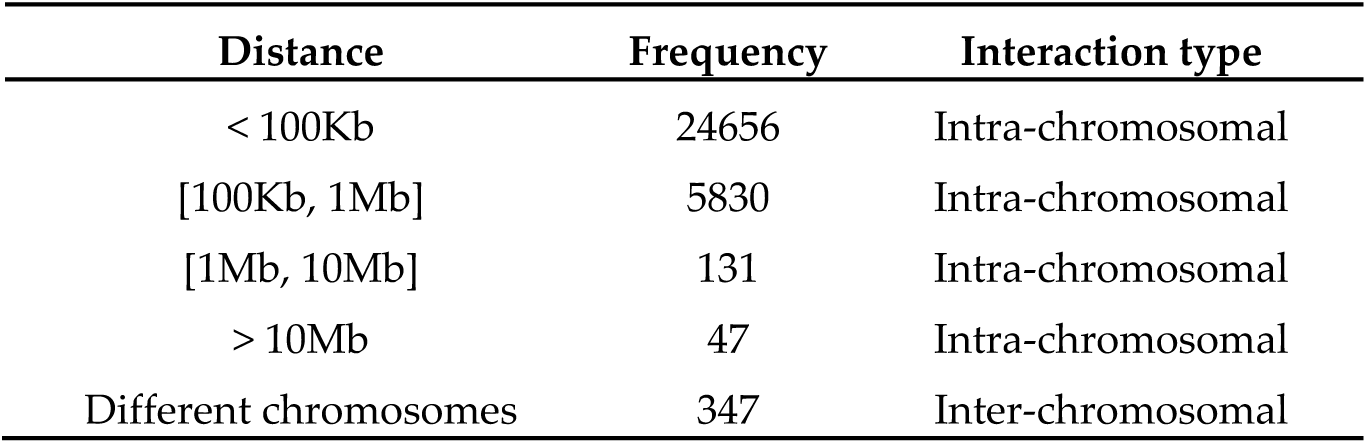
Span distribution of interactions

**Figure 10.**
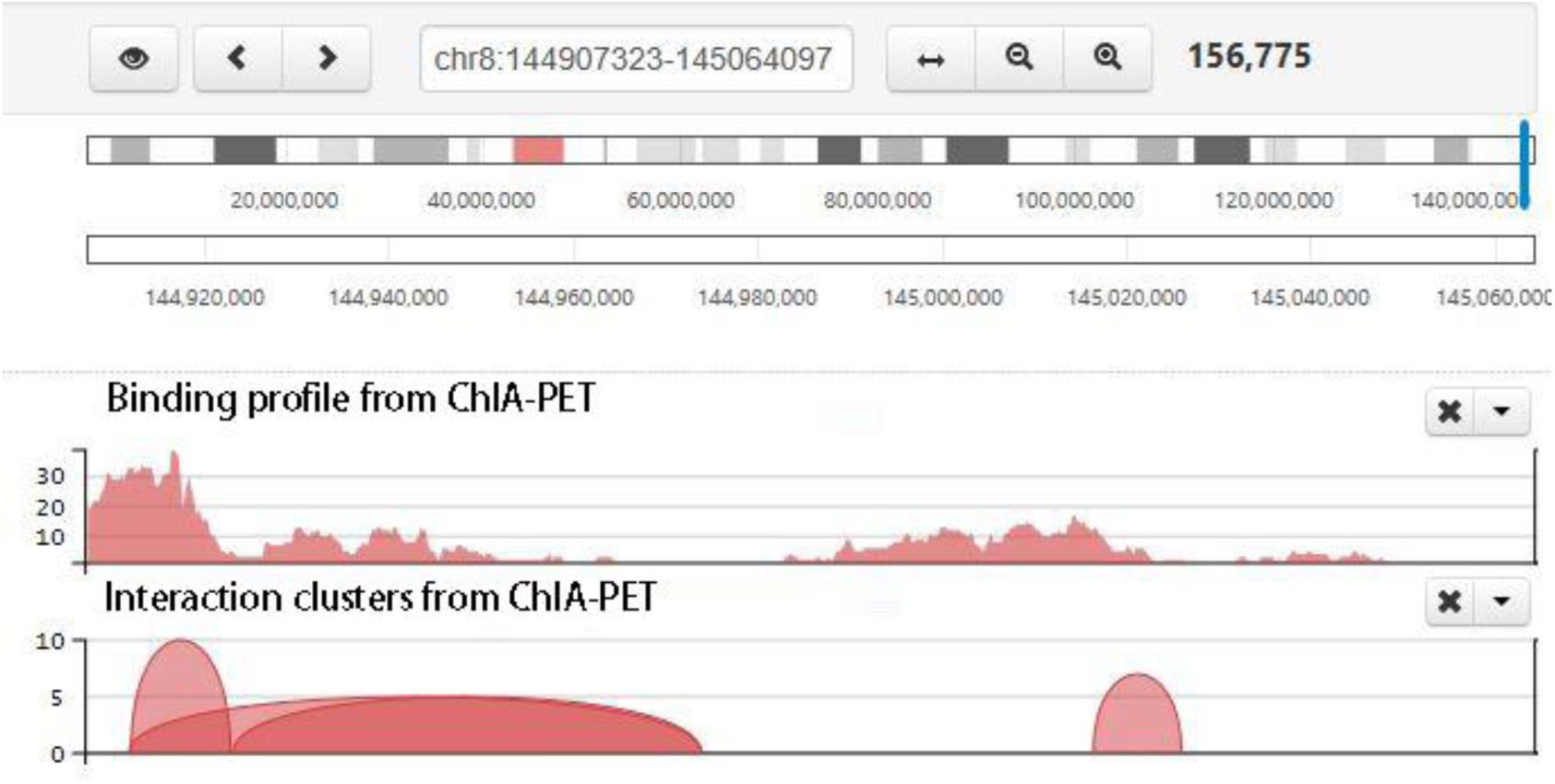
Screenshot of chromatin clusters and protein binding peaks from ChIA-PET. There are multiple tracks, including inter-ligation PETs, interaction clusters and binding profile from ChIA-PET. In the interaction cluster track, two ends of each curve are the interaction anchors, and the height of the curve is the pet count between the interaction anchors. The higher the curve, the stronger the interaction.

### 3.2. Evaluation of the data quality

One concern for the users is the quality of the ChIA-PET libraries they generated or used. For evaluation of the data, we provide the summary statistics of result and the statistics of ChIA-PET libraries.

#### 3.2.1. Result of short-read ChIA-PET data

Table 4 shows the statistics of result after finishing the data analysis. Table 5 shows the statistics of ChIA-PET libraries. The 1st line in Table 7 is the percentage of the same-linker PETs over the total PETs. By the nature of the ChIA-PET design, there will be more same-linker PETs than different-linker PETs. In general, we see the percentage of the same-linker PETs varying from 60% to 99%. If such percentage is above 75%, the library is good in the linker composition level. The second line is the percentage of the uniquely mappable PETs over the total PETs, which varies with the libraries. The 3rd line is the percentage of the PETs after merging those mapped to the same positions exactly over the uniquely mappable PETs. If this percentage is too low (less than 30%), it means that there are more PETs from PCR amplification and the data is already near saturated. Then it does not worth to re-sequence this library for more distinct PETs. If the PETs after merging those mapped to the same positions exactly are 70% or more of the uniquely mappable PETs, deeper sequencing can be applied to get more data for this library. The 4th line is the percentage of inter-ligation PETs over all the PETs after purification. This is about the efficiency catching the interacting PETs. If this percentage is low, it means that the library has too few inter-ligation PETs and it is not good enough for chromatin interaction detection. The 5th line is the percentage of intra-chromosomal inter-ligation PETs over the total inter-ligation PETs. From the current understanding, most of the chromatin interactions are within the individual chromosomes. Therefore, there should be more intra-chromosomal PETs than inter-chromosomal PETs. If there are more inter-chromosomal PETs, it means that the proximity ligation introduces many random ligations. The 6th line is the peak number. This depends on the transcription factor used, and should be compared with the background knowledge or available ChIP-Seq data. For RNAPII and CTCF, there are tens of thousands of peaks in a good ChIA-PET library from human and mouse. The 7th line is the interaction number. This depends on the transcription factors used. For RNAPII and CTCF, there are tens of thousands of interactions in a good ChIA-PET library from human and mouse.

**Table 4.**
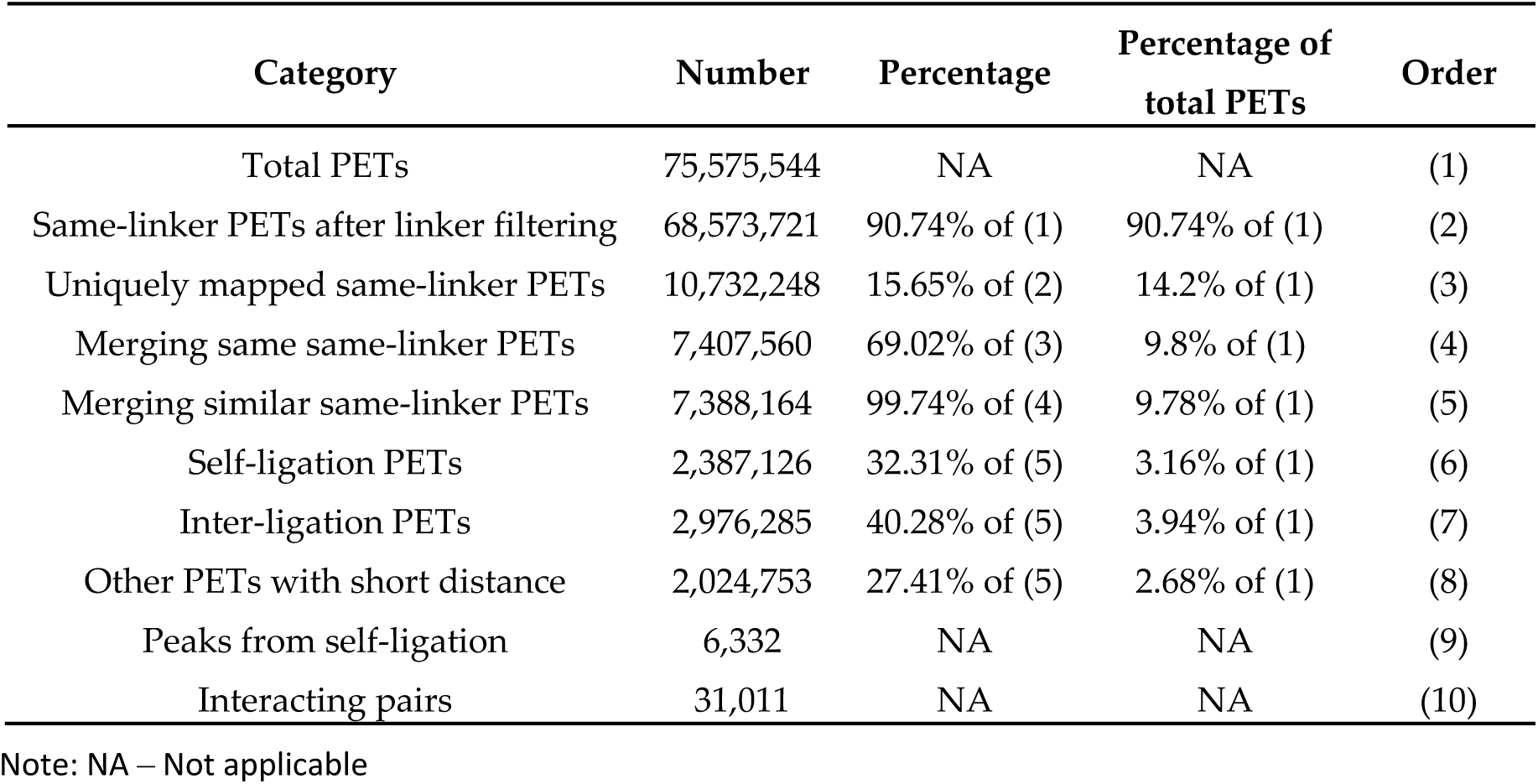
Statistics of RNAP II from human K562 cells

**Table 5.**
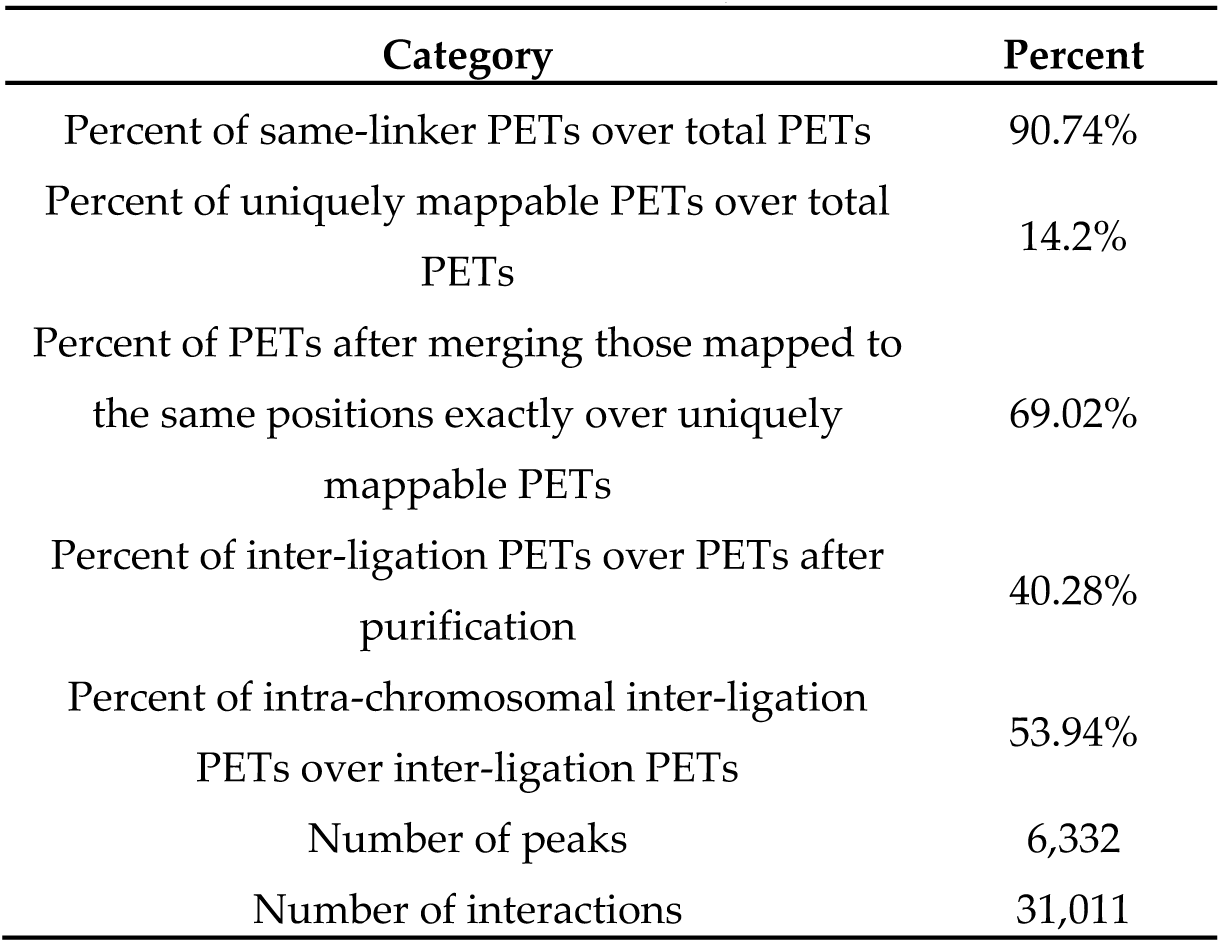
Statistics of a ChIA-PET library from human K562 cells

| Category | Percent |
| --- | --- |
| Percent of same-linker PETs over total PETs | 90.74% |
| Percent of uniquely mappable PETs over total PETs | 14.2% |
| Percent of PETs after merging those mapped to the same positions exactly over uniquely mappable PETs | 69.02% |
| Percent of inter-ligation PETs over PETs after purification | 40.28% |
| Percent of intra-chromosomal inter-ligation PETs over inter-ligation PETs | 53.94% |
| Number of peaks | 6,332 |
| Number of interactions | 31,011 |

**Table 6.**
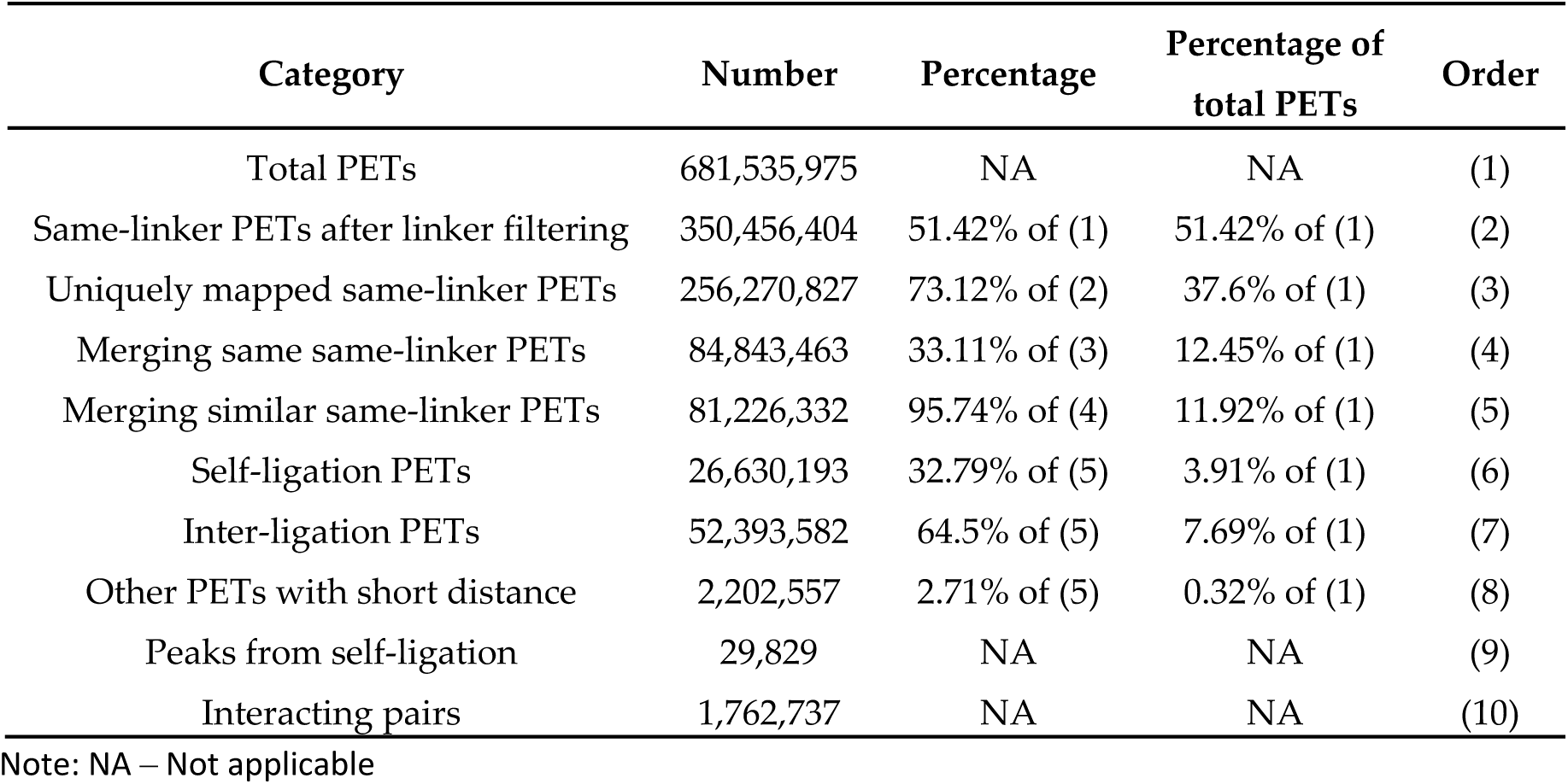
Statistics of CTCF from human GM12878 cells

**Table 7.** Statistics of a ChIA-PET library from human GM12878 cells

| Category | Percent |
| --- | --- |
| Percent of same-linker PETs over total PETs | 51.42% |
| Percent of uniquely mappable PETs over total PETs | 37.6% |
| Percent of PETs after merging those mapped to the same positions exactly over uniquely mappable PETs | 33.11% |
| Percent of inter-ligation PETs over PETs after purification | 64.5% |
| Percent of intra-chromosomal inter-ligation PETs over inter-ligation PETs | 30.49% |
| Number of peaks | 29,829 |
| Number of interactions | 1,762,737 |

#### 3.2.2. Result of long-read ChIA-PET data

The statistical indicators of result in the long-read analysis result are similar to those in short-read analysis result (Table 6 and Table 7).

### 3.3 Comparing results with other tools

#### 3.3.1 Comparing the results on short-read data

With the developing of ChIA-PET, there are more and more published tools such as, Mango [13], ChIA-PET2 [14] and ChiaSig [15] and so on, to process and analyze ChIA-PET data in recent years. In order to evaluate ChIA-PET Tool V3, we use ChIA-PET data associated with RNAP II from human K562 cells as input data and compare the results of ChIA-PET Tool V3 with other tools (Table 8). In the step of mapping PETs to a reference genome, Mango use Bowtie while ChIA-PET Tool V3 and ChIA-PET2 use BWA. Meanwhile, the standard of mapping quality score is different. So we use BWA and select 30 as the cutoff of quality score in the step of mapping instead of Bowtie when we execute Mango. Otherwise, ChiaSig only performs the step of significant loop calling. For the significant interactions in the table, there are at least 3 supportive PETs in one interaction. And the value of false discovery rate (FDR) is 0.05.

**Table 8.** Statistics of different tools from human K562 cells

| Tools | non-chimeric<br>PETs | non-duplicate<br>PETs | peaks | significant<br>interactions |
| --- | --- | --- | --- | --- |
| Mango | 67,330,362 | 7,286,203 | 16,589 | 469 |
| ChIA-PET2 | 67,347,877 | 7,303,170 | 17,121 | 5,179 |
| ChIA-PET Tool V3 | 68,573,721 | 7,388,164 | 6,332 | 8,466 |
| ChiaSig | NA | NA | NA | 7,370 |

Note: NA – Not applicable

In the results, Mango and ChIA-PET2 can detect more peaks while ChIA-PET Tool V3 and ChIA-PET2 can detect more significant interactions. The overlaps of peaks and significant interactions detected by different tools are showing as Figure 11. For peaks, more than 94% peaks of ChIA-PET Tool V3 overlap with Mango and ChIA-PET2. Scatter plots (Figure 12) and box plots (Figure 13) represent the peak intensity between every two different tools. The Pearson correlation coefficients in scatter plots between the three tools are above 0.9. Especially, the Pearson correlation coefficient is 0.99 between Mango and ChIA-PET2. It’s because both of them use MACS to call peak. For ChIA-PET Tool V3, the difference of common and unique parts in the box plots is not significant.

**Figure 11.**
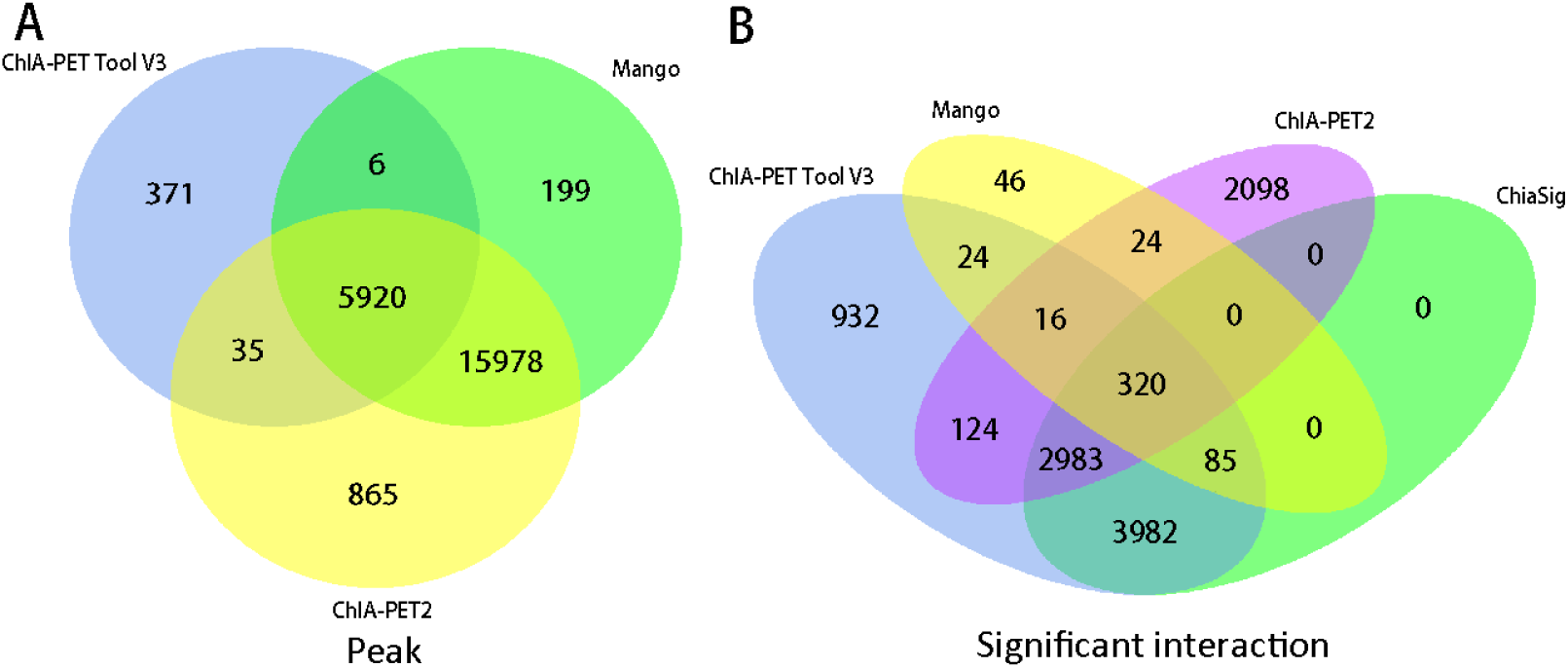
Overlap of peaks and interactions detected by different tools from human K562 cells.

**Figure 12.**
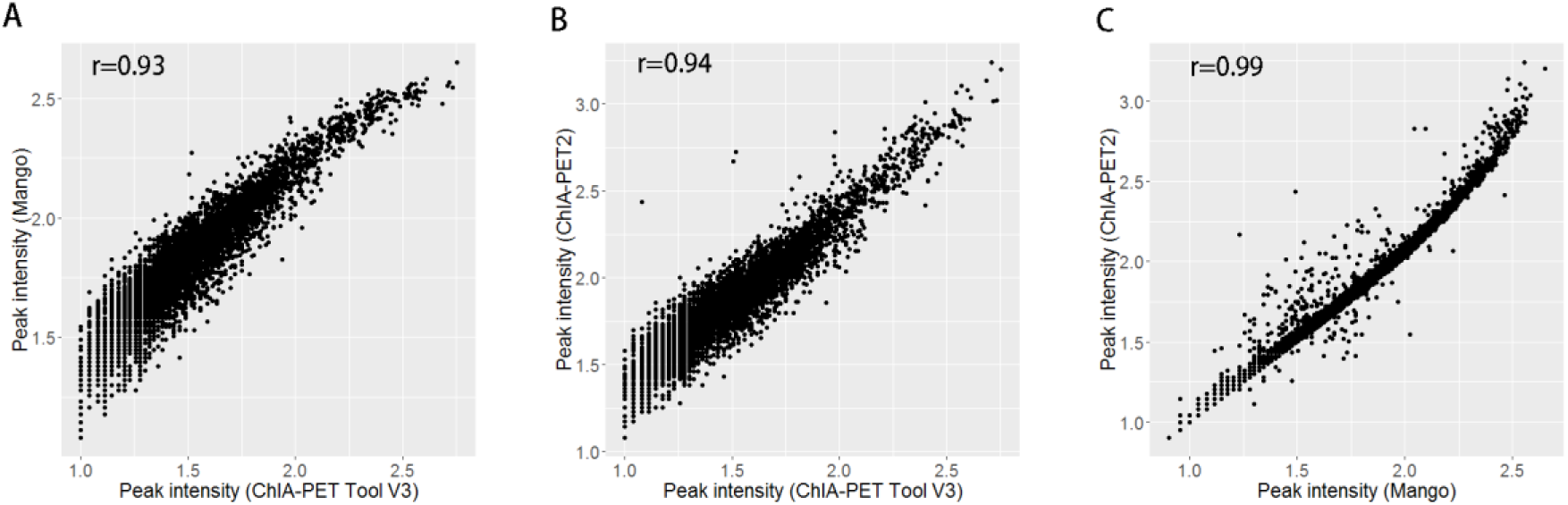
Scatter plots of peak intensity between different tools from human K562 cells.

**Figure 13.**
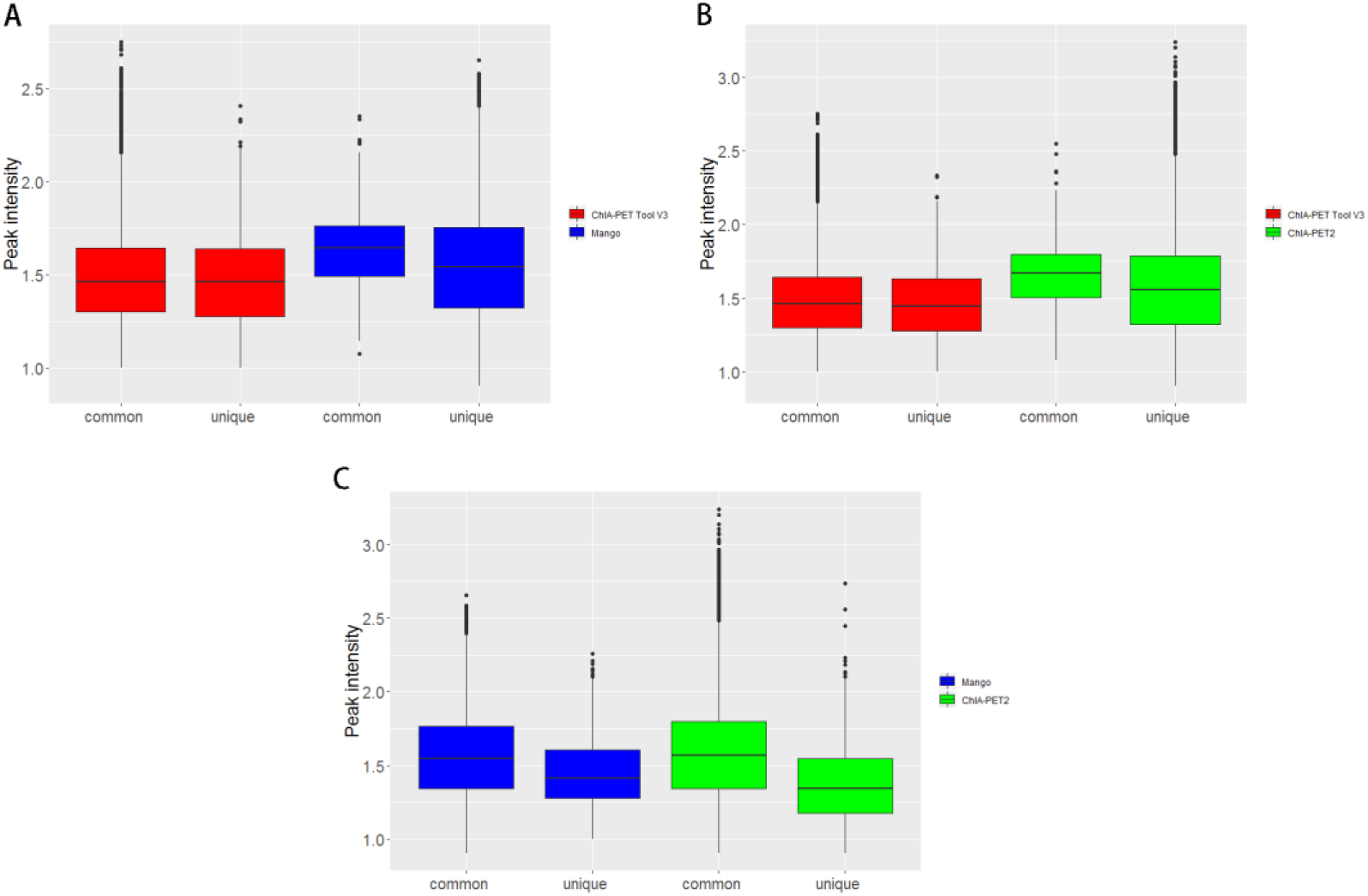
Box plots of peak intensity between different tools from human K562 cells. The common part represents peak intensity without overlap between two tools. And the unique part represents peak intensity without overlap between two tools. Middle line denotes median; box denotes IQR; whiskers denote 1.5× IQR.

For significant interactions, most of interactions of Mango and ChiaSig overlap with ChIA-PET Tool V3 except ChIA-PET2. 932 interactions (blue area) of ChIA-PET Tool V3 have no overlap with other tools. Among them, 558 interactions include peaks while 374 interactions don’t include peaks. However, all of 2,098 interactions (purple area) of ChIA-PET2 which have no overlap with other tools include peaks. Scatter plots (Figure 14) and box plots (Figure 15) represent the interaction intensity between every two different tools. The Pearson correlation coefficient between ChIA-PET Tool V3 and ChiaSig is higher as well as between Mango and ChIA-PET2. The difference of common and unique parts in the box plots is significant except ChIA-PET Tool V3 and ChiaSig.

**Figure 14.**
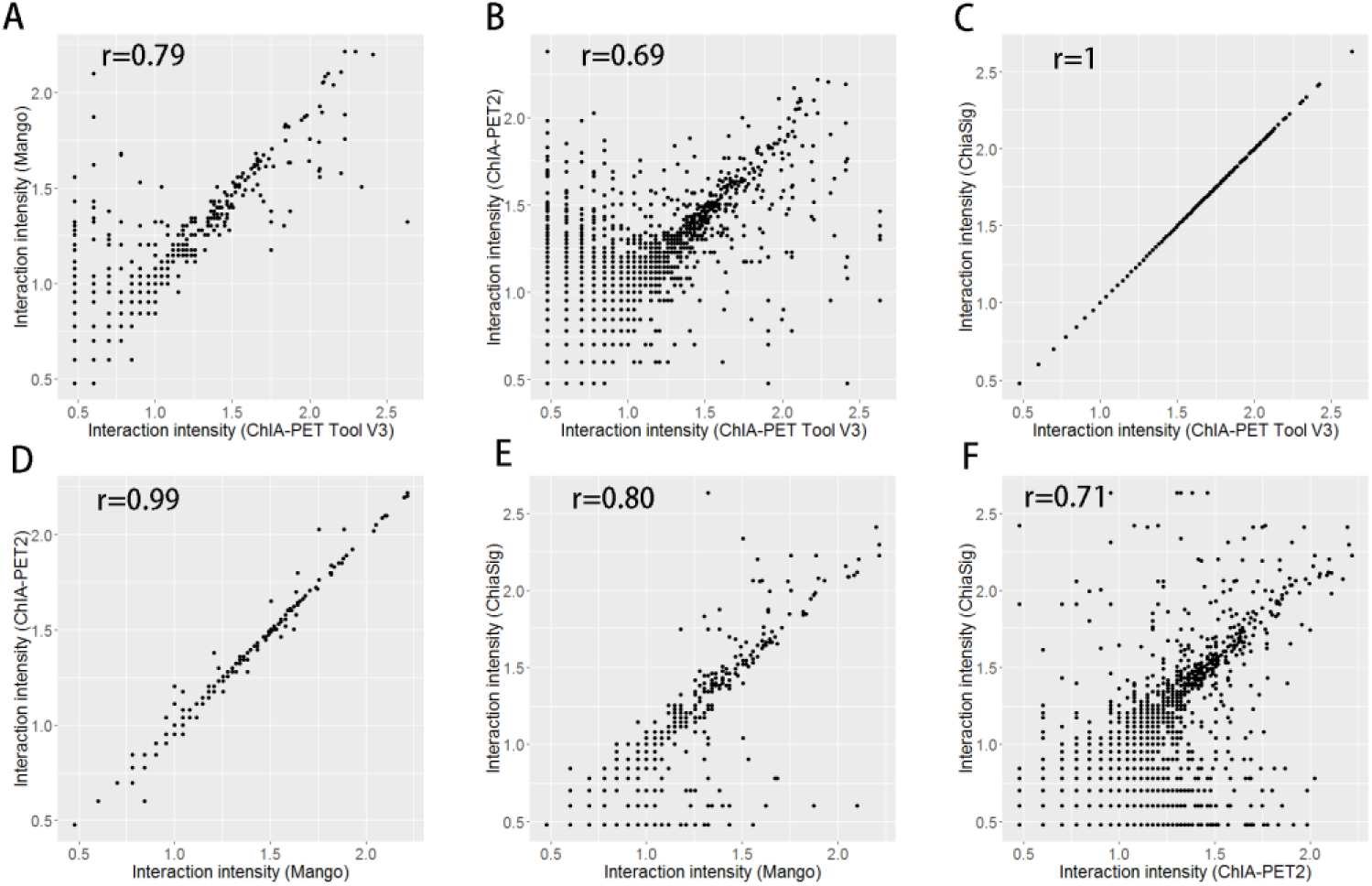
Scatter plots of interaction intensity between different tools from human K562 cells.

**Figure 15.**
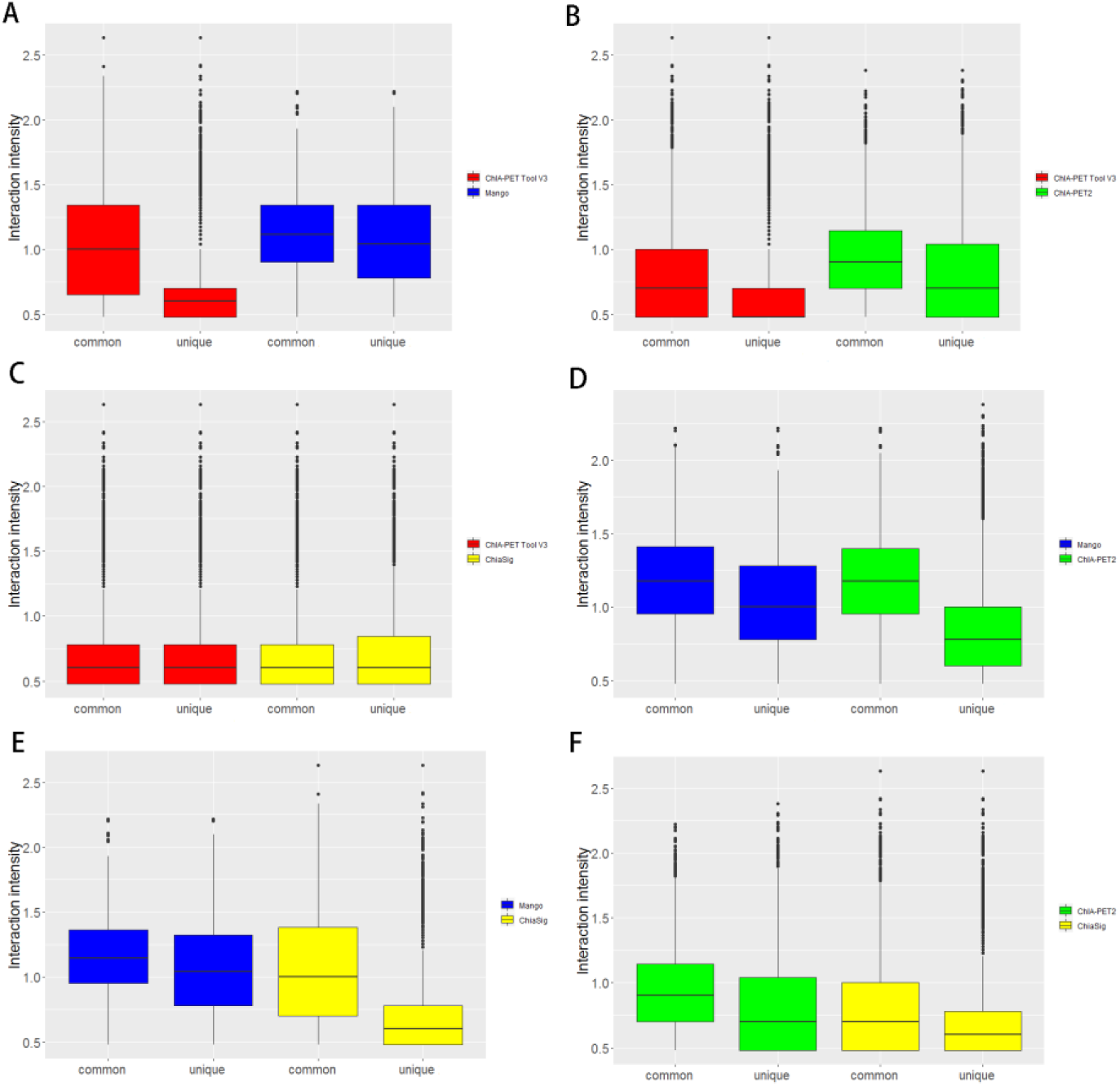
Box plots of interaction intensity between different tools from human K562 cells.

We calculated the distribution of PET counts in significant interaction (Figure A5). Most interactions have only 1 PET in ChIA-PET2 while most interactions have only 2 PETs in ChIA-PET Tool V3. We generated aggregate peak analysis (APA) plots (Figure 16) with no more than 3 PETs for the significant interactions detected by different tools. To generate APA plots, we summed interaction counts for pairs of loci in 25kb bins. The level of different sets of interactions can be quantified by an APA score. Higher scores indicate higher enrichment. In these plots, APA scores for ChIA-PET Tool V3 and ChiaSig are similar and both are higher than Mango and ChIA-PET2.

**Figure 16.**
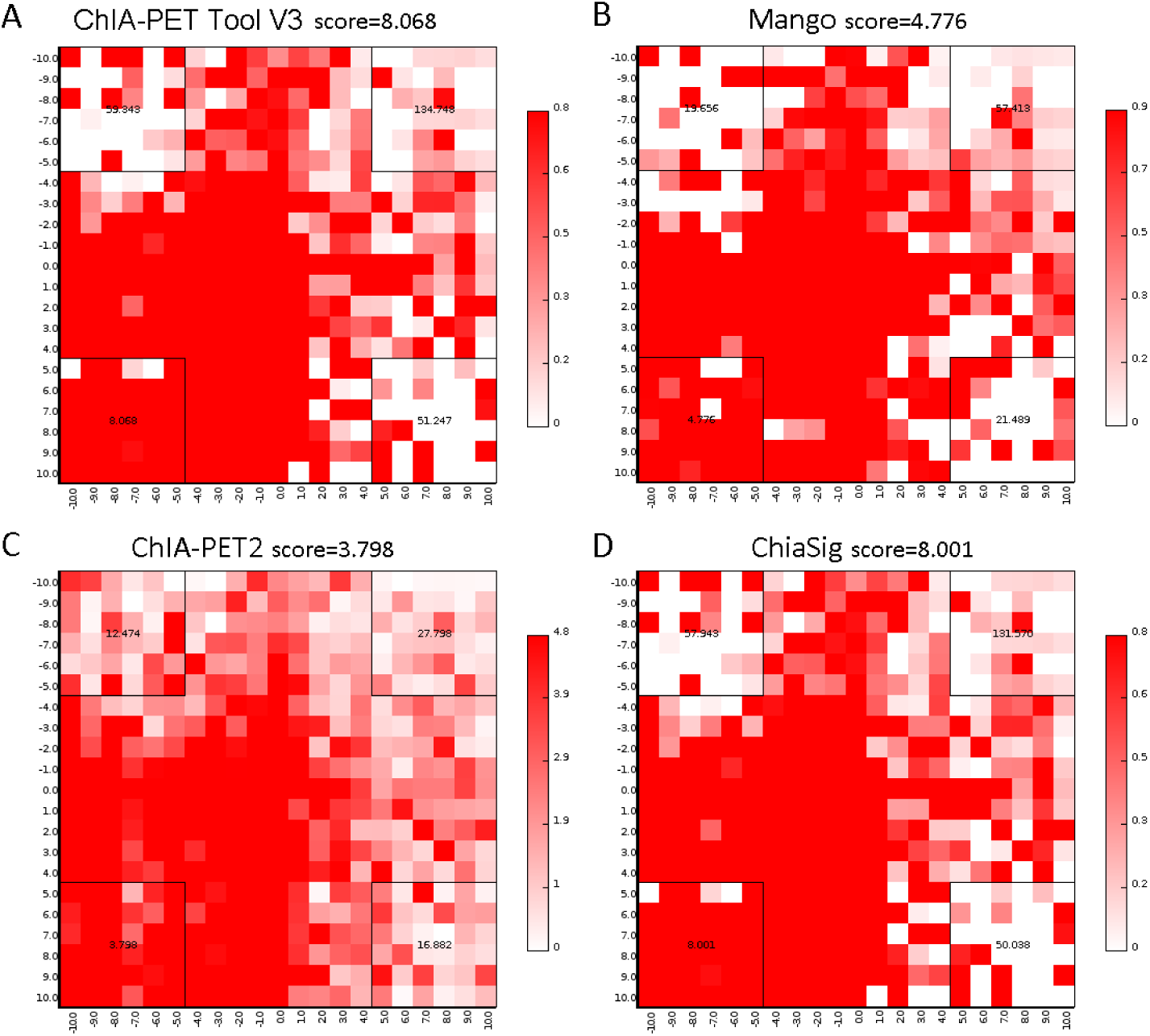
APA plots of significant interactions detected by different tools from human K562 cells. Interaction counts are summed for all pairs of loci in 25-kb bins.

#### 3.3.2 Comparing the results on long-read data

Considering both ChIA-PET Tool V3 and ChIA-PET2 can process and analyze long-read ChIA-PET data, we executed ChIA-PET Tool V3 and ChIA-PET2 using ChIA-PET data CTCF from human GM12878 cells as input. In Table 9, one interaction has at least 3 PETs. And the value of FDR is 0.05. We compared the results and generated overlap plots of peaks and significant interactions detected by ChIA-PET Tool V3 (Figure 17A) and ChIA-PET2 (Figure 17B). More than 94% peaks of ChIA-PET Tool V3 overlap with ChIA-PET2. In scatter plot (Figure 17C), the Pearson correlation coefficient between ChIA-PET Tool V3 and ChIA-PET2 is 0.94. And the difference of common and unique parts in the box plot (Figure 17D) is significant.

**Table 9.** Statistics of different tools from human GM12878 cells

| Tools | non-chimeric<br>PETs | non-duplicate<br>PETs | peaks | significant<br>interactions |
| --- | --- | --- | --- | --- |
| ChIA-PET Tool V3 | 350,456,404 | 81,226,332 | 29,829 | 420,494 |
| ChIA-PET2 | 106,975,449 | 32,278,242 | 47,725 | 37,313 |

**Figure 17.**
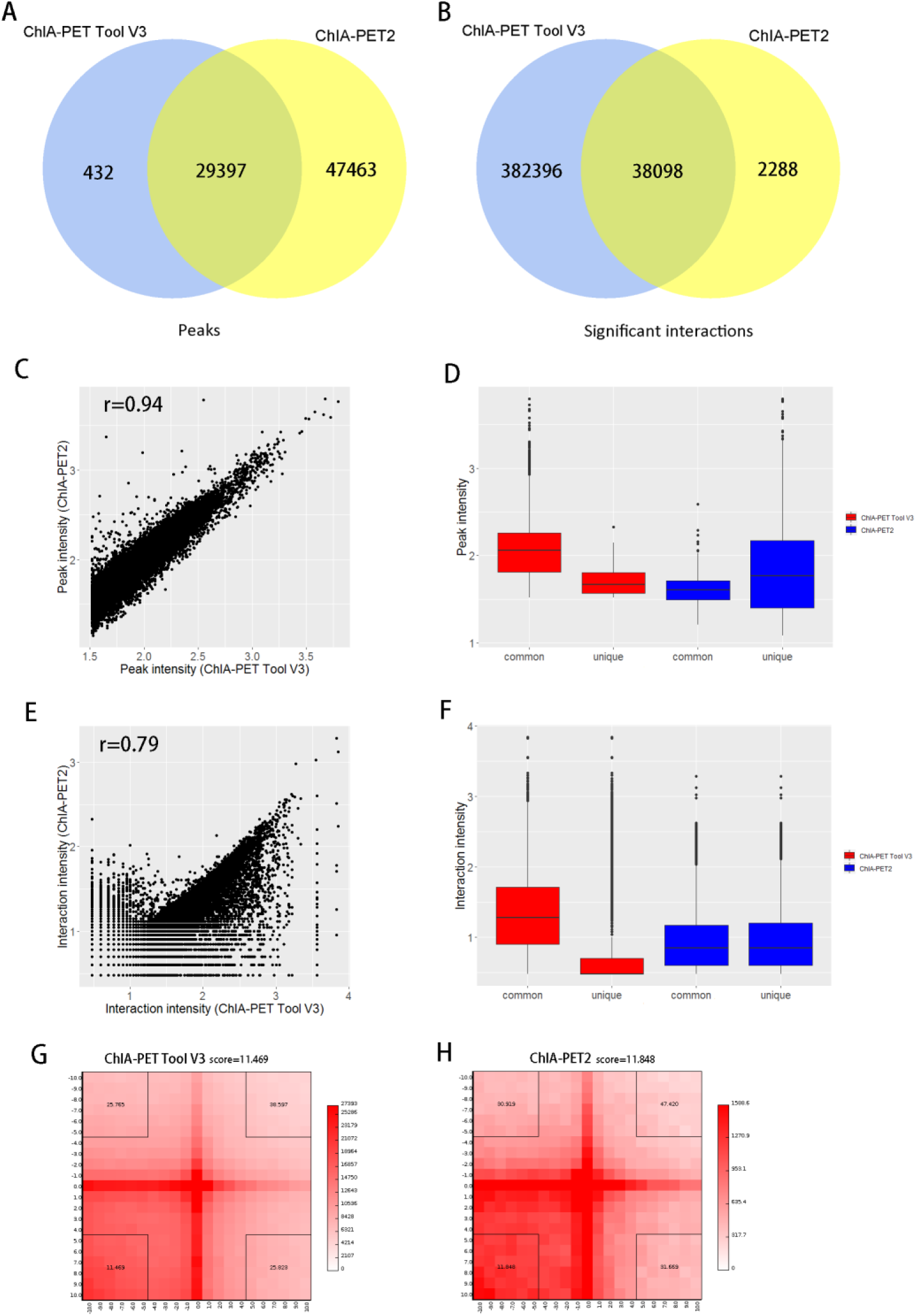
Results of comparison on long-read data. (**a**) and (**b**) Overlap of peaks and interactions detected by different tools from human GM12878 cells. (**c**) and (**d**) Scatter plot and box plot of peak intensity from human GM12878 cells. (**e**) and (**f**) Scatter plot and box plot of interaction intensity from human GM12878 cells. (**g**) and (**h**) APA plots of significant interactions from human GM12878 cells. Interaction counts are summed for all pairs of loci in 25-kb bins.

Most of interactions of ChIA-PET Tool V3 overlap with ChIA-PET2. 382,396 interactions of ChIA-PET Tool V3 have no overlap with other tools. Among them, 177,001 interactions include peaks while 205,395 interactions don’t include peaks. However, all of 2,288 interactions of ChIA-PET2 which have no overlap with other tools include peaks. In scatter plot (Figure 17E), the Pearson correlation coefficient between ChIA-PET Tool V3 and ChIA-PET2 is 0.94. And the difference of common and unique parts in the box plot (Figure 17F) is significant for ChIA-PET Tool V3. Figure A6 is the PET count distribution of significant interactions. We generate APA plots with no more than 3 PETs as Figure 17G and Figure 17H. The APA scores of ChIA-PET Tool V3 and ChIA-PET2 are similar.

## 4. Discussion

In this protocol, we introduce the design and usage of ChIA-PET Tool V3. By processing RNAP II and CTCF ChIA-PET data, we demonstrate how to apply ChIA-PET Tool V3 to the public ChIA-PET data and illustrate the details of the results.

ChIA-PET Tool V3 can process short-read and long-read ChIA-PET data from paired-end reads. It would generate enriched binding peaks and the chromatin interactions associated with a protein of interest. Multiple log files and statistics are generated for tracing the processing issues if any step goes wrong and for evaluating the quality of the libraries. ChIA-PET Tool V3 has the advantages of users-friendly, multithreading and result visualization. During the execution of ChIA-PET Tool V3, the statistics of the library is generated and summarized in a HTML file. It’s clear to check the information of the data in charts. Especially, users can zoom the plots of interactions using mouse wheel to check the interactions in details. With the development of 3D genome technologies, we believe more researchers need tools like ChIA-PET Tool V3 to analyze the chromatin interaction data and obtain more information to understand fundamental and structures of genome.

## Author Contributions

Conceptualization, G.L.; methodology, G.L.; software, G.L., T.S., H.C., and L.C.; writing—original draft preparation, T.S., H.C., and L.C.; writing—review and editing, G.L., H.P. and Q.Z.; supervision, G.L.; funding acquisition, G.L..

## Acknowledgments

We thank the group members for helpful discussions.

### Appendix A

All parameters required in ChIA-PET Tool V3 are as follows. We illustrate them further in the corresponding steps.

#### Necessary options

--mode: There are two modes for ChIA-PET Tool V3. 0 for short read, 1 for long read.

--fastq1: path of read1 fastq file.

--fastq2: path of read2 fastq file.

--linker: linker file.

--minimum_linker_alignment_score: Specifies the allowed minimum alignment score.

--GENOME_INDEX: specifies the path of BWA index file.

--GENOME_LENGTH: specifies the number of base pairs in the whole genome.

--CHROM_SIZE_INFO: specifies the file that contains the length of each chromosome.

--CYTOBAND_DATA: specifies the ideogram data used to plot intra-chromosomal peaks and interactions.

--SPECIES: specifies the genome used to plot inter-chromosomal interactions, 1 for human, 2 for mouse and 3 for others.

#### Optional options

--start_step: start with which step, 1: linker filtering; 2: mapping to genome; 3: removing redundancy; 4: categorization of PETs; 5: peak calling; 6: interaction calling; 7: visualizing, default: 1”.

--output: specifies the directory to store the output data from ChIA-PET Tool V3, default: ChIA-PET_Tool_V3/output.

--prefix: specifies the prefix of all the output files, default: out.

--maximum_tag_length: Specifies the maximum tag length. Default:1000 Specifies the maximum tag length. Default: 1000.

--output_data_with_ambiguous_linker_info: Determines whether to print the linker-ambiguity PETs. 0: not print; 1: print, Default: 1.

--thread: the number of threads used in linker filtering and mapping to genome. Default: 1.

--MAPPING_CUTOFF: The mapping threshold to remove the PETs with poor quality. Default: 30.

--MERGE_DISTANCE: specifies the distance limit to merge the PETs with similar mapping locations.

Default: 2.

--MIN_COVERAGE_FOR_PEAK: specifies the minimum coverage to define peak regions. Default:5.

--GENOME_COVERAGE_RATIO: specifies the estimated proportion of the genome covered by the reads.

Default: 0.8.

--INPUT_ANCHOR_FILE: a file which contains user-specified anchors for interaction calling. If you don’t have this file, please specify the value of this variable as “null” instead. Default: null.

--PVALUE_CUTOFF_INTERACTION: specifies p-value to filter false positive interactions. Default:0.05.

**Table A1.** Example of peak file

| chrom | summit start | summit end | peak coverage | p-value | p.adjust |
| --- | --- | --- | --- | --- | --- |
| chr1 | 840305 | 840556 | 21 | 1.18e-08 | 2.57e-07 |
| chr1 | 876391 | 876492 | 40 | 1.07e-07 | 1.99e-06 |
| chr1 | 877684 | 877775 | 44 | 6.46e-10 | 1.75e-08 |
| chr1 | 935728 | 935873 | 63 | 8.41e-11 | 2.58e-09 |
| chr1 | 936421 | 936425 | 64 | 1.67e-11 | 5.64e-10 |
| chr1 | 937164 | 937208 | 55 | 3.1e-07 | 5.25e-06 |

**chrom:** chromosome name.
**summit start:** The start coordinate of peak summit.
**summit end:** The end coordinate of peak summit.
**peak coverage:** The highest coverage by tags in a peak.
**p-value:** This value represents the statistical significance of a peak, which is calculated by Poisson distribution.
**p.adjust:** P.adjust means p-value adjusted with Benjamini-Hockberg method for multiple hypothesis testing.

**Table A2.**
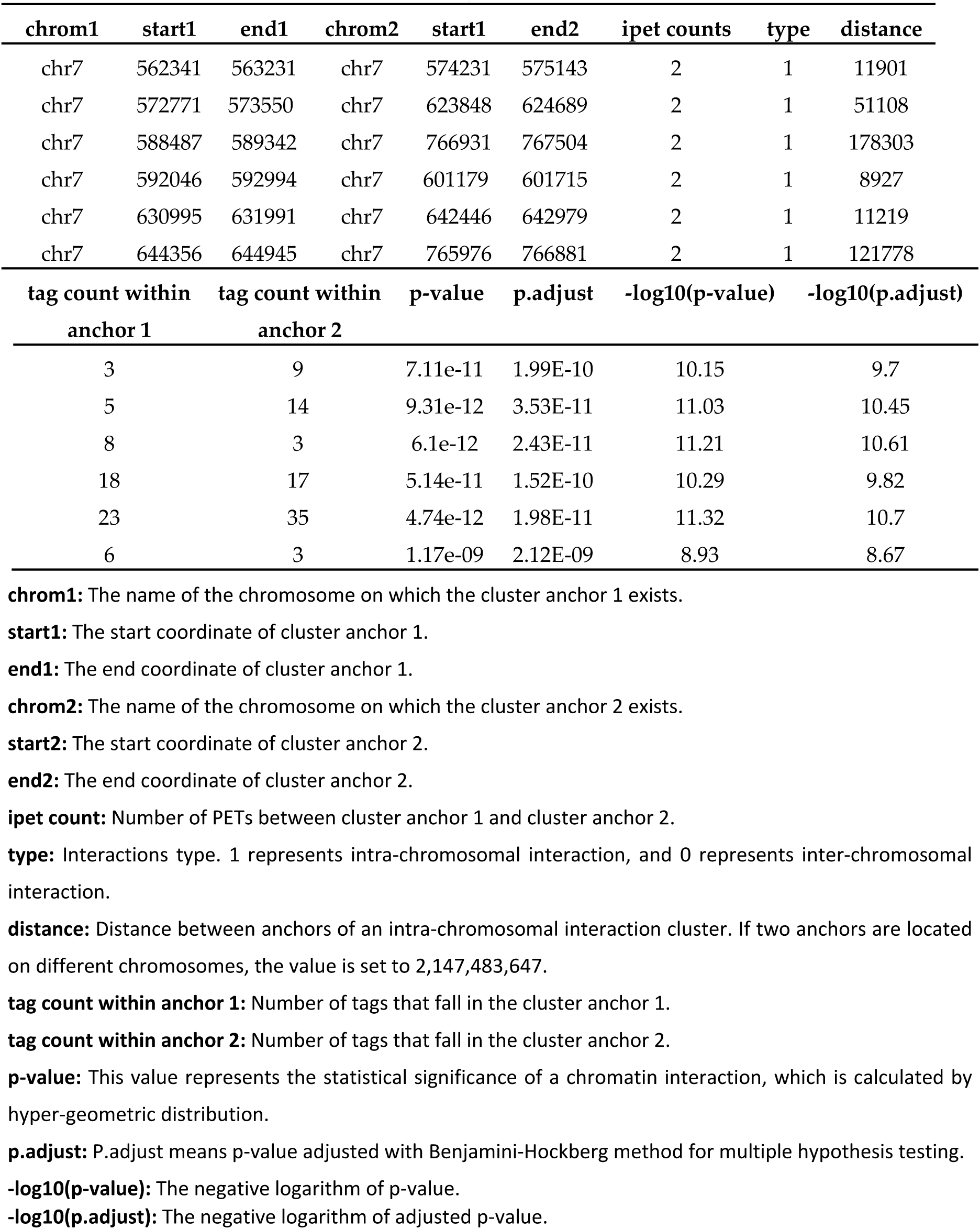
Example of interaction file

**Figure A1.**
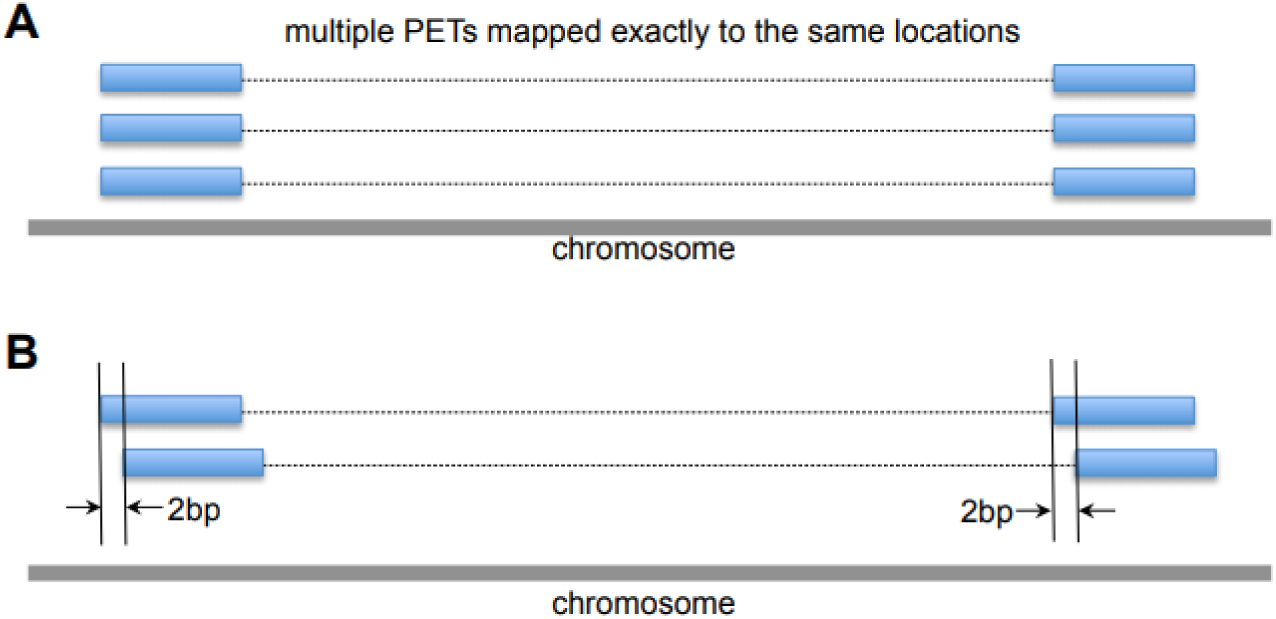
Examples of noises in the ChIA-PET data. (**a**) Duplicates of PETs from PCR amplification which are mapped exactly to the same locations. (**b**) Different PETs with tags within 2bp at both ends.

**Figure A2.**
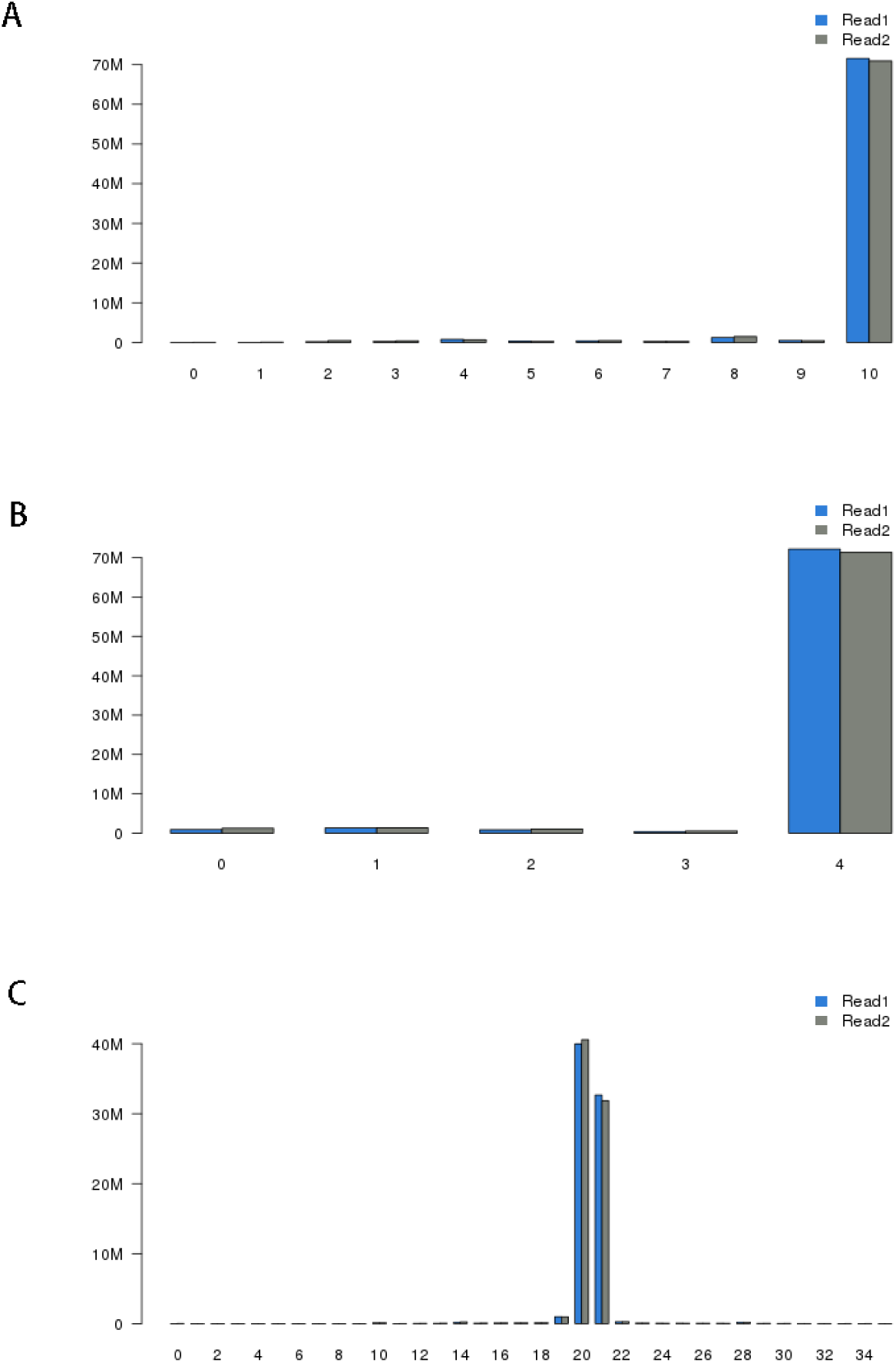
Statistics from linker filtering. (**a**) Distribution of linker alignment scores. Most of the linker alignment scores are 10, which are expected by the design. (**b**) Distribution of linker alignment score differences. (**c**) Distribution of tag lengths. Most of tags after trimming the linker sequences are 20bp or 21bp long, which is expected from enzyme Mme I’s digestion property.

**Figure A3.**
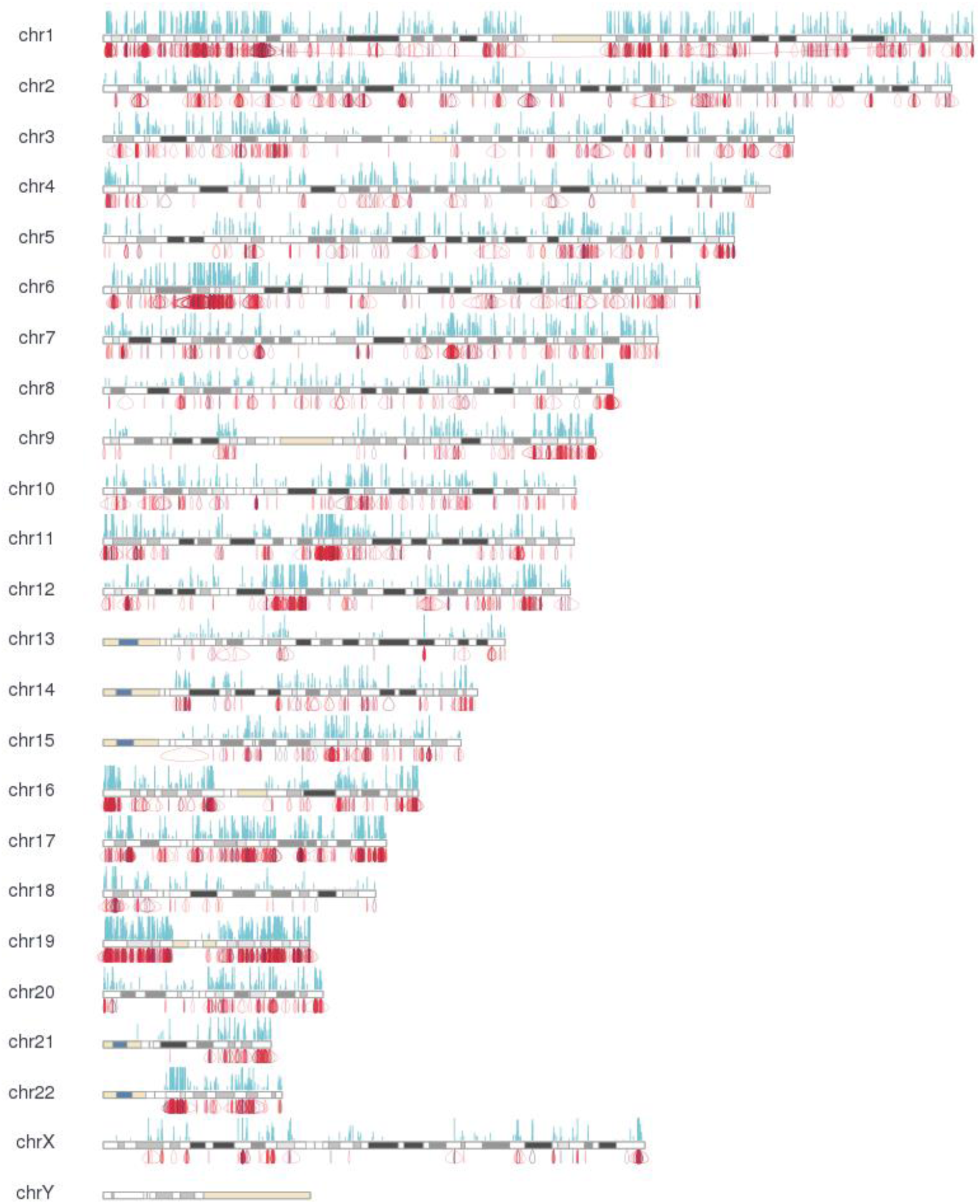
Chromosome view of binding peaks and intra-chromosomal chromatin interactions from human cell line K562. The chromosomes are shown in cytoband format. The blue bars above the chromosomes are the binding peaks, and the height of the peaks are the coverage of the self-ligation PETs over the peaks. The red curves under the chromosomes are the interaction clusters.

**Figure A4.**
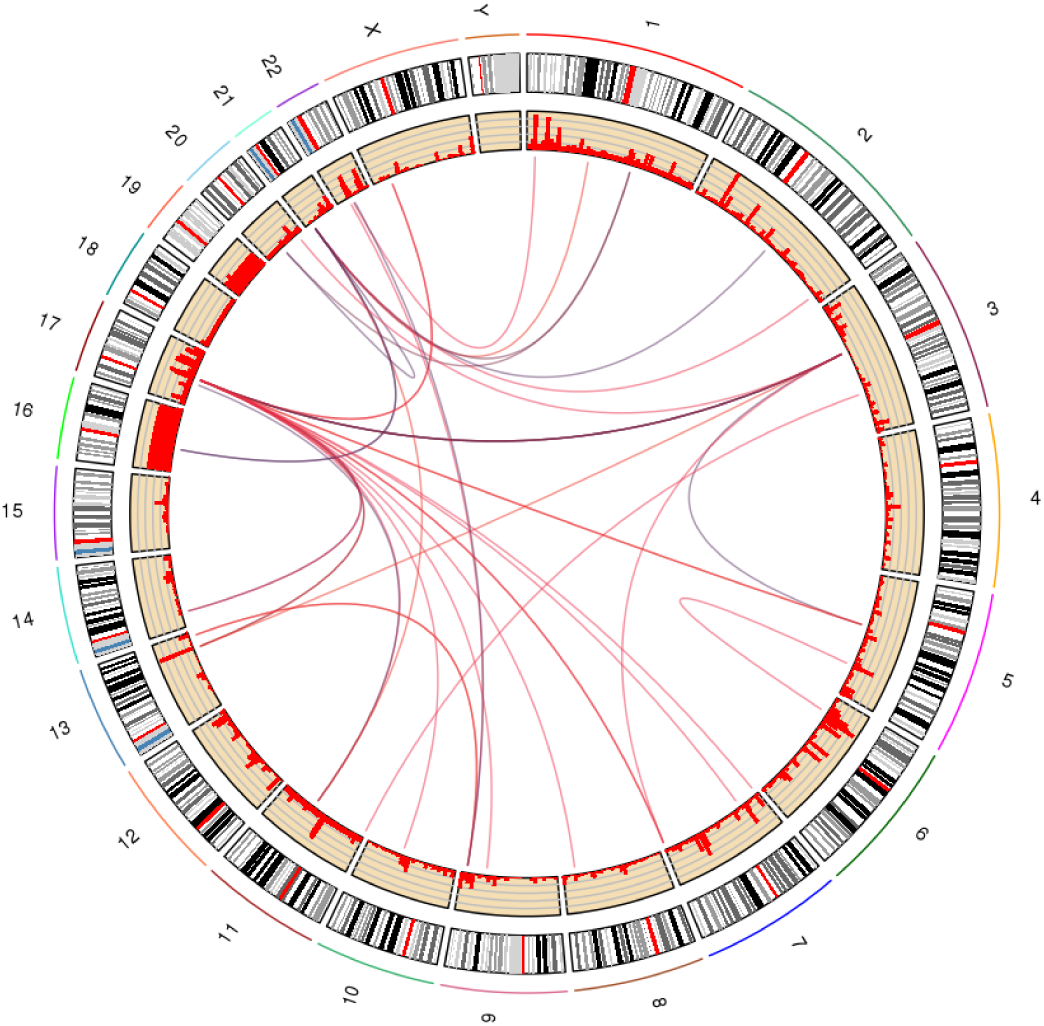
Circular view of chromatin interactions between different chromosomes from human cell line K562. Each curve represents one interaction.

**Figure A5.**
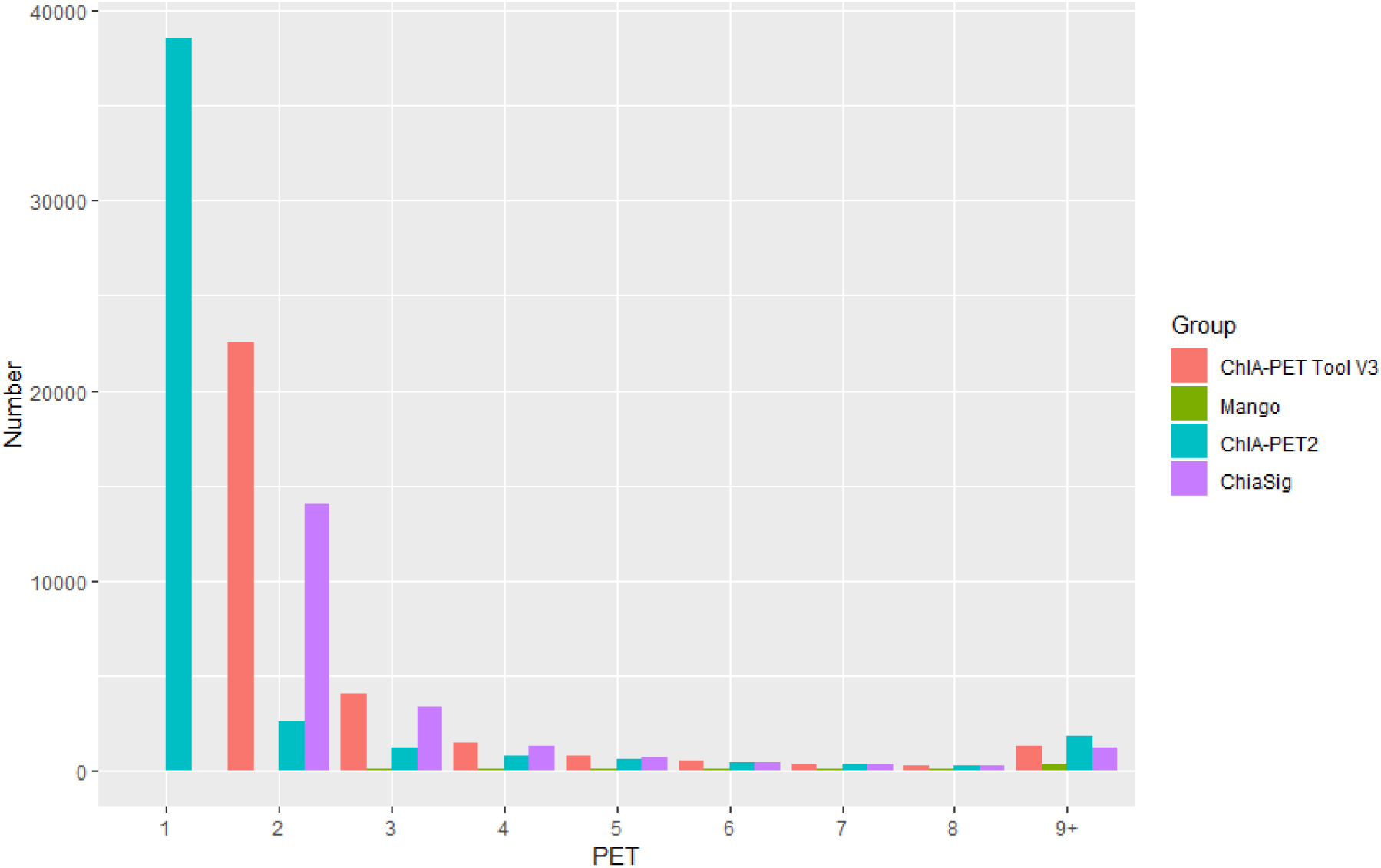
PET count distribution of significant interactions detected by different tools from human K562 cells.

**Figure A6.**
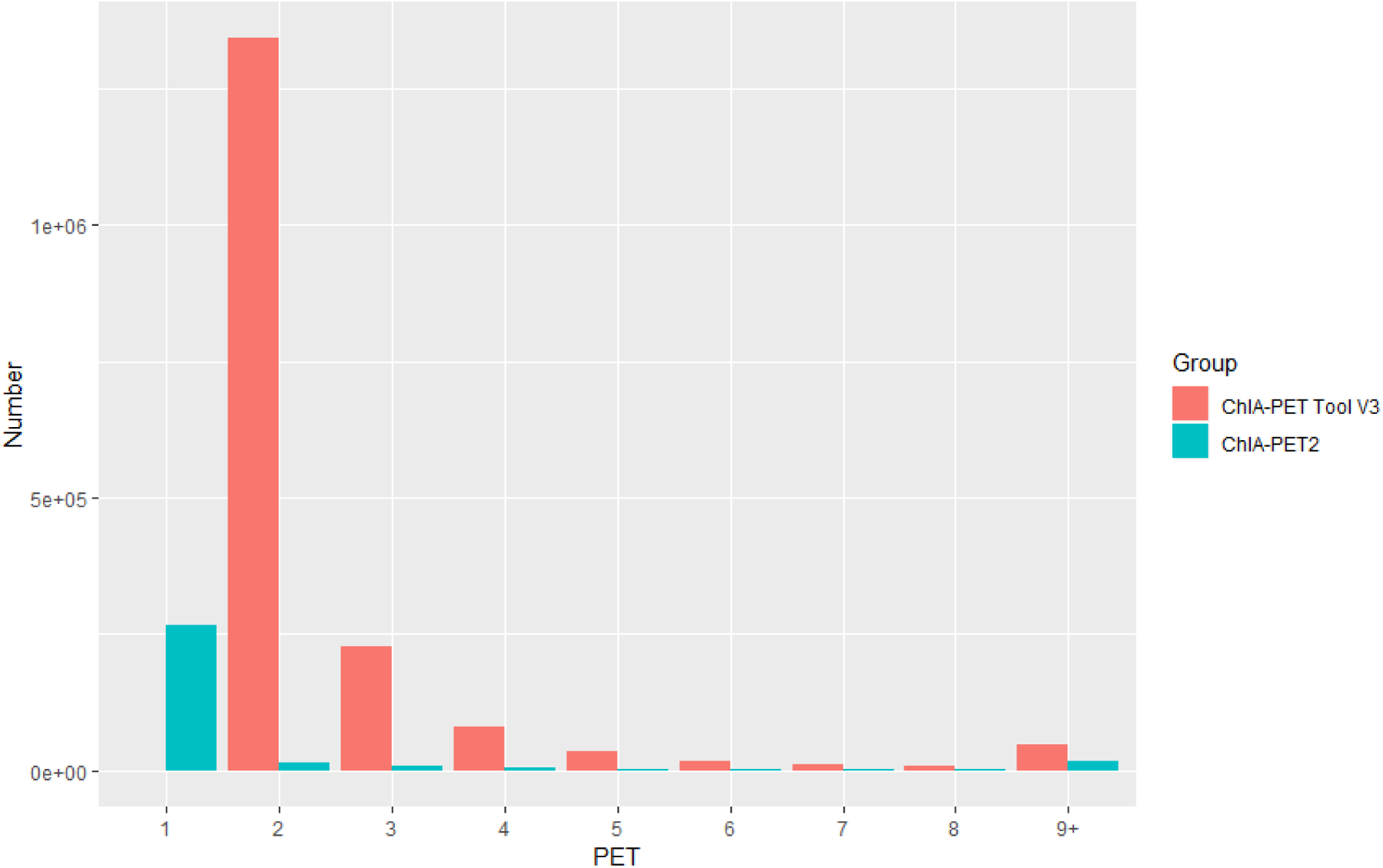
PET count distribution of significant interactions detected by different tools from human GM12878 cells.

