## Supplementary material for "Chromatin interaction analysis with updated ChIA-PET Tool (V3)": Rplot1.html

| Title | Information |
| --- | --- |
| Fastq file 1 | /home/tksun/tasks/data/K562\_RNAPII\_saturated\_1.fastq |
| Fastq file 2 | /home/tksun/tasks/data/K562\_RNAPII\_saturated\_2.fastq |
| Linker file | /home/tksun/tasks/ChIA-PET\_sun/program/../linker/linker.txt |
| LinkerA | GTTGGATAAG 7 4 |
| LinkerB | GTTGGAATGT 7 4 |
| Output folder | /home/tksun/tasks/data/test/K562\_RNAPII\_saturated |
| Output prefix | K562\_RNAPII\_saturated |
| Minimum linker alignment score | 8 |
| minimum tag length | 18 |
| maximum tag length | 1000 |
| Minumun SecondBestScore difference | 3 |
| Output data with ambiguous linker info | 1 |
