## Supplementary material for "Chromatin interaction analysis with updated ChIA-PET Tool (V3)": Rplot2.html

| PET category | Number | Percentage | Percentage of total PETs | Order |
| --- | --- | --- | --- | --- |
| Total PETs | 75,575,544 | NA | NA | (1) |
| Same-linker PETs after linker filtering | 68,573,721 | 90.74% of (1) | 90.74% of (1) | (2) |
| Uniquely Mapped same-linker PETs | 10,732,248 | 15.65% of (2) | 14.2% of (1) | (3) |
| Merging same same-linker PETs | 7,407,560 | 69.02% of (3) | 9.8% of (1) | (4) |
| Merging similar same-linker PETs | 7,388,164 | 99.74% of (4) | 9.78% of (1) | (5) |
| Self-ligation PETs | 2,387,126 | 32.31% of (5) | 3.16% of (1) | (6) |
| Inter-ligation PETs | 2,976,285 | 40.28% of (5) | 3.94% of (1) | (7) |
| Other PETs with short distance | 2,024,753 | 27.41% of (5) | 2.68% of (1) | (8) |
| Peaks from self-ligation | 6,332 | NA | NA | (9) |
| Interacting pairs | 31,011 | NA | NA | (10) |
