## Supplementary material for "Chromatin interaction analysis with updated ChIA-PET Tool (V3)": Rplot3_1.html

|  | Non-mappable | Uniquely-mapped | Others |
| --- | --- | --- | --- |
| Non-mappable | 429 | 8,055 | 22,517 |
| Uniquely-mapped | 7,435 | 10,732,248 | 15,237,351 |
| Others | 19,737 | 14,195,363 | 28,350,586 |
