## Supplementary material for "Chromatin interaction analysis with updated ChIA-PET Tool (V3)": Rplot7.html

| A\_A | A\_B | B\_A | B\_B | Ambiguous | Total |
| --- | --- | --- | --- | --- | --- |
| 33,029,693 | 420,122 | 425,932 | 35,544,028 | 6,155,769 | 75,575,544 |
| 43.70% | 0.56% | 0.56% | 47.03% | 8.15% | 100% |
