## Supplementary material for "Chromatin interaction analysis with updated ChIA-PET Tool (V3)": Rplot13.html

| Distance | Frequency | Interaction type |
| --- | --- | --- |
| < 100Kb | 24656 | Intra-chromosomal |
| [100Kb, 1Mb) | 5830 | Intra-chromosomal |
| [1Mb, 10Mb) | 131 | Intra-chromosomal |
| > 10Mb | 47 | Intra-chromosomal |
| Different chromosomes | 347 | Inter-chromosomal |
