## Supplementary material for "Chromatin interaction analysis with updated ChIA-PET Tool (V3)": Rplot21.html

| Title | Information |
| --- | --- |
| Self-ligation cutoff | 8000 |
| Min distance between peak | 500 |
| Peak mode (1: region, 2: summit) | 2 |
| Min coverage for peak | 5 |
| Genome length | 3e+09 |
| Genome coverage ratio | 0.8 |
| P-value cutoff | 1e-05 |
