## Supplementary material for "Chromatin interaction analysis with updated ChIA-PET Tool (V3)": Rplot24.html

| Module | Running time (min) |
| --- | --- |
| Linker filtering | 7.45 |
| Mapping | 128.75 |
| Removing redundancy | 19.65 |
| Categorization of PETs | 0.17 |
| Peak calling | 0.50 |
| Clustering | 4.35 |
| Total | 160.87 |
