## Supplementary material for "Chromatin interaction analysis with updated ChIA-PET Tool (V3)": Rplot25.html

| Title | Percentage |
| --- | --- |
| (same-linker PETs)/(Total PETs) | 90.74% |
| (Unique mapped same-linker PETs)/(Total PETs) | 14.2% |
| (PETs after removing redundancy)/(Total PETs) | 9.78% |
| (inter-ligation PETs)/(PETs after removing redundancy) | 40.28% |
| (intra-chromosomal inter-ligation PETs)/(inter-ligation PETs) | 98.88% |
| (intra-chromosomal inter-ligation PETs within 1Mb)/(intra-chromosomal inter-ligation PETs) | 98.31% |
