## Supplementary material for "Chromatin interaction analysis with updated ChIA-PET Tool (V3)": Rplot30.html

| PET.counts | No. of clusters | No intra chrom clusters | No inter chrom clusters | Percent of intra chrom clusters |
| --- | --- | --- | --- | --- |
| 2 | 22545 | 22240 | 305 | 98.65% |
| 3 | 4053 | 4035 | 18 | 99.56% |
| 4 | 1437 | 1432 | 5 | 99.65% |
| 5 | 749 | 745 | 4 | 99.47% |
| 6 | 453 | 448 | 5 | 98.9% |
| 7 | 298 | 296 | 2 | 99.33% |
| 8 | 238 | 236 | 2 | 99.16% |
| 9 | 173 | 173 | 0 | 100% |
| >=10 | 1065 | 1059 | 6 | 99.44% |
| Total | 31011 | 30664 | 347 | 98.88% |
