## Supplementary material for "Chromatin interaction analysis with updated ChIA-PET Tool (V3)": index.html

 


ChIA-PET Tool


- Running information
  - Running summary
  - Linker filtering
  - Mapping
  - Redundancy removal
  - Categorization of PETs
  - Peak calling
  - Clustering
  - Extra info
- Running statistics
  - Basic statistics
  - Mapping statistics
  - Running time
- Linker filtering report
  - Linker alignment score
  - Linker alignment score difference
  - Tag length
  - Linker composition
- Mapping report
  - Bedpe quality score
  - Bedpe strands
  - Filtered bedpe distribution
  - Unique intra-chrom strand distribution
- Span distribution report
  - Fragments length statistics
  - PETs span
  - Gap of PETs span
- Interactions visualization
  - PET counts
  - Interaction clusters classification
  - Intra-chromosomal inetractions visualization
  - Circular visualization


### Running information

##### Running summary

##### Linker filtering

##### Mapping

##### Remove redundancy

##### Category

##### Peak calling

##### Clustering

##### Extra Info

### Running statistics

##### Basic statistics

##### Mapping statistics of read1 (horizental) and read2 (vertical)

##### Running time

### Linker filtering report

##### Linker alignment score distribution

##### Linker alignment score difference distribution

##### Tag length distribution

##### Linker composition

### Mapping report

##### Mapping quality score distribution from Bedpe file

##### Mapping strands distribution from Bedpe file

##### Distribution of counts of PETs mapping to the same position in Bedpe file

##### Purified intra-chromosomal reads strands distribution from Bedpe file

### Span distribution

##### Fragment length statistics

##### PETs span distribution

##### Difference of PET span distributions between strands combination -+ and average number of (++, +- and --)

##### Log-log plot of difference of PET span distribution

### Cluster visualization

##### PET counts

##### Interaction clusters classification

##### Intra-chromosomal interactions visualization

Blue peaks above chromosomes are transcription factor binding sites
Red curves under chromosomes are intra-chromosomal interactions

##### Circular view

Circular view of inter-chromosomal interactions

1. Please use Chrome, Firefox or Safari for better browsing experience.

dd
