## Supplementary material for "Chromatin interaction analysis with updated ChIA-PET Tool (V3)": common.js

```
/*
	[Discuz!] (C)2001-2099 Comsenz Inc.
	This is NOT a freeware, use is subject to license terms

	$Id: common.js 32580 2013-02-22 03:40:28Z monkey $
*/

var BROWSER = {};
var USERAGENT = navigator.userAgent.toLowerCase();
browserVersion({'ie':'msie','firefox':'','chrome':'','opera':'','safari':'','mozilla':'','webkit':'','maxthon':'','qq':'qqbrowser'});
if(BROWSER.safari) {
	BROWSER.firefox = true;
}
BROWSER.opera = BROWSER.opera ? opera.version() : 0;

HTMLNODE = document.getElementsByTagName('head')[0].parentNode;
if(BROWSER.ie) {
	BROWSER.iemode = parseInt(typeof document.documentMode != 'undefined' ? document.documentMode : BROWSER.ie);
	HTMLNODE.className = 'ie_all ie' + BROWSER.iemode;
}

var CSSLOADED = [];
var JSLOADED = [];
var JSMENU = [];
JSMENU['active'] = [];
JSMENU['timer'] = [];
JSMENU['drag'] = [];
JSMENU['layer'] = 0;
JSMENU['zIndex'] = {'win':200,'menu':300,'dialog':400,'prompt':500};
JSMENU['float'] = '';
var CURRENTSTYPE = null;
var discuz_uid = isUndefined(discuz_uid) ? 0 : discuz_uid;
var creditnotice = isUndefined(creditnotice) ? '' : creditnotice;
var cookiedomain = isUndefined(cookiedomain) ? '' : cookiedomain;
var cookiepath = isUndefined(cookiepath) ? '' : cookiepath;
var EXTRAFUNC = [], EXTRASTR = '';
EXTRAFUNC['showmenu'] = [];

var DISCUZCODE = [];
DISCUZCODE['num'] = '-1';
DISCUZCODE['html'] = [];

var USERABOUT_BOX = true;
var USERCARDST = null;
var CLIPBOARDSWFDATA = '';
var NOTICETITLE = [];

if(BROWSER.firefox && window.HTMLElement) {
	HTMLElement.prototype.__defineGetter__( "innerText", function(){
		var anyString = "";
		var childS = this.childNodes;
		for(var i=0; i <childS.length; i++) {
			if(childS[i].nodeType==1) {
				anyString += childS[i].tagName=="BR" ? '\n' : childS[i].innerText;
			} else if(childS[i].nodeType==3) {
				anyString += childS[i].nodeValue;
			}
		}
		return anyString;
	});
	HTMLElement.prototype.__defineSetter__( "innerText", function(sText){
		this.textContent=sText;
	});
	HTMLElement.prototype.__defineSetter__('outerHTML', function(sHTML) {
			var r = this.ownerDocument.createRange();
		r.setStartBefore(this);
		var df = r.createContextualFragment(sHTML);
		this.parentNode.replaceChild(df,this);
		return sHTML;
	});

	HTMLElement.prototype.__defineGetter__('outerHTML', function() {
		var attr;
		var attrs = this.attributes;
		var str = '<' + this.tagName.toLowerCase();
		for(var i = 0;i < attrs.length;i++){
			attr = attrs[i];
			if(attr.specified)
			str += ' ' + attr.name + '="' + attr.value + '"';
		}
		if(!this.canHaveChildren) {
			return str + '>';
		}
		return str + '>' + this.innerHTML + '</' + this.tagName.toLowerCase() + '>';
		});

	HTMLElement.prototype.__defineGetter__('canHaveChildren', function() {
		switch(this.tagName.toLowerCase()) {
			case 'area':case 'base':case 'basefont':case 'col':case 'frame':case 'hr':case 'img':case 'br':case 'input':case 'isindex':case 'link':case 'meta':case 'param':
			return false;
			}
		return true;
	});
}

function $(id) {
	return !id ? null : document.getElementById(id);
}

function $C(classname, ele, tag) {
	var returns = [];
	ele = ele || document;
	tag = tag || '*';
	if(ele.getElementsByClassName) {
		var eles = ele.getElementsByClassName(classname);
		if(tag != '*') {
			for (var i = 0, L = eles.length; i < L; i++) {
				if (eles[i].tagName.toLowerCase() == tag.toLowerCase()) {
						returns.push(eles[i]);
				}
			}
		} else {
			returns = eles;
		}
	}else {
		eles = ele.getElementsByTagName(tag);
		var pattern = new RegExp("(^|\\s)"+classname+"(\\s|$)");
		for (i = 0, L = eles.length; i < L; i++) {
				if (pattern.test(eles[i].className)) {
						returns.push(eles[i]);
				}
		}
	}
	return returns;
}

function _attachEvent(obj, evt, func, eventobj) {
	eventobj = !eventobj ? obj : eventobj;
	if(obj.addEventListener) {
		obj.addEventListener(evt, func, false);
	} else if(eventobj.attachEvent) {
		obj.attachEvent('on' + evt, func);
	}
}

function _detachEvent(obj, evt, func, eventobj) {
	eventobj = !eventobj ? obj : eventobj;
	if(obj.removeEventListener) {
		obj.removeEventListener(evt, func, false);
	} else if(eventobj.detachEvent) {
		obj.detachEvent('on' + evt, func);
	}
}

function browserVersion(types) {
	var other = 1;
	for(i in types) {
		var v = types[i] ? types[i] : i;
		if(USERAGENT.indexOf(v) != -1) {
			var re = new RegExp(v + '(\\/|\\s)([\\d\\.]+)', 'ig');
			var matches = re.exec(USERAGENT);
			var ver = matches != null ? matches[2] : 0;
			other = ver !== 0 && v != 'mozilla' ? 0 : other;
		}else {
			var ver = 0;
		}
		eval('BROWSER.' + i + '= ver');
	}
	BROWSER.other = other;
}

function getEvent() {
	if(document.all) return window.event;
	func = getEvent.caller;
	while(func != null) {
		var arg0 = func.arguments[0];
		if (arg0) {
			if((arg0.constructor  == Event || arg0.constructor == MouseEvent) || (typeof(arg0) == "object" && arg0.preventDefault && arg0.stopPropagation)) {
				return arg0;
			}
		}
		func=func.caller;
	}
	return null;
}

function isUndefined(variable) {
	return typeof variable == 'undefined' ? true : false;
}

function in_array(needle, haystack) {
	if(typeof needle == 'string' || typeof needle == 'number') {
		for(var i in haystack) {
			if(haystack[i] == needle) {
					return true;
			}
		}
	}
	return false;
}

function trim(str) {
	return (str + '').replace(/(\s+)$/g, '').replace(/^\s+/g, '');
}

function strlen(str) {
	return (BROWSER.ie && str.indexOf('\n') != -1) ? str.replace(/\r?\n/g, '_').length : str.length;
}

function mb_strlen(str) {
	var len = 0;
	for(var i = 0; i < str.length; i++) {
		len += str.charCodeAt(i) < 0 || str.charCodeAt(i) > 255 ? (charset == 'utf-8' ? 3 : 2) : 1;
	}
	return len;
}

function mb_cutstr(str, maxlen, dot) {
	var len = 0;
	var ret = '';
	var dot = !dot ? '...' : dot;
	maxlen = maxlen - dot.length;
	for(var i = 0; i < str.length; i++) {
		len += str.charCodeAt(i) < 0 || str.charCodeAt(i) > 255 ? (charset == 'utf-8' ? 3 : 2) : 1;
		if(len > maxlen) {
			ret += dot;
			break;
		}
		ret += str.substr(i, 1);
	}
	return ret;
}

function preg_replace(search, replace, str, regswitch) {
	var regswitch = !regswitch ? 'ig' : regswitch;
	var len = search.length;
	for(var i = 0; i < len; i++) {
		re = new RegExp(search[i], regswitch);
		str = str.replace(re, typeof replace == 'string' ? replace : (replace[i] ? replace[i] : replace[0]));
	}
	return str;
}

function htmlspecialchars(str) {
	return preg_replace(['&', '<', '>', '"'], ['&amp;', '&lt;', '&gt;', '&quot;'], str);
}

function display(id) {
	var obj = $(id);
	if(obj.style.visibility) {
		obj.style.visibility = obj.style.visibility == 'visible' ? 'hidden' : 'visible';
	} else {
		obj.style.display = obj.style.display == '' ? 'none' : '';
	}
}

function checkall(form, prefix, checkall) {
	var checkall = checkall ? checkall : 'chkall';
	count = 0;
	for(var i = 0; i < form.elements.length; i++) {
		var e = form.elements[i];
		if(e.name && e.name != checkall && !e.disabled && (!prefix || (prefix && e.name.match(prefix)))) {
			e.checked = form.elements[checkall].checked;
			if(e.checked) {
				count++;
			}
		}
	}
	return count;
}

function setcookie(cookieName, cookieValue, seconds, path, domain, secure) {
	var expires = new Date();
	if(cookieValue == '' || seconds < 0) {
		cookieValue = '';
		seconds = -2592000;
	}
	expires.setTime(expires.getTime() + seconds * 1000);
	domain = !domain ? cookiedomain : domain;
	path = !path ? cookiepath : path;
	document.cookie = escape(cookiepre + cookieName) + '=' + escape(cookieValue)
		+ (expires ? '; expires=' + expires.toGMTString() : '')
		+ (path ? '; path=' + path : '/')
		+ (domain ? '; domain=' + domain : '')
		+ (secure ? '; secure' : '');
}

function getcookie(name, nounescape) {
	name = cookiepre + name;
	var cookie_start = document.cookie.indexOf(name);
	var cookie_end = document.cookie.indexOf(";", cookie_start);
	if(cookie_start == -1) {
		return '';
	} else {
		var v = document.cookie.substring(cookie_start + name.length + 1, (cookie_end > cookie_start ? cookie_end : document.cookie.length));
		return !nounescape ? unescape(v) : v;
	}
}

function Ajax(recvType, waitId) {

	var aj = new Object();

	aj.loading = 'ÇëÉÔºò...';
	aj.recvType = recvType ? recvType : 'XML';
	aj.waitId = waitId ? $(waitId) : null;

	aj.resultHandle = null;
	aj.sendString = '';
	aj.targetUrl = '';

	aj.setLoading = function(loading) {
		if(typeof loading !== 'undefined' && loading !== null) aj.loading = loading;
	};

	aj.setRecvType = function(recvtype) {
		aj.recvType = recvtype;
	};

	aj.setWaitId = function(waitid) {
		aj.waitId = typeof waitid == 'object' ? waitid : $(waitid);
	};

	aj.createXMLHttpRequest = function() {
		var request = false;
		if(window.XMLHttpRequest) {
			request = new XMLHttpRequest();
			if(request.overrideMimeType) {
				request.overrideMimeType('text/xml');
			}
		} else if(window.ActiveXObject) {
			var versions = ['Microsoft.XMLHTTP', 'MSXML.XMLHTTP', 'Microsoft.XMLHTTP', 'Msxml2.XMLHTTP.7.0', 'Msxml2.XMLHTTP.6.0', 'Msxml2.XMLHTTP.5.0', 'Msxml2.XMLHTTP.4.0', 'MSXML2.XMLHTTP.3.0', 'MSXML2.XMLHTTP'];
			for(var i=0; i<versions.length; i++) {
				try {
					request = new ActiveXObject(versions[i]);
					if(request) {
						return request;
					}
				} catch(e) {}
			}
		}
		return request;
	};

	aj.XMLHttpRequest = aj.createXMLHttpRequest();
	aj.showLoading = function() {
		if(aj.waitId && (aj.XMLHttpRequest.readyState != 4 || aj.XMLHttpRequest.status != 200)) {
			aj.waitId.style.display = '';
			aj.waitId.innerHTML = '<span><img src="'+ IMGDIR + '/loading.gif" class="vm"> ' + aj.loading + '</span>';
		}
	};

	aj.processHandle = function() {
		if(aj.XMLHttpRequest.readyState == 4 && aj.XMLHttpRequest.status == 200) {
			if(aj.waitId) {
				aj.waitId.style.display = 'none';
			}
			if(aj.recvType == 'HTML') {
				aj.resultHandle(aj.XMLHttpRequest.responseText, aj);
			} else if(aj.recvType == 'XML') {
				if(!aj.XMLHttpRequest.responseXML || !aj.XMLHttpRequest.responseXML.lastChild || aj.XMLHttpRequest.responseXML.lastChild.localName == 'parsererror') {
					aj.resultHandle('<a href="' + aj.targetUrl + '" target="_blank" style="color:red">ÄÚ²¿´íÎó£¬ÎÞ·¨ÏÔÊ¾´ËÄÚÈÝ</a>' , aj);
				} else {
					aj.resultHandle(aj.XMLHttpRequest.responseXML.lastChild.firstChild.nodeValue, aj);
				}
			}
		}
	};

	aj.get = function(targetUrl, resultHandle) {
		targetUrl = hostconvert(targetUrl);
		setTimeout(function(){aj.showLoading()}, 250);
		aj.targetUrl = targetUrl;
		aj.XMLHttpRequest.onreadystatechange = aj.processHandle;
		aj.resultHandle = resultHandle;
		var attackevasive = isUndefined(attackevasive) ? 0 : attackevasive;
		if(window.XMLHttpRequest) {
			aj.XMLHttpRequest.open('GET', aj.targetUrl);
			aj.XMLHttpRequest.setRequestHeader('X-Requested-With', 'XMLHttpRequest');
			aj.XMLHttpRequest.send(null);
		} else {
			aj.XMLHttpRequest.open("GET", targetUrl, true);
			aj.XMLHttpRequest.setRequestHeader('X-Requested-With', 'XMLHttpRequest');
			aj.XMLHttpRequest.send();
		}
	};
	aj.post = function(targetUrl, sendString, resultHandle) {
		targetUrl = hostconvert(targetUrl);
		setTimeout(function(){aj.showLoading()}, 250);
		aj.targetUrl = targetUrl;
		aj.sendString = sendString;
		aj.XMLHttpRequest.onreadystatechange = aj.processHandle;
		aj.resultHandle = resultHandle;
		aj.XMLHttpRequest.open('POST', targetUrl);
		aj.XMLHttpRequest.setRequestHeader('Content-Type', 'application/x-www-form-urlencoded');
		aj.XMLHttpRequest.setRequestHeader('X-Requested-With', 'XMLHttpRequest');
		aj.XMLHttpRequest.send(aj.sendString);
	};
	return aj;
}

function getHost(url) {
	var host = "null";
	if(typeof url == "undefined"|| null == url) {
		url = window.location.href;
	}
	var regex = /^\w+\:\/\/([^\/]*).*/;
	var match = url.match(regex);
	if(typeof match != "undefined" && null != match) {
		host = match[1];
	}
	return host;
}

function hostconvert(url) {
	if(!url.match(/^https?:\/\//)) url = SITEURL + url;
	var url_host = getHost(url);
	var cur_host = getHost().toLowerCase();
	if(url_host && cur_host != url_host) {
		url = url.replace(url_host, cur_host);
	}
	return url;
}

function newfunction(func) {
	var args = [];
	for(var i=1; i<arguments.length; i++) args.push(arguments[i]);
	return function(event) {
		doane(event);
		window[func].apply(window, args);
		return false;
	}
}

function evalscript(s) {
	if(s.indexOf('<script') == -1) return s;
	var p = /<script[^\>]*?>([^\x00]*?)<\/script>/ig;
	var arr = [];
	while(arr = p.exec(s)) {
		var p1 = /<script[^\>]*?src=\"([^\>]*?)\"[^\>]*?(reload=\"1\")?(?:charset=\"([\w\-]+?)\")?><\/script>/i;
		var arr1 = [];
		arr1 = p1.exec(arr[0]);
		if(arr1) {
			appendscript(arr1[1], '', arr1[2], arr1[3]);
		} else {
			p1 = /<script(.*?)>([^\x00]+?)<\/script>/i;
			arr1 = p1.exec(arr[0]);
			appendscript('', arr1[2], arr1[1].indexOf('reload=') != -1);
		}
	}
	return s;
}

var safescripts = {}, evalscripts = [];
function safescript(id, call, seconds, times, timeoutcall, endcall, index) {
	seconds = seconds || 1000;
	times = times || 0;
	var checked = true;
	try {
		if(typeof call == 'function') {
			call();
		} else {
			eval(call);
		}
	} catch(e) {
		checked = false;
	}
	if(!checked) {
		if(!safescripts[id] || !index) {
			safescripts[id] = safescripts[id] || [];
			safescripts[id].push({
				'times':0,
				'si':setInterval(function () {
					safescript(id, call, seconds, times, timeoutcall, endcall, safescripts[id].length);
				}, seconds)
			});
		} else {
			index = (index || 1) - 1;
			safescripts[id][index]['times']++;
			if(safescripts[id][index]['times'] >= times) {
				clearInterval(safescripts[id][index]['si']);
				if(typeof timeoutcall == 'function') {
					timeoutcall();
				} else {
					eval(timeoutcall);
				}
			}
		}
	} else {
		try {
			index = (index || 1) - 1;
			if(safescripts[id][index]['si']) {
				clearInterval(safescripts[id][index]['si']);
			}
			if(typeof endcall == 'function') {
				endcall();
			} else {
				eval(endcall);
			}
		} catch(e) {}
	}
}

function $F(func, args, script) {
	var run = function () {
		var argc = args.length, s = '';
		for(i = 0;i < argc;i++) {
			s += ',args[' + i + ']';
		}
		eval('var check = typeof ' + func + ' == \'function\'');
		if(check) {
			eval(func + '(' + s.substr(1) + ')');
		} else {
			setTimeout(function () { checkrun(); }, 50);
		}
	};
	var checkrun = function () {
		if(JSLOADED[src]) {
			run();
		} else {
			setTimeout(function () { checkrun(); }, 50);
		}
	};
	script = script || 'common_extra';
	src = JSPATH + script + '.js?' + VERHASH;
	if(!JSLOADED[src]) {
		appendscript(src);
	}
	checkrun();
}

function appendscript(src, text, reload, charset) {
	var id = hash(src + text);
	if(!reload && in_array(id, evalscripts)) return;
	if(reload && $(id)) {
		$(id).parentNode.removeChild($(id));
	}

	evalscripts.push(id);
	var scriptNode = document.createElement("script");
	scriptNode.type = "text/javascript";
	scriptNode.id = id;
	scriptNode.charset = charset ? charset : (BROWSER.firefox ? document.characterSet : document.charset);
	try {
		if(src) {
			scriptNode.src = src;
			scriptNode.onloadDone = false;
			scriptNode.onload = function () {
				scriptNode.onloadDone = true;
				JSLOADED[src] = 1;
			};
			scriptNode.onreadystatechange = function () {
				if((scriptNode.readyState == 'loaded' || scriptNode.readyState == 'complete') && !scriptNode.onloadDone) {
					scriptNode.onloadDone = true;
					JSLOADED[src] = 1;
				}
			};
		} else if(text){
			scriptNode.text = text;
		}
		document.getElementsByTagName('head')[0].appendChild(scriptNode);
	} catch(e) {}
}

function stripscript(s) {
	return s.replace(/<script.*?>.*?<\/script>/ig, '');
}

function ajaxupdateevents(obj, tagName) {
	tagName = tagName ? tagName : 'A';
	var objs = obj.getElementsByTagName(tagName);
	for(k in objs) {
		var o = objs[k];
		ajaxupdateevent(o);
	}
}

function ajaxupdateevent(o) {
	if(typeof o == 'object' && o.getAttribute) {
		if(o.getAttribute('ajaxtarget')) {
			if(!o.id) o.id = Math.random();
			var ajaxevent = o.getAttribute('ajaxevent') ? o.getAttribute('ajaxevent') : 'click';
			var ajaxurl = o.getAttribute('ajaxurl') ? o.getAttribute('ajaxurl') : o.href;
			_attachEvent(o, ajaxevent, newfunction('ajaxget', ajaxurl, o.getAttribute('ajaxtarget'), o.getAttribute('ajaxwaitid'), o.getAttribute('ajaxloading'), o.getAttribute('ajaxdisplay')));
			if(o.getAttribute('ajaxfunc')) {
				o.getAttribute('ajaxfunc').match(/(\w+)\((.+?)\)/);
				_attachEvent(o, ajaxevent, newfunction(RegExp.$1, RegExp.$2));
			}
		}
	}
}

function ajaxget(url, showid, waitid, loading, display, recall) {
	waitid = typeof waitid == 'undefined' || waitid === null ? showid : waitid;
	var x = new Ajax();
	x.setLoading(loading);
	x.setWaitId(waitid);
	x.display = typeof display == 'undefined' || display == null ? '' : display;
	x.showId = $(showid);

	if(url.substr(strlen(url) - 1) == '#') {
		url = url.substr(0, strlen(url) - 1);
		x.autogoto = 1;
	}

	var url = url + '&inajax=1&ajaxtarget=' + showid;
	x.get(url, function(s, x) {
		var evaled = false;
		if(s.indexOf('ajaxerror') != -1) {
			evalscript(s);
			evaled = true;
		}
		if(!evaled && (typeof ajaxerror == 'undefined' || !ajaxerror)) {
			if(x.showId) {
				x.showId.style.display = x.display;
				ajaxinnerhtml(x.showId, s);
				ajaxupdateevents(x.showId);
				if(x.autogoto) scroll(0, x.showId.offsetTop);
			}
		}

		ajaxerror = null;
		if(recall && typeof recall == 'function') {
			recall();
		} else if(recall) {
			eval(recall);
		}
		if(!evaled) evalscript(s);
	});
}

function ajaxpost(formid, showid, waitid, showidclass, submitbtn, recall) {
	var waitid = typeof waitid == 'undefined' || waitid === null ? showid : (waitid !== '' ? waitid : '');
	var showidclass = !showidclass ? '' : showidclass;
	var ajaxframeid = 'ajaxframe';
	var ajaxframe = $(ajaxframeid);
	var curform = $(formid);
	var formtarget = curform.target;

	var handleResult = function() {
		var s = '';
		var evaled = false;

		showloading('none');
		try {
			s = $(ajaxframeid).contentWindow.document.XMLDocument.text;
		} catch(e) {
			try {
				s = $(ajaxframeid).contentWindow.document.documentElement.firstChild.wholeText;
			} catch(e) {
				try {
					s = $(ajaxframeid).contentWindow.document.documentElement.firstChild.nodeValue;
				} catch(e) {
					s = 'ÄÚ²¿´íÎó£¬ÎÞ·¨ÏÔÊ¾´ËÄÚÈÝ';
				}
			}
		}
		if(s != '' && s.indexOf('ajaxerror') != -1) {
			evalscript(s);
			evaled = true;
		}
		if(showidclass) {
			if(showidclass != 'onerror') {
				$(showid).className = showidclass;
			} else {
				showError(s);
				ajaxerror = true;
			}
		}
		if(submitbtn) {
			submitbtn.disabled = false;
		}
		if(!evaled && (typeof ajaxerror == 'undefined' || !ajaxerror)) {
			ajaxinnerhtml($(showid), s);
		}
		ajaxerror = null;
		if($(formid)) $(formid).target = formtarget;
		if(typeof recall == 'function') {
			recall();
		} else {
			eval(recall);
		}
		if(!evaled) evalscript(s);
		ajaxframe.loading = 0;
		if(!BROWSER.firefox) {
			$('append_parent').removeChild(ajaxframe.parentNode);
		} else {
			setTimeout(
				function(){
					$('append_parent').removeChild(ajaxframe.parentNode);
				},
				100
			);
		}
	};
	if(!ajaxframe) {
		var div = document.createElement('div');
		div.style.display = 'none';
		div.innerHTML = '<iframe name="' + ajaxframeid + '" id="' + ajaxframeid + '" loading="1"></iframe>';
		$('append_parent').appendChild(div);
		ajaxframe = $(ajaxframeid);
	} else if(ajaxframe.loading) {
		return false;
	}

	_attachEvent(ajaxframe, 'load', handleResult);

	showloading();
	curform.target = ajaxframeid;
	var action = curform.getAttribute('action');
	action = hostconvert(action);
	curform.action = action.replace(/\&inajax\=1/g, '')+'&inajax=1';
	curform.submit();
	if(submitbtn) {
		submitbtn.disabled = true;
	}
	doane();
	return false;
}

function ajaxmenu(ctrlObj, timeout, cache, duration, pos, recall, idclass, contentclass) {
	if(!ctrlObj.getAttribute('mid')) {
		var ctrlid = ctrlObj.id;
		if(!ctrlid) {
			ctrlObj.id = 'ajaxid_' + Math.random();
		}
	} else {
		var ctrlid = ctrlObj.getAttribute('mid');
		if(!ctrlObj.id) {
			ctrlObj.id = 'ajaxid_' + Math.random();
		}
	}
	var menuid = ctrlid + '_menu';
	var menu = $(menuid);
	if(isUndefined(timeout)) timeout = 3000;
	if(isUndefined(cache)) cache = 1;
	if(isUndefined(pos)) pos = '43';
	if(isUndefined(duration)) duration = timeout > 0 ? 0 : 3;
	if(isUndefined(idclass)) idclass = 'p_pop';
	if(isUndefined(contentclass)) contentclass = 'p_opt';
	var func = function() {
		showMenu({'ctrlid':ctrlObj.id,'menuid':menuid,'duration':duration,'timeout':timeout,'pos':pos,'cache':cache,'layer':2});
		if(typeof recall == 'function') {
			recall();
		} else {
			eval(recall);
		}
	};

	if(menu) {
		if(menu.style.display == '') {
			hideMenu(menuid);
		} else {
			func();
		}
	} else {
		menu = document.createElement('div');
		menu.id = menuid;
		menu.style.display = 'none';
		menu.className = idclass;
		menu.innerHTML = '<div class="' + contentclass + '" id="' + menuid + '_content"></div>';
		$('append_parent').appendChild(menu);
		var url = (!isUndefined(ctrlObj.attributes['shref']) ? ctrlObj.attributes['shref'].value : (!isUndefined(ctrlObj.href) ? ctrlObj.href : ctrlObj.attributes['href'].value));
		url += (url.indexOf('?') != -1 ? '&' :'?') + 'ajaxmenu=1';
		ajaxget(url, menuid + '_content', 'ajaxwaitid', '', '', func);
	}
	doane();
}

function hash(string, length) {
	var length = length ? length : 32;
	var start = 0;
	var i = 0;
	var result = '';
	filllen = length - string.length % length;
	for(i = 0; i < filllen; i++){
		string += "0";
	}
	while(start < string.length) {
		result = stringxor(result, string.substr(start, length));
		start += length;
	}
	return result;
}

function stringxor(s1, s2) {
	var s = '';
	var hash = 'abcdefghijklmnopqrstuvwxyzABCDEFGHIJKLMNOPQRSTUVWXYZ';
	var max = Math.max(s1.length, s2.length);
	for(var i=0; i<max; i++) {
		var k = s1.charCodeAt(i) ^ s2.charCodeAt(i);
		s += hash.charAt(k % 52);
	}
	return s;
}

function showPreview(val, id) {
	var showObj = $(id);
	if(showObj) {
		showObj.innerHTML = val.replace(/\n/ig, "<bupdateseccoder />");
	}
}

function showloading(display, waiting) {
	var display = display ? display : 'block';
	var waiting = waiting ? waiting : 'ÇëÉÔºò...';
	$('ajaxwaitid').innerHTML = waiting;
	$('ajaxwaitid').style.display = display;
}

function ajaxinnerhtml(showid, s) {
	if(showid.tagName != 'TBODY') {
		showid.innerHTML = s;
	} else {
		while(showid.firstChild) {
			showid.firstChild.parentNode.removeChild(showid.firstChild);
		}
		var div1 = document.createElement('DIV');
		div1.id = showid.id+'_div';
		div1.innerHTML = '<table><tbody id="'+showid.id+'_tbody">'+s+'</tbody></table>';
		$('append_parent').appendChild(div1);
		var trs = div1.getElementsByTagName('TR');
		var l = trs.length;
		for(var i=0; i<l; i++) {
			showid.appendChild(trs[0]);
		}
		var inputs = div1.getElementsByTagName('INPUT');
		var l = inputs.length;
		for(var i=0; i<l; i++) {
			showid.appendChild(inputs[0]);
		}
		div1.parentNode.removeChild(div1);
	}
}

function doane(event, preventDefault, stopPropagation) {
	var preventDefault = isUndefined(preventDefault) ? 1 : preventDefault;
	var stopPropagation = isUndefined(stopPropagation) ? 1 : stopPropagation;
	e = event ? event : window.event;
	if(!e) {
		e = getEvent();
	}
	if(!e) {
		return null;
	}
	if(preventDefault) {
		if(e.preventDefault) {
			e.preventDefault();
		} else {
			e.returnValue = false;
		}
	}
	if(stopPropagation) {
		if(e.stopPropagation) {
			e.stopPropagation();
		} else {
			e.cancelBubble = true;
		}
	}
	return e;
}


function showMenu(v) {
	var ctrlid = isUndefined(v['ctrlid']) ? v : v['ctrlid'];
	var showid = isUndefined(v['showid']) ? ctrlid : v['showid'];
	var menuid = isUndefined(v['menuid']) ? showid + '_menu' : v['menuid'];
	var ctrlObj = $(ctrlid);
	var menuObj = $(menuid);
	if(!menuObj) return;
	var mtype = isUndefined(v['mtype']) ? 'menu' : v['mtype'];
	var evt = isUndefined(v['evt']) ? 'mouseover' : v['evt'];
	var pos = isUndefined(v['pos']) ? '43' : v['pos'];
	var layer = isUndefined(v['layer']) ? 1 : v['layer'];
	var duration = isUndefined(v['duration']) ? 2 : v['duration'];
	var timeout = isUndefined(v['timeout']) ? 250 : v['timeout'];
	var maxh = isUndefined(v['maxh']) ? 600 : v['maxh'];
	var cache = isUndefined(v['cache']) ? 1 : v['cache'];
	var drag = isUndefined(v['drag']) ? '' : v['drag'];
	var dragobj = drag && $(drag) ? $(drag) : menuObj;
	var fade = isUndefined(v['fade']) ? 0 : v['fade'];
	var cover = isUndefined(v['cover']) ? 0 : v['cover'];
	var zindex = isUndefined(v['zindex']) ? JSMENU['zIndex']['menu'] : v['zindex'];
	var ctrlclass = isUndefined(v['ctrlclass']) ? '' : v['ctrlclass'];
	var winhandlekey = isUndefined(v['win']) ? '' : v['win'];
	zindex = cover ? zindex + 500 : zindex;
	if(typeof JSMENU['active'][layer] == 'undefined') {
		JSMENU['active'][layer] = [];
	}

	for(i in EXTRAFUNC['showmenu']) {
		try {
			eval(EXTRAFUNC['showmenu'][i] + '()');
		} catch(e) {}
	}

	if(evt == 'click' && in_array(menuid, JSMENU['active'][layer]) && mtype != 'win') {
		hideMenu(menuid, mtype);
		return;
	}
	if(mtype == 'menu') {
		hideMenu(layer, mtype);
	}

	if(ctrlObj) {
		if(!ctrlObj.getAttribute('initialized')) {
			ctrlObj.setAttribute('initialized', true);
			ctrlObj.unselectable = true;

			ctrlObj.outfunc = typeof ctrlObj.onmouseout == 'function' ? ctrlObj.onmouseout : null;
			ctrlObj.onmouseout = function() {
				if(this.outfunc) this.outfunc();
				if(duration < 3 && !JSMENU['timer'][menuid]) {
					JSMENU['timer'][menuid] = setTimeout(function () {
						hideMenu(menuid, mtype);
					}, timeout);
				}
			};

			ctrlObj.overfunc = typeof ctrlObj.onmouseover == 'function' ? ctrlObj.onmouseover : null;
			ctrlObj.onmouseover = function(e) {
				doane(e);
				if(this.overfunc) this.overfunc();
				if(evt == 'click') {
					clearTimeout(JSMENU['timer'][menuid]);
					JSMENU['timer'][menuid] = null;
				} else {
					for(var i in JSMENU['timer']) {
						if(JSMENU['timer'][i]) {
							clearTimeout(JSMENU['timer'][i]);
							JSMENU['timer'][i] = null;
						}
					}
				}
			};
		}
	}

	if(!menuObj.getAttribute('initialized')) {
		menuObj.setAttribute('initialized', true);
		menuObj.ctrlkey = ctrlid;
		menuObj.mtype = mtype;
		menuObj.layer = layer;
		menuObj.cover = cover;
		if(ctrlObj && ctrlObj.getAttribute('fwin')) {menuObj.scrolly = true;}
		menuObj.style.position = 'absolute';
		menuObj.style.zIndex = zindex + layer;
		menuObj.onclick = function(e) {
			return doane(e, 0, 1);
		};
		if(duration < 3) {
			if(duration > 1) {
				menuObj.onmouseover = function() {
					clearTimeout(JSMENU['timer'][menuid]);
					JSMENU['timer'][menuid] = null;
				};
			}
			if(duration != 1) {
				menuObj.onmouseout = function() {
					JSMENU['timer'][menuid] = setTimeout(function () {
						hideMenu(menuid, mtype);
					}, timeout);
				};
			}
		}
		if(cover) {
			var coverObj = document.createElement('div');
			coverObj.id = menuid + '_cover';
			coverObj.style.position = 'absolute';
			coverObj.style.zIndex = menuObj.style.zIndex - 1;
			coverObj.style.left = coverObj.style.top = '0px';
			coverObj.style.width = '100%';
			coverObj.style.height = Math.max(document.documentElement.clientHeight, document.body.offsetHeight) + 'px';
			coverObj.style.backgroundColor = '#000';
			coverObj.style.filter = 'progid:DXImageTransform.Microsoft.Alpha(opacity=50)';
			coverObj.style.opacity = 0.5;
			coverObj.onclick = function () { hideMenu(); };
			$('append_parent').appendChild(coverObj);
			_attachEvent(window, 'load', function () {
				coverObj.style.height = Math.max(document.documentElement.clientHeight, document.body.offsetHeight) + 'px';
			}, document);
		}
	}
	if(drag) {
		dragobj.style.cursor = 'move';
		dragobj.onmousedown = function(event) {try{dragMenu(menuObj, event, 1);}catch(e){}};
	}

	if(cover) $(menuid + '_cover').style.display = '';
	if(fade) {
		var O = 0;
		var fadeIn = function(O) {
			if(O > 100) {
				clearTimeout(fadeInTimer);
				return;
			}
			menuObj.style.filter = 'progid:DXImageTransform.Microsoft.Alpha(opacity=' + O + ')';
			menuObj.style.opacity = O / 100;
			O += 20;
			var fadeInTimer = setTimeout(function () {
				fadeIn(O);
			}, 40);
		};
		fadeIn(O);
		menuObj.fade = true;
	} else {
		menuObj.fade = false;
	}
	menuObj.style.display = '';
	if(ctrlObj && ctrlclass) {
		ctrlObj.className += ' ' + ctrlclass;
		menuObj.setAttribute('ctrlid', ctrlid);
		menuObj.setAttribute('ctrlclass', ctrlclass);
	}
	if(pos != '*') {
		setMenuPosition(showid, menuid, pos);
	}
	if(BROWSER.ie && BROWSER.ie < 7 && winhandlekey && $('fwin_' + winhandlekey)) {
		$(menuid).style.left = (parseInt($(menuid).style.left) - parseInt($('fwin_' + winhandlekey).style.left)) + 'px';
		$(menuid).style.top = (parseInt($(menuid).style.top) - parseInt($('fwin_' + winhandlekey).style.top)) + 'px';
	}
	if(maxh && menuObj.scrollHeight > maxh) {
		menuObj.style.height = maxh + 'px';
		if(BROWSER.opera) {
			menuObj.style.overflow = 'auto';
		} else {
			menuObj.style.overflowY = 'auto';
		}
	}

	if(!duration) {
		setTimeout('hideMenu(\'' + menuid + '\', \'' + mtype + '\')', timeout);
	}

	if(!in_array(menuid, JSMENU['active'][layer])) JSMENU['active'][layer].push(menuid);
	menuObj.cache = cache;
	if(layer > JSMENU['layer']) {
		JSMENU['layer'] = layer;
	}
	var hasshow = function(ele) {
		while(ele.parentNode && ((typeof(ele['currentStyle']) === 'undefined') ? window.getComputedStyle(ele,null) : ele['currentStyle'])['display'] !== 'none') {
			ele = ele.parentNode;
		}
		if(ele === document) {
			return true;
		} else {
			return false;
		}
	};
	if(!menuObj.getAttribute('disautofocus')) {
		try{
			var focused = false;
			var tags = ['input', 'select', 'textarea', 'button', 'a'];
			for(var i = 0; i < tags.length; i++) {
				var _all = menuObj.getElementsByTagName(tags[i]);
				if(_all.length) {
					for(j = 0; j < _all.length; j++) {
						if((!_all[j]['type'] || _all[j]['type'] != 'hidden') && hasshow(_all[j])) {
							_all[j].className += ' hidefocus';
							_all[j].focus();
							focused = true;
							var cobj = _all[j];
							_attachEvent(_all[j], 'blur', function (){cobj.className = trim(cobj.className.replace(' hidefocus', ''));});
							break;
						}
					}
				}
				if(focused) {
					break;
				}
			}
		} catch (e) {
		}
	}
}
var delayShowST = null;
function delayShow(ctrlObj, call, time) {
	if(typeof ctrlObj == 'object') {
		var ctrlid = ctrlObj.id;
		call = call || function () { showMenu(ctrlid); };
	}
	var time = isUndefined(time) ? 500 : time;
	delayShowST = setTimeout(function () {
		if(typeof call == 'function') {
			call();
		} else {
			eval(call);
		}
	}, time);
	if(!ctrlObj.delayinit) {
		_attachEvent(ctrlObj, 'mouseout', function() {clearTimeout(delayShowST);});
		ctrlObj.delayinit = 1;
	}
}

var dragMenuDisabled = false;
function dragMenu(menuObj, e, op) {
	e = e ? e : window.event;
	if(op == 1) {
		if(dragMenuDisabled || in_array(e.target ? e.target.tagName : e.srcElement.tagName, ['TEXTAREA', 'INPUT', 'BUTTON', 'SELECT'])) {
			return;
		}
		JSMENU['drag'] = [e.clientX, e.clientY];
		JSMENU['drag'][2] = parseInt(menuObj.style.left);
		JSMENU['drag'][3] = parseInt(menuObj.style.top);
		document.onmousemove = function(e) {try{dragMenu(menuObj, e, 2);}catch(err){}};
		document.onmouseup = function(e) {try{dragMenu(menuObj, e, 3);}catch(err){}};
		doane(e);
	}else if(op == 2 && JSMENU['drag'][0]) {
		var menudragnow = [e.clientX, e.clientY];
		menuObj.style.left = (JSMENU['drag'][2] + menudragnow[0] - JSMENU['drag'][0]) + 'px';
		menuObj.style.top = (JSMENU['drag'][3] + menudragnow[1] - JSMENU['drag'][1]) + 'px';
		menuObj.removeAttribute('top_');menuObj.removeAttribute('left_');
		doane(e);
	}else if(op == 3) {
		JSMENU['drag'] = [];
		document.onmousemove = null;
		document.onmouseup = null;
	}
}
function setMenuPosition(showid, menuid, pos) {
	var showObj = $(showid);
	var menuObj = menuid ? $(menuid) : $(showid + '_menu');
	if(isUndefined(pos) || !pos) pos = '43';
	var basePoint = parseInt(pos.substr(0, 1));
	var direction = parseInt(pos.substr(1, 1));
	var important = pos.indexOf('!') != -1 ? 1 : 0;
	var sxy = 0, sx = 0, sy = 0, sw = 0, sh = 0, ml = 0, mt = 0, mw = 0, mcw = 0, mh = 0, mch = 0, bpl = 0, bpt = 0;

	if(!menuObj || (basePoint > 0 && !showObj)) return;
	if(showObj) {
		sxy = fetchOffset(showObj);
		sx = sxy['left'];
		sy = sxy['top'];
		sw = showObj.offsetWidth;
		sh = showObj.offsetHeight;
	}
	mw = menuObj.offsetWidth;
	mcw = menuObj.clientWidth;
	mh = menuObj.offsetHeight;
	mch = menuObj.clientHeight;

	switch(basePoint) {
		case 1:
			bpl = sx;
			bpt = sy;
			break;
		case 2:
			bpl = sx + sw;
			bpt = sy;
			break;
		case 3:
			bpl = sx + sw;
			bpt = sy + sh;
			break;
		case 4:
			bpl = sx;
			bpt = sy + sh;
			break;
	}
	switch(direction) {
		case 0:
			menuObj.style.left = (document.body.clientWidth - menuObj.clientWidth) / 2 + 'px';
			mt = (document.documentElement.clientHeight - menuObj.clientHeight) / 2;
			break;
		case 1:
			ml = bpl - mw;
			mt = bpt - mh;
			break;
		case 2:
			ml = bpl;
			mt = bpt - mh;
			break;
		case 3:
			ml = bpl;
			mt = bpt;
			break;
		case 4:
			ml = bpl - mw;
			mt = bpt;
			break;
	}
	var scrollTop = Math.max(document.documentElement.scrollTop, document.body.scrollTop);
	var scrollLeft = Math.max(document.documentElement.scrollLeft, document.body.scrollLeft);
	if(!important) {
		if(in_array(direction, [1, 4]) && ml < 0) {
			ml = bpl;
			if(in_array(basePoint, [1, 4])) ml += sw;
		} else if(ml + mw > scrollLeft + document.body.clientWidth && sx >= mw) {
			ml = bpl - mw;
			if(in_array(basePoint, [2, 3])) {
				ml -= sw;
			} else if(basePoint == 4) {
				ml += sw;
			}
		}
		if(in_array(direction, [1, 2]) && mt < 0) {
			mt = bpt;
			if(in_array(basePoint, [1, 2])) mt += sh;
		} else if(mt + mh > scrollTop + document.documentElement.clientHeight && sy >= mh) {
			mt = bpt - mh;
			if(in_array(basePoint, [3, 4])) mt -= sh;
		}
	}
	if(pos.substr(0, 3) == '210') {
		ml += 69 - sw / 2;
		mt -= 5;
		if(showObj.tagName == 'TEXTAREA') {
			ml -= sw / 2;
			mt += sh / 2;
		}
	}
	if(direction == 0 || menuObj.scrolly) {
		if(BROWSER.ie && BROWSER.ie < 7) {
			if(direction == 0) mt += scrollTop;
		} else {
			if(menuObj.scrolly) mt -= scrollTop;
			menuObj.style.position = 'fixed';
		}
	}
	if(ml) menuObj.style.left = ml + 'px';
	if(mt) menuObj.style.top = mt + 'px';
	if(direction == 0 && BROWSER.ie && !document.documentElement.clientHeight) {
		menuObj.style.position = 'absolute';
		menuObj.style.top = (document.body.clientHeight - menuObj.clientHeight) / 2 + 'px';
	}
	if(menuObj.style.clip && !BROWSER.opera) {
		menuObj.style.clip = 'rect(auto, auto, auto, auto)';
	}
}

function hideMenu(attr, mtype) {
	attr = isUndefined(attr) ? '' : attr;
	mtype = isUndefined(mtype) ? 'menu' : mtype;
	if(attr == '') {
		for(var i = 1; i <= JSMENU['layer']; i++) {
			hideMenu(i, mtype);
		}
		return;
	} else if(typeof attr == 'number') {
		for(var j in JSMENU['active'][attr]) {
			hideMenu(JSMENU['active'][attr][j], mtype);
		}
		return;
	}else if(typeof attr == 'string') {
		var menuObj = $(attr);
		if(!menuObj || (mtype && menuObj.mtype != mtype)) return;
		var ctrlObj = '', ctrlclass = '';
		if((ctrlObj = $(menuObj.getAttribute('ctrlid'))) && (ctrlclass = menuObj.getAttribute('ctrlclass'))) {
			var reg = new RegExp(' ' + ctrlclass);
			ctrlObj.className = ctrlObj.className.replace(reg, '');
		}
		clearTimeout(JSMENU['timer'][attr]);
		var hide = function() {
			if(menuObj.cache) {
				if(menuObj.style.visibility != 'hidden') {
					menuObj.style.display = 'none';
					if(menuObj.cover) $(attr + '_cover').style.display = 'none';
				}
			}else {
				menuObj.parentNode.removeChild(menuObj);
				if(menuObj.cover) $(attr + '_cover').parentNode.removeChild($(attr + '_cover'));
			}
			var tmp = [];
			for(var k in JSMENU['active'][menuObj.layer]) {
				if(attr != JSMENU['active'][menuObj.layer][k]) tmp.push(JSMENU['active'][menuObj.layer][k]);
			}
			JSMENU['active'][menuObj.layer] = tmp;
		};
		if(menuObj.fade) {
			var O = 100;
			var fadeOut = function(O) {
				if(O == 0) {
					clearTimeout(fadeOutTimer);
					hide();
					return;
				}
				menuObj.style.filter = 'progid:DXImageTransform.Microsoft.Alpha(opacity=' + O + ')';
				menuObj.style.opacity = O / 100;
				O -= 20;
				var fadeOutTimer = setTimeout(function () {
					fadeOut(O);
				}, 40);
			};
			fadeOut(O);
		} else {
			hide();
		}
	}
}

function getCurrentStyle(obj, cssproperty, csspropertyNS) {
	if(obj.style[cssproperty]){
		return obj.style[cssproperty];
	}
	if (obj.currentStyle) {
		return obj.currentStyle[cssproperty];
	} else if (document.defaultView.getComputedStyle(obj, null)) {
		var currentStyle = document.defaultView.getComputedStyle(obj, null);
		var value = currentStyle.getPropertyValue(csspropertyNS);
		if(!value){
			value = currentStyle[cssproperty];
		}
		return value;
	} else if (window.getComputedStyle) {
		var currentStyle = window.getComputedStyle(obj, "");
		return currentStyle.getPropertyValue(csspropertyNS);
	}
}

function fetchOffset(obj, mode) {
	var left_offset = 0, top_offset = 0, mode = !mode ? 0 : mode;

	if(obj.getBoundingClientRect && !mode) {
		var rect = obj.getBoundingClientRect();
		var scrollTop = Math.max(document.documentElement.scrollTop, document.body.scrollTop);
		var scrollLeft = Math.max(document.documentElement.scrollLeft, document.body.scrollLeft);
		if(document.documentElement.dir == 'rtl') {
			scrollLeft = scrollLeft + document.documentElement.clientWidth - document.documentElement.scrollWidth;
		}
		left_offset = rect.left + scrollLeft - document.documentElement.clientLeft;
		top_offset = rect.top + scrollTop - document.documentElement.clientTop;
	}
	if(left_offset <= 0 || top_offset <= 0) {
		left_offset = obj.offsetLeft;
		top_offset = obj.offsetTop;
		while((obj = obj.offsetParent) != null) {
			position = getCurrentStyle(obj, 'position', 'position');
			if(position == 'relative') {
				continue;
			}
			left_offset += obj.offsetLeft;
			top_offset += obj.offsetTop;
		}
	}
	return {'left' : left_offset, 'top' : top_offset};
}


function showWindow(k, url, mode, cache, menuv) {
	mode = isUndefined(mode) ? 'get' : mode;
	cache = isUndefined(cache) ? 1 : cache;
	var menuid = 'fwin_' + k;
	var menuObj = $(menuid);
	var drag = null;
	var loadingst = null;
	var hidedom = '';

	if(disallowfloat && disallowfloat.indexOf(k) != -1) {
		if(BROWSER.ie) url += (url.indexOf('?') != -1 ?  '&' : '?') + 'referer=' + escape(location.href);
		location.href = url;
		doane();
		return;
	}

	var fetchContent = function() {
		if(mode == 'get') {
			menuObj.url = url;
			url += (url.search(/\?/) > 0 ? '&' : '?') + 'infloat=yes&handlekey=' + k;
			url += cache == -1 ? '&t='+(+ new Date()) : '';
			if(BROWSER.ie &&  url.indexOf('referer=') < 0) {
				url = url + '&referer=' + encodeURIComponent(location);
			}
			ajaxget(url, 'fwin_content_' + k, null, '', '', function() {initMenu();show();});
		} else if(mode == 'post') {
			menuObj.act = $(url).action;
			ajaxpost(url, 'fwin_content_' + k, '', '', '', function() {initMenu();show();});
		}
		if(parseInt(BROWSER.ie) != 6) {
			loadingst = setTimeout(function() {showDialog('', 'info', '<img src="'+ IMGDIR + '/loading.gif"> ÇëÉÔºò...')}, 500);
		}
	};
	var initMenu = function() {
		clearTimeout(loadingst);
		var objs = menuObj.getElementsByTagName('*');
		var fctrlidinit = false;
		for(var i = 0; i < objs.length; i++) {
			if(objs[i].id) {
				objs[i].setAttribute('fwin', k);
			}
			if(objs[i].className == 'flb' && !fctrlidinit) {
				if(!objs[i].id) objs[i].id = 'fctrl_' + k;
				drag = objs[i].id;
				fctrlidinit = true;
			}
		}
	};
	var show = function() {
		hideMenu('fwin_dialog', 'dialog');
		v = {'mtype':'win','menuid':menuid,'duration':3,'pos':'00','zindex':JSMENU['zIndex']['win'],'drag':typeof drag == null ? '' : drag,'cache':cache};
		for(k in menuv) {
			v[k] = menuv[k];
		}
		showMenu(v);
	};

	if(!menuObj) {
		menuObj = document.createElement('div');
		menuObj.id = menuid;
		menuObj.className = 'fwinmask';
		menuObj.style.display = 'none';
		$('append_parent').appendChild(menuObj);
		evt = ' style="cursor:move" onmousedown="dragMenu($(\'' + menuid + '\'), event, 1)" ondblclick="hideWindow(\'' + k + '\')"';
		if(!BROWSER.ie) {
			hidedom = '<style type="text/css">object{visibility:hidden;}</style>';
		}
		menuObj.innerHTML = hidedom + '<table cellpadding="0" cellspacing="0" class="fwin"><tr><td class="t_l"></td><td class="t_c"' + evt + '></td><td class="t_r"></td></tr><tr><td class="m_l"' + evt + ')">&nbsp;&nbsp;</td><td class="m_c" id="fwin_content_' + k + '">'
			+ '</td><td class="m_r"' + evt + '"></td></tr><tr><td class="b_l"></td><td class="b_c"' + evt + '></td><td class="b_r"></td></tr></table>';
		if(mode == 'html') {
			$('fwin_content_' + k).innerHTML = url;
			initMenu();
			show();
		} else {
			fetchContent();
		}
	} else if((mode == 'get' && (url != menuObj.url || cache != 1)) || (mode == 'post' && $(url).action != menuObj.act)) {
		fetchContent();
	} else {
		show();
	}
	doane();
}


function AC_FL_RunContent() {
	var str = '';
	var ret = AC_GetArgs(arguments, "clsid:d27cdb6e-ae6d-11cf-96b8-444553540000", "application/x-shockwave-flash");
	if(BROWSER.ie && !BROWSER.opera) {
		str += '<object ';
		for (var i in ret.objAttrs) {
			str += i + '="' + ret.objAttrs[i] + '" ';
		}
		str += '>';
		for (var i in ret.params) {
			str += '<param name="' + i + '" value="' + ret.params[i] + '" /> ';
		}
		str += '</object>';
	} else {
		str += '<embed ';
		for (var i in ret.embedAttrs) {
			str += i + '="' + ret.embedAttrs[i] + '" ';
		}
		str += '></embed>';
	}
	return str;
}


function thumbImg(obj, method) {
	if(!obj) {
		return;
	}
	obj.onload = null;
	file = obj.src;
	zw = obj.offsetWidth;
	zh = obj.offsetHeight;
	if(zw < 2) {
		if(!obj.id) {
			obj.id = 'img_' + Math.random();
		}
		setTimeout("thumbImg($('" + obj.id + "'), " + method + ")", 100);
		return;
	}
	zr = zw / zh;
	method = !method ? 0 : 1;
	if(method) {
		fixw = obj.getAttribute('_width');
		fixh = obj.getAttribute('_height');
		if(zw > fixw) {
			zw = fixw;
			zh = zw / zr;
		}
		if(zh > fixh) {
			zh = fixh;
			zw = zh * zr;
		}
	} else {
		fixw = typeof imagemaxwidth == 'undefined' ? '600' : imagemaxwidth;
		if(zw > fixw) {
			zw = fixw;
			zh = zw / zr;
			obj.style.cursor = 'pointer';
			if(!obj.onclick) {
				obj.onclick = function() {
					zoom(obj, obj.src);
				};
			}
		}
	}
	obj.width = zw;
	obj.height = zh;
}

var zoomstatus = 1;
function zoom(obj, zimg, nocover, pn, showexif) {
	$F('_zoom', arguments);
}
```
