## Supplementary material for "Chromatin interaction analysis with updated ChIA-PET Tool (V3)": common_extra.js

```
/*
	[Discuz!] (C)2001-2099 Comsenz Inc.
	This is NOT a freeware, use is subject to license terms

	$Id: common_extra.js 31722 2012-09-25 03:22:12Z zhangguosheng $
*/

function _relatedlinks(rlinkmsgid) {
	if(!$(rlinkmsgid) || $(rlinkmsgid).innerHTML.match(/<script[^\>]*?>/i)) {
		return;
	}
	var alink = new Array(), ignore = new Array();
	var i = 0;
	var msg = $(rlinkmsgid).innerHTML;
	msg = msg.replace(/(<ignore_js_op\>[\s|\S]*?<\/ignore_js_op\>)/ig, function($1) {
		ignore[i] = $1;
		i++;
		return '#ignore_js_op '+(i - 1)+'#';
	});
	var alink_i = 0;
	msg = msg.replace(/(<a.*?<\/a\>)/ig, function($1) {
		alink[alink_i] = $1;
		alink_i++;
		return '#alink '+(alink_i - 1)+'#';
	});
	var relatedid = new Array();
	msg = msg.replace(/(^|>)([^<]+)(?=<|$)/ig, function($1, $2, $3) {
		for(var j = 0; j < relatedlink.length; j++) {
			if(relatedlink[j] && !relatedid[j]) {
				if(relatedlink[j]['surl'] != '') {
					var ra = '<a href="'+relatedlink[j]['surl']+'" target="_blank" class="relatedlink">'+relatedlink[j]['sname']+'</a>';
						alink[alink_i] = ra;
						ra = '#alink '+alink_i+'#';
						alink_i++;
				} else {
					var ra = '<strong><font color="#FF0000">'+relatedlink[j]['sname']+'</font></strong>';
				}
				var $rtmp = $3;
				$3 = $3.replace(relatedlink[j]['sname'], ra);
				if($3 != $rtmp) {
					relatedid[j] = 1;
				}
			}
		}
		return $2 + $3;
    	});

	for(var k in alink) {
		msg = msg.replace('#alink '+k+'#', alink[k]);
	}

	for(var l in ignore) {
		msg = msg.replace('#ignore_js_op '+l+'#', ignore[l]);
	}
	$(rlinkmsgid).innerHTML = msg;
}

function _updatesecqaa(idhash) {
	if($('secqaa_' + idhash)) {
		$('secqaaverify_' + idhash).value = '';
		if(secST['qaa_' + idhash]) {
			clearTimeout(secST['qaa_' + idhash]);
		}
		$('checksecqaaverify_' + idhash).innerHTML = '<img src="'+ IMGDIR + '/none.gif" width="16" height="16" class="vm" />';
		ajaxget('misc.php?mod=secqaa&action=update&idhash=' + idhash, 'secqaa_' + idhash, null, '', '', function() {
			secST['qaa_' + idhash] = setTimeout(function() {$('secqaa_' + idhash).innerHTML = '<span class="xi2 cur1" onclick="updatesecqaa(\''+idhash+'\')">Ë¢ÐÂÑéÖ¤ÎÊ´ð</span>';}, 180000);
		});
	}
}

function _updateseccode(idhash, play) {
	if(isUndefined(play)) {
		if($('seccode_' + idhash)) {
			$('seccodeverify_' + idhash).value = '';
			if(secST['code_' + idhash]) {
				clearTimeout(secST['code_' + idhash]);
			}
			$('checkseccodeverify_' + idhash).innerHTML = '<img src="'+ IMGDIR + '/none.gif" width="16" height="16" class="vm" />';
			ajaxget('misc.php?mod=seccode&action=update&idhash=' + idhash, 'seccode_' + idhash, null, '', '', function() {
				secST['code_' + idhash] = setTimeout(function() {$('seccode_' + idhash).innerHTML = '<span class="xi2 cur1" onclick="updateseccode(\''+idhash+'\')">Ë¢ÐÂÑéÖ¤Âë</span>';}, 180000);
			});
		}
	} else {
		eval('window.document.seccodeplayer_' + idhash + '.SetVariable("isPlay", "1")');
	}
}

function _checksec(type, idhash, showmsg, recall) {
	var showmsg = !showmsg ? 0 : showmsg;
	var secverify = $('sec' + type + 'verify_' + idhash).value;
	if(!secverify) {
		return;
	}
	var x = new Ajax('XML', 'checksec' + type + 'verify_' + idhash);
	x.loading = '';
	$('checksec' + type + 'verify_' + idhash).innerHTML = '<img src="'+ IMGDIR + '/loading.gif" width="16" height="16" class="vm" />';
	x.get('misc.php?mod=sec' + type + '&action=check&inajax=1&&idhash=' + idhash + '&secverify=' + (BROWSER.ie && document.charset == 'utf-8' ? encodeURIComponent(secverify) : secverify), function(s){
		var obj = $('checksec' + type + 'verify_' + idhash);
		obj.style.display = '';
		if(s.substr(0, 7) == 'succeed') {
			obj.innerHTML = '<img src="'+ IMGDIR + '/check_right.gif" width="16" height="16" class="vm" />';
			if(showmsg) {
				recall(1);
			}
		} else {
			obj.innerHTML = '<img src="'+ IMGDIR + '/check_error.gif" width="16" height="16" class="vm" />';
			if(showmsg) {
				if(type == 'code') {
					showError('ÑéÖ¤Âë´íÎó£¬ÇëÖØÐÂÌîÐ´');
				} else if(type == 'qaa') {
					showError('ÑéÖ¤ÎÊ´ð´íÎó£¬ÇëÖØÐÂÌîÐ´');
				}
				recall(0);
			}
		}
	});
}

function _setDoodle(fid, oid, url, tid, from) {
	if(tid == null) {
		hideWindow(fid);
	} else {
		$(tid).style.display = '';
		$(fid).style.display = 'none';
	}
	var doodleText = '[img]'+url+'[/img]';
	if($(oid) != null) {
		if(from == "editor") {
			insertImage(url);
		} else if(from == "fastpost") {
			seditor_insertunit('fastpost', doodleText);
		} else if(from == "forumeditor") {
			if(wysiwyg) {
				insertText('<img src="' + url + '" border="0" alt="" />', false);
			} else {
				insertText(doodleText, strlen(doodleText), 0);
			}
		} else {
			insertContent(oid, doodleText);
		}
	}
}

function _showdistrict(container, elems, totallevel, changelevel, containertype) {
	var getdid = function(elem) {
		var op = elem.options[elem.selectedIndex];
		return op['did'] || op.getAttribute('did') || '0';
	};
	var pid = changelevel >= 1 && elems[0] && $(elems[0]) ? getdid($(elems[0])) : 0;
	var cid = changelevel >= 2 && elems[1] && $(elems[1]) ? getdid($(elems[1])) : 0;
	var did = changelevel >= 3 && elems[2] && $(elems[2]) ? getdid($(elems[2])) : 0;
	var coid = changelevel >= 4 && elems[3] && $(elems[3]) ? getdid($(elems[3])) : 0;
	var url = "home.php?mod=misc&ac=ajax&op=district&container="+container+"&containertype="+containertype
		+"&province="+elems[0]+"&city="+elems[1]+"&district="+elems[2]+"&community="+elems[3]
		+"&pid="+pid + "&cid="+cid+"&did="+did+"&coid="+coid+'&level='+totallevel+'&handlekey='+container+'&inajax=1'+(!changelevel ? '&showdefault=1' : '');
	ajaxget(url, container, '');
}

function _copycode(obj) {
	if(!obj) return false;
	if(window.getSelection) {
		var sel = window.getSelection();
		if (sel.setBaseAndExtent) {
			sel.setBaseAndExtent(obj, 0, obj, 1);
		} else {
			var rng = document.createRange();
			rng.selectNodeContents(obj);
			sel.addRange(rng);
		}
	} else {
		var rng = document.body.createTextRange();
		rng.moveToElementText(obj);
		rng.select();
	}
	setCopy(BROWSER.ie ? obj.innerText.replace(/\r\n\r\n/g, '\r\n') : obj.textContent, '´úÂëÒÑ¸´ÖÆµ½¼ôÌù°å');
}

function _setCopy(text, msg){
	if(BROWSER.ie) {
		var r = clipboardData.setData('Text', text);
		if(r) {
			if(msg) {
				showPrompt(null, null, '<span>' + msg + '</span>', 1500);
			}
		} else {
			showDialog('<div class="c"><div style="width: 200px; text-align: center;">¸´ÖÆÊ§°Ü£¬ÇëÑ¡Ôñ¡°ÔÊÐí·ÃÎÊ¡±</div></div>', 'alert');
		}
	} else {
		var msg = '<div class="c"><div style="width: 200px; text-align: center; text-decoration:underline;">µã´Ë¸´ÖÆµ½¼ôÌù°å</div>' +
		AC_FL_RunContent('id', 'clipboardswf', 'name', 'clipboardswf', 'devicefont', 'false', 'width', '200', 'height', '40', 'src', STATICURL + 'image/common/clipboard.swf', 'menu', 'false',  'allowScriptAccess', 'sameDomain', 'swLiveConnect', 'true', 'wmode', 'transparent', 'style' , 'margin-top:-20px') + '</div>';
		showDialog(msg, 'info');
		text = text.replace(/[\xA0]/g, ' ');
		CLIPBOARDSWFDATA = text;
	}
}

function _showselect(obj, inpid, t, rettype) {
	var showselect_row = function (inpid, s, v, notime, rettype) {
		if(v >= 0) {
			if(!rettype) {
				var notime = !notime ? 0 : 1;
				var t = today.getTime();
				t += 86400000 * v;
				var d = new Date();
				d.setTime(t);
				var month = d.getMonth() + 1;
				month = month < 10 ? '0' + month : month;
				var day = d.getDate();
				day = day < 10 ? '0' + day : day;
				var hour = d.getHours();
				hour = hour < 10 ? '0' + hour : hour;
				var minute = d.getMinutes();
				minute = minute < 10 ? '0' + minute : minute;
				return '<a href="javascript:;" onclick="$(\'' + inpid + '\').value = \'' + d.getFullYear() + '-' + month + '-' + day + (!notime ? ' ' + hour + ':' + minute: '') + '\'">' + s + '</a>';
			} else {
				return '<a href="javascript:;" onclick="$(\'' + inpid + '\').value = \'' + v + '\'">' + s + '</a>';
			}
		} else if(v == -1) {
			return '<a href="javascript:;" onclick="$(\'' + inpid + '\').focus()">' + s + '</a>';
		} else if(v == -2) {
			return '<a href="javascript:;" onclick="$(\'' + inpid + '\').onclick()">' + s + '</a>';
		}
	};

	if(!obj.id) {
		var t = !t ? 0 : t;
		var rettype = !rettype ? 0 : rettype;
		obj.id = 'calendarexp_' + Math.random();
		div = document.createElement('div');
		div.id = obj.id + '_menu';
		div.style.display = 'none';
		div.className = 'p_pop';
		$('append_parent').appendChild(div);
		s = '';
		if(!t) {
			s += showselect_row(inpid, 'Ò»Ìì', 1, 0, rettype);
			s += showselect_row(inpid, 'Ò»ÖÜ', 7, 0, rettype);
			s += showselect_row(inpid, 'Ò»¸öÔÂ', 30, 0, rettype);
			s += showselect_row(inpid, 'Èý¸öÔÂ', 90, 0, rettype);
			s += showselect_row(inpid, '×Ô¶¨Òå', -2);
		} else {
			if($(t)) {
				var lis = $(t).getElementsByTagName('LI');
				for(i = 0;i < lis.length;i++) {
					s += '<a href="javascript:;" onclick="$(\'' + inpid + '\').value = this.innerHTML;$(\''+obj.id+'_menu\').style.display=\'none\'">' + lis[i].innerHTML + '</a>';
				}
				s += showselect_row(inpid, '×Ô¶¨Òå', -1);
			} else {
				s += '<a href="javascript:;" onclick="$(\'' + inpid + '\').value = \'0\'">ÓÀ¾Ã</a>';
				s += showselect_row(inpid, '7 Ìì', 7, 1, rettype);
				s += showselect_row(inpid, '14 Ìì', 14, 1, rettype);
				s += showselect_row(inpid, 'Ò»¸öÔÂ', 30, 1, rettype);
				s += showselect_row(inpid, 'Èý¸öÔÂ', 90, 1, rettype);
				s += showselect_row(inpid, '°ëÄê', 182, 1, rettype);
				s += showselect_row(inpid, 'Ò»Äê', 365, 1, rettype);
				s += showselect_row(inpid, '×Ô¶¨Òå', -1);
			}
		}
		$(div.id).innerHTML = s;
	}
	showMenu({'ctrlid':obj.id,'evt':'click'});
	if(BROWSER.ie && BROWSER.ie < 7) {
		doane(event);
	}
}

function _zoom(obj, zimg, nocover, pn, showexif) {
	zimg = !zimg ? obj.src : zimg;
	showexif = !parseInt(showexif) ? 0 : showexif;
	if(!zoomstatus) {
		window.open(zimg, '', '');
		return;
	}
	if(!obj.id) obj.id = 'img_' + Math.random();
	var faid = !obj.getAttribute('aid') ? 0 : obj.getAttribute('aid');
	var menuid = 'imgzoom';
	var menu = $(menuid);
	var zoomid = menuid + '_zoom';
	var imgtitle = !nocover && obj.title ? '<div class="imgzoom_title">' + obj.title + '</div>' +
		(showexif ? '<div id="' + zoomid + '_exif" class="imgzoom_exif" onmouseover="this.className=\'imgzoom_exif imgzoom_exif_hover\'" onmouseout="this.className=\'imgzoom_exif\'"></div>' : '')
		: '';
	var cover = !nocover ? 1 : 0;
	var pn = !pn ? 0 : 1;
	var maxh = (document.documentElement.clientHeight ? document.documentElement.clientHeight : document.body.clientHeight) - 70;
	var loadCheck = function (obj) {
		if(obj.complete) {
			var imgw = loading.width;
			var imgh = loading.height;
			var r = imgw / imgh;
			var w = document.body.clientWidth * 0.95;
			w = imgw > w ? w : imgw;
			var h = w / r;
			if(w < 100 & h < 100) {
				$(menuid + '_waiting').style.display = 'none';
				hideMenu();
				return;
			}
			if(h > maxh) {
				h = maxh;
				w = h * r;
			}
			if($(menuid)) {
				$(menuid).removeAttribute('top_');$(menuid).removeAttribute('left_');
				clearTimeout($(menuid).getAttribute('timer'));
			}
			showimage(zimg, w, h, imgw, imgh);
			if(showexif && faid) {
				var x = new Ajax();
				x.get('forum.php?mod=ajax&action=exif&aid=' + faid + '&inajax=1', function(s, x) {
					if(s) {
						$(zoomid + '_exif').style.display = '';
						$(zoomid + '_exif').innerHTML = s;
					} else {
						$(zoomid + '_exif').style.display = 'none';
					}
				});
			}
		} else {
			setTimeout(function () { loadCheck(loading); }, 100);
		}
	};
	var showloading = function (zimg, pn) {
		if(!pn) {
			if(!$(menuid + '_waiting')) {
				waiting = document.createElement('img');
				waiting.id = menuid + '_waiting';
				waiting.src = IMGDIR + '/imageloading.gif';
				waiting.style.opacity = '0.8';
				waiting.style.filter = 'alpha(opacity=80)';
				waiting.style.position = 'absolute';
				waiting.style.zIndex = '100000';
				$('append_parent').appendChild(waiting);
			}
		}
		$(menuid + '_waiting').style.display = '';
		$(menuid + '_waiting').style.left = (document.body.clientWidth - 42) / 2 + 'px';
		$(menuid + '_waiting').style.top = ((document.documentElement.clientHeight - 42) / 2 + Math.max(document.documentElement.scrollTop, document.body.scrollTop)) + 'px';
		loading = new Image();
		setTimeout(function () { loadCheck(loading); }, 100);
		if(!pn) {
			$(menuid + '_zoomlayer').style.display = 'none';
		}
		loading.src = zimg;
	};
	var adjustpn = function(h) {
		h = h < 90 ? 90 : h;
		if($('zimg_prev')) {
			$('zimg_prev').style.height= parseInt(h) + 'px';
		}
		if($('zimg_next')) {
			$('zimg_next').style.height= parseInt(h) + 'px';
		}
	};
	var showimage = function (zimg, w, h, imgw, imgh) {
		$(menuid + '_waiting').style.display = 'none';
		$(menuid + '_zoomlayer').style.display = '';
		$(menuid + '_img').style.width = 'auto';
		$(menuid + '_img').style.height = 'auto';
		$(menuid).style.width = (w < 300 ? 320 : w + 20) + 'px';
		mheight = h + 63;
		menu.style.height = mheight + 'px';
		$(menuid + '_zoomlayer').style.height = (mheight < 120 ? 120 : mheight) + 'px';
		$(menuid + '_img').innerHTML = '<img id="' + zoomid + '" src="' + zimg + '" width="' + w + '" height="' + h + '" w="' + imgw + '" h="' + imgh + '" />' + imgtitle;
		if($(menuid + '_imglink')) {
			$(menuid + '_imglink').href = zimg;
		}
		setMenuPosition('', menuid, '00');
		adjustpn(h);
	};
	var adjustTimer = 0;
	var adjustTimerCount = 0;
	var wheelDelta = 0;
	var clientX = 0;
	var clientY = 0;
	var adjust = function(e, a) {
		if(BROWSER.ie && BROWSER.ie<7) {
		} else {
			if(adjustTimerCount) {
				adjustTimer = (function(){
					return setTimeout(function () {
						adjustTimerCount++;
						adjust(e);
					}, 20);
					})();
					$(menuid).setAttribute('timer', adjustTimer);
				if(adjustTimerCount > 17) {
					clearTimeout(adjustTimer);
					adjustTimerCount = 0;
					doane();
				}
			} else if(!a) {
				adjustTimerCount = 1;
				if(adjustTimer) {
					clearTimeout(adjustTimer);
					adjust(e, a);
				} else {
					adjust(e, a);
				}
				doane();
			}
		}
		var ele = $(zoomid);
		if(!ele) {
			return;
		}
		var imgw = ele.getAttribute('w');
		var imgh = ele.getAttribute('h');

		if(!a) {
			e = e || window.event;
			try {
				if(e.altKey || e.shiftKey || e.ctrlKey) return;
			} catch (e) {
				e = {'wheelDelta':wheelDelta, 'clientX':clientX, 'clientY':clientY};
			}
			var step = 0;
			if(e.wheelDelta <= 0 || e.detail > 0) {
				if(ele.width - 1 <= 200 || ele.height - 1 <= 200) {
					clearTimeout(adjustTimer);
					adjustTimerCount = 0;
					doane(e);return;
				}
				step = parseInt(imgw/ele.width)-4;
			} else {
				if(ele.width + 1 >= imgw*40) {
					clearTimeout(adjustTimer);
					adjustTimerCount = 0;
					doane(e);return;
				}
				step = 4-parseInt(imgw/ele.width) || 2;
			}
			if(BROWSER.ie && BROWSER.ie<7) { step *= 5;}
			wheelDelta = e.wheelDelta;
			clientX = e.clientX;
			clientY = e.clientY;
			var ratio = 0;
			if(imgw > imgh) {
				ratio = step/ele.height;
				ele.height += step;
				ele.width = imgw*(ele.height/imgh);
			} else if(imgw < imgh) {
				ratio = step/ele.width;
				ele.width += step;
				ele.height = imgh*(ele.width/imgw);
			}
			if(BROWSER.ie && BROWSER.ie<7) {
				setMenuPosition('', menuid, '00');
			} else {
				var menutop = parseFloat(menu.getAttribute('top_') || menu.style.top);
				var menuleft = parseFloat(menu.getAttribute('left_') || menu.style.left);
				var imgY = clientY - menutop - 39;
				var imgX = clientX - menuleft - 10;
				var newTop = (menutop - imgY*ratio) + 'px';
				var newLeft = (menuleft - imgX*ratio) + 'px';
				menu.style.top = newTop;
				menu.style.left = newLeft;
				menu.setAttribute('top_', newTop);
				menu.setAttribute('left_', newLeft);
			}
		} else {
			ele.width = imgw;
			ele.height = imgh;
		}
		menu.style.width = (parseInt(ele.width < 300 ? 300 : parseInt(ele.width)) + 20) + 'px';
		var mheight = (parseInt(ele.height) + 50);
		menu.style.height = mheight + 'px';
		$(menuid + '_zoomlayer').style.height = (mheight < 120 ? 120 : mheight) + 'px';
		adjustpn(ele.height);
		doane(e);
	};
	if(!menu && !pn) {
		menu = document.createElement('div');
		menu.id = menuid;
		if(cover) {
			menu.innerHTML = '<div class="zoominner" id="' + menuid + '_zoomlayer" style="display:none"><p><span class="y"><a id="' + menuid + '_imglink" class="imglink" target="_blank" title="ÔÚÐÂ´°¿Ú´ò¿ª">ÔÚÐÂ´°¿Ú´ò¿ª</a><a id="' + menuid + '_adjust" href="javascipt:;" class="imgadjust" title="Êµ¼Ê´óÐ¡">Êµ¼Ê´óÐ¡</a>' +
				'<a href="javascript:;" onclick="hideMenu()" class="imgclose" title="¹Ø±Õ">¹Ø±Õ</a></span></p>' +
				'<div class="zimg_p" id="' + menuid + '_picpage"></div><div class="hm" id="' + menuid + '_img"></div></div>';
		} else {
			menu.innerHTML = '<div class="popupmenu_popup" id="' + menuid + '_zoomlayer" style="width:auto"><span class="right y"><a href="javascript:;" onclick="hideMenu()" class="flbc" style="width:20px;margin:0 0 2px 0">¹Ø±Õ</a></span><div class="zimg_p" id="' + menuid + '_picpage"></div><div class="hm" id="' + menuid + '_img"></div></div>';
		}
		if(BROWSER.ie || BROWSER.chrome){
			menu.onmousewheel = adjust;
		} else {
			menu.addEventListener('DOMMouseScroll', adjust, false);
		}
		$('append_parent').appendChild(menu);
		if($(menuid + '_adjust')) {
			$(menuid + '_adjust').onclick = function(e) {adjust(e, 1)};
		}
	}
	showloading(zimg, pn);
	picpage = '';
	$(menuid + '_picpage').innerHTML = '';
	if(typeof zoomgroup == 'object' && zoomgroup[obj.id] && typeof aimgcount == 'object' && aimgcount[zoomgroup[obj.id]]) {
		authorimgs = aimgcount[zoomgroup[obj.id]];
		var aid = obj.id.substr(5), authorlength = authorimgs.length, authorcurrent = '';
		if(authorlength > 1) {
			for(i = 0; i < authorlength;i++) {
				if(aid == authorimgs[i]) {
					authorcurrent = i;
				}
			}
			if(authorcurrent !== '') {
				paid = authorcurrent > 0 ? authorimgs[authorcurrent - 1] : authorimgs[authorlength - 1];
				picpage += ' <div id="zimg_prev" onmouseover="dragMenuDisabled=true;this.style.backgroundPosition=\'0 50px\'" onmouseout="dragMenuDisabled=false;this.style.backgroundPosition=\'0 -100px\';" onclick="_zoom_page(\'' + paid + '\', ' + (showexif ? 1 : 0) + ')" class="zimg_prev"><strong>ÉÏÒ»ÕÅ</strong></div> ';
				paid = authorcurrent < authorlength - 1 ? authorimgs[authorcurrent + 1] : authorimgs[0];
				picpage += ' <div id="zimg_next" onmouseover="dragMenuDisabled=true;this.style.backgroundPosition=\'100% 50px\'" onmouseout="dragMenuDisabled=false;this.style.backgroundPosition=\'100% -100px\';" onclick="_zoom_page(\'' + paid + '\', ' + (showexif ? 1 : 0) + ')" class="zimg_next"><strong>ÏÂÒ»ÕÅ</strong></div> ';
			}
			if(picpage) {
				$(menuid + '_picpage').innerHTML = picpage;
			}
		}
	}
	showMenu({'ctrlid':obj.id,'menuid':menuid,'duration':3,'pos':'00','cover':cover,'drag':menuid,'maxh':''});
}

function _zoom_page(paid, showexif) {
	var imagesrc = $('aimg_' + paid).getAttribute('zoomfile') ? $('aimg_' + paid).getAttribute('zoomfile') : $('aimg_' + paid).getAttribute('file');
	zoom($('aimg_' + paid), imagesrc, 0, 1, showexif ? 1 : 0);
}

function _switchTab(prefix, current, total, activeclass) {
	activeclass = !activeclass ? 'a' : activeclass;
	for(var i = 1; i <= total;i++) {
		var classname = ' '+$(prefix + '_' + i).className+' ';
		$(prefix + '_' + i).className = classname.replace(' '+activeclass+' ','').substr(1);
		$(prefix + '_c_' + i).style.display = 'none';
	}
	$(prefix + '_' + current).className = $(prefix + '_' + current).className + ' '+activeclass;
	$(prefix + '_c_' + current).style.display = '';
}

function _initTab(frameId, type) {
	if (typeof document['diyform'] == 'object' || $(frameId).className.indexOf('tab') < 0) return false;
	type = type || 'click';
	var tabs = $(frameId+'_title').childNodes;
	var arrTab = [];
	for(var i in tabs) {
		if (tabs[i]['nodeType'] == 1 && tabs[i]['className'].indexOf('move-span') > -1) {
			arrTab.push(tabs[i]);
		}
	}
	var counter = 0;
	var tab = document.createElement('ul');
	tab.className = 'tb cl';
	var len = arrTab.length;
	for(var i = 0;i < len; i++) {
		var tabId = arrTab[i].id;
		if (hasClass(arrTab[i],'frame') || hasClass(arrTab[i],'tab')) {
			var arrColumn = [];
			for (var j in arrTab[i].childNodes) {
				if (typeof arrTab[i].childNodes[j] == 'object' && !hasClass(arrTab[i].childNodes[j],'title')) arrColumn.push(arrTab[i].childNodes[j]);
			}
			var frameContent = document.createElement('div');
			frameContent.id = tabId+'_content';
			frameContent.className = hasClass(arrTab[i],'frame') ? 'content cl '+arrTab[i].className.substr(arrTab[i].className.lastIndexOf(' ')+1) : 'content cl';
			var colLen = arrColumn.length;
			for (var k = 0; k < colLen; k++) {
				frameContent.appendChild(arrColumn[k]);
			}
		} else {
			var frameContent = $(tabId+'_content');
			frameContent = frameContent || document.createElement('div');
		}
		frameContent.style.display = counter ? 'none' : '';
		$(frameId+'_content').appendChild(frameContent);

		var li = document.createElement('li');
		li.id = tabId;
		li.className = counter ? '' : 'a';
		var reg = new RegExp('style=\"(.*?)\"', 'gi');
		var matchs = '', style = '', imgs = '';
		while((matchs = reg.exec(arrTab[i].innerHTML))) {
			if(matchs[1].substr(matchs[1].length,1) != ';') {
				matchs[1] += ';';
			}
			style += matchs[1];
		}
		style = style ? ' style="'+style+'"' : '';
		reg = new RegExp('(<img.*?>)', 'gi');
		while((matchs = reg.exec(arrTab[i].innerHTML))) {
			imgs += matchs[1];
		}
		li.innerHTML = arrTab[i]['innerText'] ? arrTab[i]['innerText'] : arrTab[i]['textContent'];
		var a = arrTab[i].getElementsByTagName('a');
		var href = a && a[0] ? a[0].href : 'javascript:;';
		var onclick = type == 'click' ? ' onclick="return false;"' : '';
		li.innerHTML = '<a href="' + href + '"' + onclick + ' onfocus="this.blur();" ' + style + '>' + imgs + li.innerHTML + '</a>';
		_attachEvent(li, type, switchTabUl);
		tab.appendChild(li);
		$(frameId+'_title').removeChild(arrTab[i]);
		counter++;
	}
	$(frameId+'_title').appendChild(tab);
}

function switchTabUl (e) {
	e = e || window.event;
	var aim = e.target || e.srcElement;
	var tabId = aim.id;
	var parent = aim.parentNode;
	while(parent['nodeName'] != 'UL' && parent['nodeName'] != 'BODY') {
		tabId = parent.id;
		parent = parent.parentNode;
	}
	if(parent['nodeName'] == 'BODY') return false;
	var tabs = parent.childNodes;
	var len2 = tabs.length;
	for(var j = 0; j < len2; j++) {
		tabs[j].className = (tabs[j].id == tabId) ? 'a' : '';
		var content = $(tabs[j].id+'_content');
		if (content) content.style.display = tabs[j].id == tabId ? '' : 'none';
	}
}

function slideshow(el) {
	var obj = this;
	if(!el.id) el.id = Math.random();
	if(typeof slideshow.entities == 'undefined') {
		slideshow.entities = {};
	}
	this.id = el.id;
	if(slideshow.entities[this.id]) return false;
	slideshow.entities[this.id] = this;

	this.slideshows = [];
	this.slidebar = [];
	this.slideother = [];
	this.slidebarup = '';
	this.slidebardown = '';
	this.slidenum = 0;
	this.slidestep = 0;

	this.container = el;
	this.imgs = [];
	this.imgLoad = [];
	this.imgLoaded = 0;
	this.imgWidth = 0;
	this.imgHeight = 0;

	this.getMEvent = function(ele, value) {
		value = !value ? 'mouseover' : value;
		var mevent = !ele ? '' : ele.getAttribute('mevent');
		mevent = (mevent == 'click' || mevent == 'mouseover') ? mevent : value;
		return mevent;
	};
	this.slideshows = $C('slideshow', el);
	this.slideshows = this.slideshows.length>0 ? this.slideshows[0].childNodes : null;
	this.slidebar = $C('slidebar', el);
	this.slidebar = this.slidebar.length>0 ? this.slidebar[0] : null;
	this.barmevent = this.getMEvent(this.slidebar);
	this.slideother = $C('slideother', el);
	this.slidebarup = $C('slidebarup', el);
	this.slidebarup = this.slidebarup.length>0 ? this.slidebarup[0] : null;
	this.barupmevent = this.getMEvent(this.slidebarup, 'click');
	this.slidebardown = $C('slidebardown', el);
	this.slidebardown = this.slidebardown.length>0 ? this.slidebardown[0] : null;
	this.bardownmevent = this.getMEvent(this.slidebardown, 'click');
	this.slidenum = parseInt(this.container.getAttribute('slidenum'));
	this.slidestep = parseInt(this.container.getAttribute('slidestep'));
	this.timestep = parseInt(this.container.getAttribute('timestep'));
	this.timestep = !this.timestep ? 2500 : this.timestep;

	this.index = this.length = 0;
	this.slideshows = !this.slideshows ? filterTextNode(el.childNodes) : filterTextNode(this.slideshows);

	this.length = this.slideshows.length;

	for(i=0; i<this.length; i++) {
		this.slideshows[i].style.display = "none";
		_attachEvent(this.slideshows[i], 'mouseover', function(){obj.stop();});
		_attachEvent(this.slideshows[i], 'mouseout', function(){obj.goon();});
	}

	for(i=0, L=this.slideother.length; i<L; i++) {
		for(var j=0;j<this.slideother[i].childNodes.length;j++) {
			if(this.slideother[i].childNodes[j].nodeType == 1) {
				this.slideother[i].childNodes[j].style.display = "none";
			}
		}
	}

	if(!this.slidebar) {
		if(!this.slidenum && !this.slidestep) {
			this.container.parentNode.style.position = 'relative';
			this.slidebar = document.createElement('div');
			this.slidebar.className = 'slidebar';
			this.slidebar.style.position = 'absolute';
			this.slidebar.style.top = '5px';
			this.slidebar.style.left = '4px';
			this.slidebar.style.display = 'none';
			var html = '<ul>';
			for(var i=0; i<this.length; i++) {
				html += '<li on'+this.barmevent+'="slideshow.entities[' + this.id + '].xactive(' + i + '); return false;">' + (i + 1).toString() + '</li>';
			}
			html += '</ul>';
			this.slidebar.innerHTML = html;
			this.container.parentNode.appendChild(this.slidebar);
			this.controls = this.slidebar.getElementsByTagName('li');
		}
	} else {
		this.controls = filterTextNode(this.slidebar.childNodes);
		for(i=0; i<this.controls.length; i++) {
			if(this.slidebarup == this.controls[i] || this.slidebardown == this.controls[i]) continue;
			_attachEvent(this.controls[i], this.barmevent, function(){slidexactive()});
			_attachEvent(this.controls[i], 'mouseout', function(){obj.goon();});
		}
	}
	if(this.slidebarup) {
		_attachEvent(this.slidebarup, this.barupmevent, function(){slidexactive('up')});
	}
	if(this.slidebardown) {
		_attachEvent(this.slidebardown, this.bardownmevent, function(){slidexactive('down')});
	}
	this.activeByStep = function(index) {
		var showindex = 0,i = 0;
		if(index == 'down') {
			showindex = this.index + 1;
			if(showindex > this.length) {
				this.runRoll();
			} else {
				for (i = 0; i < this.slidestep; i++) {
					if(showindex >= this.length) showindex = 0;
					this.index = this.index - this.slidenum + 1;
					if(this.index < 0) this.index = this.length + this.index;
					this.active(showindex);
					showindex++;
				}
			}
		} else if (index == 'up') {
			var tempindex = this.index;
			showindex = this.index - this.slidenum;
			if(showindex < 0) return false;
			for (i = 0; i < this.slidestep; i++) {
				if(showindex < 0) showindex = this.length - Math.abs(showindex);
				this.active(showindex);
				this.index = tempindex = tempindex - 1;
				if(this.index <0) this.index = this.length - 1;
				showindex--;
			}
		}
		return false;
	};
	this.active = function(index) {
		this.slideshows[this.index].style.display = "none";
		this.slideshows[index].style.display = "block";
		if(this.controls && this.controls.length > 0) {
			this.controls[this.index].className = '';
			this.controls[index].className = 'on';
		}
		for(var i=0,L=this.slideother.length; i<L; i++) {
			this.slideother[i].childNodes[this.index].style.display = "none";
			this.slideother[i].childNodes[index].style.display = "block";
		}
		this.index = index;
	};
	this.xactive = function(index) {
		if(!this.slidenum && !this.slidestep) {
			this.stop();
			if(index == 'down') index = this.index == this.length-1 ? 0 : this.index+1;
			if(index == 'up') index = this.index == 0 ? this.length-1 : this.index-1;
			this.active(index);
		} else {
			this.activeByStep(index);
		}
	};
	this.goon = function() {
		this.stop();
		var curobj = this;
		this.timer = setTimeout(function () {
			curobj.run();
		}, this.timestep);
	};
	this.stop = function() {
		clearTimeout(this.timer);
	};
	this.run = function() {
		var index = this.index + 1 < this.length ? this.index + 1 : 0;
		if(!this.slidenum && !this.slidestep) {
			this.active(index);
		} else {
			this.activeByStep('down');
		}
		var ss = this;
		this.timer = setTimeout(function(){
			ss.run();
		}, this.timestep);
	};

	this.runRoll = function() {
		for(var i = 0; i < this.slidenum; i++) {
			if(this.slideshows[i] && typeof this.slideshows[i].style != 'undefined') this.slideshows[i].style.display = 'block';
			for(var j=0,L=this.slideother.length; j<L; j++) {
				this.slideother[j].childNodes[i].style.display = 'block';
			}
		}
		this.index = this.slidenum - 1;
	};
	var imgs = this.slideshows.length ? this.slideshows[0].parentNode.getElementsByTagName('img') : [];
	for(i=0, L=imgs.length; i<L; i++) {
		this.imgs.push(imgs[i]);
		this.imgLoad.push(new Image());
		this.imgLoad[i].onerror = function (){obj.imgLoaded ++;};
		this.imgLoad[i].src = this.imgs[i].src;
	}

	this.getSize = function () {
		if(this.imgs.length == 0) return false;
		var img = this.imgs[0];
		this.imgWidth = img.width ? parseInt(img.width) : 0;
		this.imgHeight = img.height ? parseInt(img.height) : 0;
		var ele = img.parentNode;
		while ((!this.imgWidth || !this.imgHeight) && !hasClass(ele,'slideshow') && ele != document.body) {
			this.imgWidth = ele.style.width ? parseInt(ele.style.width) : 0;
			this.imgHeight = ele.style.height ? parseInt(ele.style.height) : 0;
			ele = ele.parentNode;
		}
		return true;
	};

	this.getSize();

	this.checkLoad = function () {
		var obj = this;
		this.container.style.display = 'block';
		for(i = 0;i < this.imgs.length;i++) {
			if(this.imgLoad[i].complete && !this.imgLoad[i].status) {
				this.imgLoaded++;
				this.imgLoad[i].status = 1;
			}
		}
		var percentEle = $(this.id+'_percent');
		if(this.imgLoaded < this.imgs.length) {
			if (!percentEle) {
				var dom = document.createElement('div');
				dom.id = this.id+"_percent";
				dom.style.width = this.imgWidth ? this.imgWidth+'px' : '150px';
				dom.style.height = this.imgHeight ? this.imgHeight+'px' : '150px';
				dom.style.lineHeight = this.imgHeight ? this.imgHeight+'px' : '150px';
				dom.style.backgroundColor = '#ccc';
				dom.style.textAlign = 'center';
				dom.style.top = '0';
				dom.style.left = '0';
				dom.style.marginLeft = 'auto';
				dom.style.marginRight = 'auto';
				this.slideshows[0].parentNode.appendChild(dom);
				percentEle = dom;
			}
			el.parentNode.style.position = 'relative';
			percentEle.innerHTML = (parseInt(this.imgLoaded / this.imgs.length * 100)) + '%';
			setTimeout(function () {obj.checkLoad();}, 100);
		} else {
			if (percentEle) percentEle.parentNode.removeChild(percentEle);
			if(this.slidebar) this.slidebar.style.display = '';
			this.index = this.length - 1 < 0 ? 0 : this.length - 1;
			if(this.slideshows.length > 0) {
				if(!this.slidenum || !this.slidestep) {
					this.run();
				} else {
					this.runRoll();
				}
			}
		}
	};
	this.checkLoad();
}

function slidexactive(step) {
	var e = getEvent();
	var aim = e.target || e.srcElement;
	var parent = aim.parentNode;
	var xactivei = null, slideboxid = null,currentslideele = null;
	currentslideele = hasClass(aim, 'slidebarup') || hasClass(aim, 'slidebardown') || hasClass(parent, 'slidebar') ? aim : null;
	while(parent && parent != document.body) {
		if(!currentslideele && hasClass(parent, 'slidebar')) {
			currentslideele = parent;
		}
		if(!currentslideele && (hasClass(parent, 'slidebarup') || hasClass(parent, 'slidebardown'))) {
			currentslideele = parent;
		}
		if(hasClass(parent, 'slidebox')) {
			slideboxid = parent.id;
			break;
		}
		parent = parent.parentNode;
	}
	var slidebar = $C('slidebar', parent);
	var children = slidebar.length == 0 ? [] : filterTextNode(slidebar[0].childNodes);
	if(currentslideele && (hasClass(currentslideele, 'slidebarup') || hasClass(currentslideele, 'slidebardown'))) {
		xactivei = step;
	} else {
		for(var j=0,i=0,L=children.length;i<L;i++){
			if(currentslideele && children[i] == currentslideele) {
				xactivei = j;
				break;
			}
			if(!hasClass(children[i], 'slidebarup') && !hasClass(children[i], 'slidebardown')) j++;
		}
	}
	if(slideboxid != null && xactivei != null) slideshow.entities[slideboxid].xactive(xactivei);
}

function filterTextNode(list) {
	var newlist = [];
	for(var i=0; i<list.length; i++) {
		if (list[i].nodeType == 1) {
			newlist.push(list[i]);
		}
	}
	return newlist;
}

function _runslideshow() {
	var slideshows = $C('slidebox');
	for(var i=0,L=slideshows.length; i<L; i++) {
		new slideshow(slideshows[i]);
	}
}

function _showTip(ctrlobj) {
	if(!ctrlobj.id) {
		ctrlobj.id = 'tip_' + Math.random();
	}
	menuid = ctrlobj.id + '_menu';
	if(!$(menuid)) {
		var div = document.createElement('div');
		div.id = ctrlobj.id + '_menu';
		div.className = 'tip tip_4';
		div.style.display = 'none';
		div.innerHTML = '<div class="tip_horn"></div><div class="tip_c">' + ctrlobj.getAttribute('tip') + '</div>';
		$('append_parent').appendChild(div);
	}
	$(ctrlobj.id).onmouseout = function () { hideMenu('', 'prompt'); };
	showMenu({'mtype':'prompt','ctrlid':ctrlobj.id,'pos':'12!','duration':2,'zindex':JSMENU['zIndex']['prompt']});
}

function _showPrompt(ctrlid, evt, msg, timeout) {
	var menuid = ctrlid ? ctrlid + '_pmenu' : 'ntcwin';
	var duration = timeout ? 0 : 3;
	if($(menuid)) {
		$(menuid).parentNode.removeChild($(menuid));
	}
	var div = document.createElement('div');
	div.id = menuid;
	div.className = ctrlid ? 'tip tip_js' : 'ntcwin';
	div.style.display = 'none';
	$('append_parent').appendChild(div);
	if(ctrlid) {
		msg = '<div id="' + ctrlid + '_prompt"><div class="tip_horn"></div><div class="tip_c">' + msg + '</div>';
	} else {
		msg = '<table cellspacing="0" cellpadding="0" class="popupcredit"><tr><td class="pc_l">&nbsp;</td><td class="pc_c"><div class="pc_inner">' + msg +
			'</td><td class="pc_r">&nbsp;</td></tr></table>';
	}
	div.innerHTML = msg;
	if(ctrlid) {
		if(!timeout) {
			evt = 'click';
		}
		if($(ctrlid)) {
			if($(ctrlid).evt !== false) {
				var prompting = function() {
					showMenu({'mtype':'prompt','ctrlid':ctrlid,'evt':evt,'menuid':menuid,'pos':'210'});
				};
				if(evt == 'click') {
					$(ctrlid).onclick = prompting;
				} else {
					$(ctrlid).onmouseover = prompting;
				}
			}
			showMenu({'mtype':'prompt','ctrlid':ctrlid,'evt':evt,'menuid':menuid,'pos':'210','duration':duration,'timeout':timeout,'zindex':JSMENU['zIndex']['prompt']});
			$(ctrlid).unselectable = false;
		}
	} else {
		showMenu({'mtype':'prompt','pos':'00','menuid':menuid,'duration':duration,'timeout':timeout,'zindex':JSMENU['zIndex']['prompt']});
		$(menuid).style.top = (parseInt($(menuid).style.top) - 100) + 'px';
	}
}
function _showCreditPrompt() {
	var notice = getcookie('creditnotice').split('D');
	var basev = getcookie('creditbase').split('D');
	var creditrule = decodeURI(getcookie('creditrule', 1)).replace(String.fromCharCode(9), ' ');
	if(!discuz_uid || notice.length < 2 || notice[9] != discuz_uid) {
		setcookie('creditnotice', '');
		setcookie('creditrule', '');
		return;
	}
	var creditnames = creditnotice.split(',');
	var creditinfo = [];
	var e;
	for(var i = 0; i < creditnames.length; i++) {
		e = creditnames[i].split('|');
		creditinfo[e[0]] = [e[1], e[2]];
	}
	creditShow(creditinfo, notice, basev, 0, 1, creditrule);
}

function creditShow(creditinfo, notice, basev, bk, first, creditrule) {
	var s = '', check = 0;
	for(i = 1; i <= 8; i++) {
		v = parseInt(Math.abs(parseInt(notice[i])) / 5) + 1;
		if(notice[i] !== '0' && creditinfo[i]) {
			s += '<span>' + creditinfo[i][0] + (notice[i] != 0 ? (notice[i] > 0 ? '<em>+' : '<em class="desc">') + notice[i] + '</em>' : '') + creditinfo[i][1] + '</span>';
		}
		if(notice[i] > 0) {
			notice[i] = parseInt(notice[i]) - v;
			basev[i] = parseInt(basev[i]) + v;
		} else if(notice[i] < 0) {
			notice[i] = parseInt(notice[i]) + v;
			basev[i] = parseInt(basev[i]) - v;
		}
		if($('hcredit_' + i)) {
			$('hcredit_' + i).innerHTML = basev[i];
		}
	}
	for(i = 1; i <= 8; i++) {
		if(notice[i] != 0) {
			check = 1;
		}
	}
	if(!s || first) {
		setcookie('creditnotice', '');
		setcookie('creditbase', '');
		setcookie('creditrule', '');
		if(!s) {
			return;
		}
	}
	if(!$('creditpromptdiv')) {
		showPrompt(null, null, '<div id="creditpromptdiv">' + (creditrule ? '<i>' + creditrule + '</i> ' : '') + s + '</div>', 0);
	} else {
		$('creditpromptdiv').innerHTML = s;
	}
	setTimeout(function () {hideMenu(1, 'prompt');$('append_parent').removeChild($('ntcwin'));}, 1500);
}

function _showColorBox(ctrlid, layer, k, bgcolor) {
	var tag1 = !bgcolor ? 'color' : 'backcolor', tag2 = !bgcolor ? 'forecolor' : 'backcolor';
	if(!$(ctrlid + '_menu')) {
		var menu = document.createElement('div');
		menu.id = ctrlid + '_menu';
		menu.className = 'p_pop colorbox';
		menu.unselectable = true;
		menu.style.display = 'none';
		var coloroptions = ['Black', 'Sienna', 'DarkOliveGreen', 'DarkGreen', 'DarkSlateBlue', 'Navy', 'Indigo', 'DarkSlateGray', 'DarkRed', 'DarkOrange', 'Olive', 'Green', 'Teal', 'Blue', 'SlateGray', 'DimGray', 'Red', 'SandyBrown', 'YellowGreen', 'SeaGreen', 'MediumTurquoise', 'RoyalBlue', 'Purple', 'Gray', 'Magenta', 'Orange', 'Yellow', 'Lime', 'Cyan', 'DeepSkyBlue', 'DarkOrchid', 'Silver', 'Pink', 'Wheat', 'LemonChiffon', 'PaleGreen', 'PaleTurquoise', 'LightBlue', 'Plum', 'White'];
		var colortexts = ['ºÚÉ«', 'ô÷É«', '°µéÏé­ÂÌÉ«', '°µÂÌÉ«', '°µ»ÒÀ¶É«', 'º£¾üÉ«', 'µåÇàÉ«', 'Ä«ÂÌÉ«', '°µºìÉ«', '°µ½Û»ÆÉ«', 'éÏé­É«', 'ÂÌÉ«', 'Ë®Ñ¼É«', 'À¶É«', '»ÒÊ¯É«', '°µ»ÒÉ«', 'ºìÉ«', 'É³ºÖÉ«', '»ÆÂÌÉ«', 'º£ÂÌÉ«', '¼äÂÌ±¦Ê¯', '»Ê¼ÒÀ¶', '×ÏÉ«', '»ÒÉ«', 'ºì×ÏÉ«', '³ÈÉ«', '»ÆÉ«', 'Ëá³ÈÉ«', 'ÇàÉ«', 'ÉîÌìÀ¶É«', '°µ×ÏÉ«', 'ÒøÉ«', '·ÛÉ«', 'Ç³»ÆÉ«', 'ÄûÃÊ³ñÉ«', '²ÔÂÌÉ«', '²Ô±¦Ê¯ÂÌ', 'ÁÁÀ¶É«', 'ÑóÀîÉ«', '°×É«'];
		var str = '';
		for(var i = 0; i < 40; i++) {
			str += '<input type="button" style="background-color: ' + coloroptions[i] + '"' + (typeof setEditorTip == 'function' ? ' onmouseover="setEditorTip(\'' + colortexts[i] + '\')" onmouseout="setEditorTip(\'\')"' : '') + ' onclick="'
			+ (typeof wysiwyg == 'undefined' ? 'seditor_insertunit(\'' + k + '\', \'[' + tag1 + '=' + coloroptions[i] + ']\', \'[/' + tag1 + ']\')' : (ctrlid == editorid + '_tbl_param_4' ? '$(\'' + ctrlid + '\').value=\'' + coloroptions[i] + '\';hideMenu(2)' : 'discuzcode(\'' + tag2 + '\', \'' + coloroptions[i] + '\')'))
			+ '" title="' + colortexts[i] + '" />' + (i < 39 && (i + 1) % 8 == 0 ? '<br />' : '');
		}
		menu.innerHTML = str;
		$('append_parent').appendChild(menu);
	}
	showMenu({'ctrlid':ctrlid,'evt':'click','layer':layer});
}

function _toggle_collapse(objname, noimg, complex, lang) {
	var obj = $(objname);
	if(obj) {
		obj.style.display = obj.style.display == '' ? 'none' : '';
		var collapsed = getcookie('collapse');
		collapsed = updatestring(collapsed, objname, !obj.style.display);
		setcookie('collapse', collapsed, (collapsed ? 2592000 : -2592000));
	}
	if(!noimg) {
		var img = $(objname + '_img');
		if(img.tagName != 'IMG') {
			if(img.className.indexOf('_yes') == -1) {
				img.className = img.className.replace(/_no/, '_yes');
				if(lang) {
					img.innerHTML = lang[0];
				}
			} else {
				img.className = img.className.replace(/_yes/, '_no');
				if(lang) {
					img.innerHTML = lang[1];
				}
			}
		} else {
			img.src = img.src.indexOf('_yes.gif') == -1 ? img.src.replace(/_no\.gif/, '_yes\.gif') : img.src.replace(/_yes\.gif/, '_no\.gif');
		}
		img.blur();
	}
	if(complex) {
		var objc = $(objname + '_c');
		if(objc) {
			objc.className = objc.className == 'umh' ? 'umh umn' : 'umh';
		}
	}

}

function _extstyle(css) {
	if(!$('css_extstyle')) {
		loadcss('extstyle');
	}
	$('css_extstyle').href = css ? css + '/style.css' : STATICURL + 'image/common/extstyle_none.css';
	currentextstyle = css;
	setcookie('extstyle', css, 86400 * 30);
	if($('css_widthauto') && !$('css_widthauto').disabled) {
		CSSLOADED['widthauto'] = 0;
		loadcss('widthauto');
	}
}

function _widthauto(obj) {
	if($('css_widthauto')) {
		CSSLOADED['widthauto'] = 1;
	}
	if(!CSSLOADED['widthauto'] || $('css_widthauto').disabled) {
		if(!CSSLOADED['widthauto']) {
			loadcss('widthauto');
		} else {
			$('css_widthauto').disabled = false;
		}
		HTMLNODE.className += ' widthauto';
		setcookie('widthauto', 1, 86400 * 30);
		obj.innerHTML = 'ÇÐ»»µ½Õ­°æ';
	} else {
		$('css_widthauto').disabled = true;
		HTMLNODE.className = HTMLNODE.className.replace(' widthauto', '');
		setcookie('widthauto', -1, 86400 * 30);
		obj.innerHTML = 'ÇÐ»»µ½¿í°æ';
	}
	hideMenu();
}

function _showCreditmenu() {
	if(!$('extcreditmenu_menu')) {
		menu = document.createElement('div');
		menu.id = 'extcreditmenu_menu';
		menu.style.display = 'none';
		menu.className = 'p_pop';
		menu.innerHTML = '<div class="p_opt"><img src="'+ IMGDIR + '/loading.gif" width="16" height="16" class="vm" /> ÇëÉÔºò...</div>';
		$('append_parent').appendChild(menu);
		ajaxget($('extcreditmenu').href, 'extcreditmenu_menu', 'ajaxwaitid');
	}
	showMenu({'ctrlid':'extcreditmenu','ctrlclass':'a','duration':1});
}

function _showForummenu(fid) {
	if(!$('fjump_menu')) {
		fid = !fid ? 0 : fid;
		menu = document.createElement('div');
		menu.id = 'fjump_menu';
		menu.style.display = 'none';
		menu.className = 'p_pop';
		menu.innerHTML = '<div class="p_opt"><img src="'+ IMGDIR + '/loading.gif" width="16" height="16" class="vm" /> ÇëÉÔºò...</div>';
		$('append_parent').appendChild(menu);
		ajaxget('forum.php?mod=ajax&action=forumjump&jfid=' + fid, 'fjump_menu', 'ajaxwaitid');
	}
	showMenu({'ctrlid':'fjump','ctrlclass':'a','duration':2});
}

function _imageRotate(imgid, direct) {
	var image = $(imgid);
	if(!image.getAttribute('deg')) {
		var deg = 0;
		image.setAttribute('ow', image.width);
		image.setAttribute('oh', image.height);
		if(BROWSER.ie) {
			image.setAttribute('om', parseInt(image.currentStyle.marginBottom));
		}
	} else {
		var deg = parseInt(image.getAttribute('deg'));
	}
	var ow = image.getAttribute('ow');
	var oh = image.getAttribute('oh');
	deg = direct == 1 ? deg - 90 : deg + 90;
	if(deg > 270) {
		deg = 0;
	} else if(deg < 0) {
		deg = 270;
	}
	image.setAttribute('deg', deg);
	if(BROWSER.ie) {
		if(!isNaN(image.getAttribute('om'))) {
			image.style.marginBottom = (image.getAttribute('om') + (BROWSER.ie < 8 ? 0 : (deg == 90 || deg == 270 ? Math.abs(ow - oh) : 0))) + 'px';
		}
		image.style.filter = 'progid:DXImageTransform.Microsoft.BasicImage(rotation=' + (deg / 90) + ')';
	} else {
	        switch(deg) {
			case 90:var cow = oh, coh = ow, cx = 0, cy = -oh;break;
			case 180:var cow = ow, coh = oh, cx = -ow, cy = -oh;break;
			case 270:var cow = oh, coh = ow, cx = -ow, cy = 0;break;
	        }
		var canvas = $(image.getAttribute('canvasid'));
		if(!canvas) {
			var i = document.createElement("canvas");
			i.id = 'canva_' + Math.random();
			image.setAttribute('canvasid', i.id);
			image.parentNode.insertBefore(i, image);
			canvas = $(i.id);
		}
		if(deg) {
			var canvasContext = canvas.getContext('2d');
			canvas.setAttribute('width', cow);
			canvas.setAttribute('height', coh);
			canvasContext.rotate(deg * Math.PI / 180);
			canvasContext.drawImage(image, cx, cy, ow, oh);
			image.style.display = 'none';
			canvas.style.display = '';
		} else {
			image.style.display = '';
			canvas.style.display = 'none';
		}
	}
}

function _createPalette(colorid, id, func) {
	var iframe = "<iframe name=\"c"+colorid+"_frame\" src=\"\" frameborder=\"0\" width=\"210\" height=\"148\" scrolling=\"no\"></iframe>";
	if (!$("c"+colorid+"_menu")) {
		var dom = document.createElement('span');
		dom.id = "c"+colorid+"_menu";
		dom.style.display = 'none';
		dom.innerHTML = iframe;
		$('append_parent').appendChild(dom);
	}
	func = !func ? '' : '|' + func;
	window.frames["c"+colorid+"_frame"].location.href = SITEURL+STATICURL+"image/admincp/getcolor.htm?c"+colorid+"|"+id+func;
	showMenu({'ctrlid':'c'+colorid});
	var iframeid = "c"+colorid+"_menu";
	_attachEvent(window, 'scroll', function(){hideMenu(iframeid);});
}
```
