## Supplementary material for "Chromatin interaction analysis with updated ChIA-PET Tool (V3)": forum_viewthread.js

```
/*
	[Discuz!] (C)2001-2099 Comsenz Inc.
	This is NOT a freeware, use is subject to license terms

	$Id: forum_viewthread.js 28794 2012-03-13 05:39:46Z zhangguosheng $
*/

var replyreload = '', attachimgST = new Array(), zoomgroup = new Array(), zoomgroupinit = new Array();

function attachimggroup(pid) {
	if(!zoomgroupinit[pid]) {
		for(i = 0;i < aimgcount[pid].length;i++) {
			zoomgroup['aimg_' + aimgcount[pid][i]] = pid;
		}
		zoomgroupinit[pid] = true;
	}
}

function attachimgshow(pid, onlyinpost) {
	onlyinpost = !onlyinpost ? false : onlyinpost;
	aimgs = aimgcount[pid];
	aimgcomplete = 0;
	loadingcount = 0;
	for(i = 0;i < aimgs.length;i++) {
		obj = $('aimg_' + aimgs[i]);
		if(!obj) {
			aimgcomplete++;
			continue;
		}
		if(onlyinpost && obj.getAttribute('inpost') || !onlyinpost) {
			if(!obj.status) {
				obj.status = 1;
				if(obj.getAttribute('file')) obj.src = obj.getAttribute('file');
				loadingcount++;
			} else if(obj.status == 1) {
				if(obj.complete) {
					obj.status = 2;
				} else {
					loadingcount++;
				}
			} else if(obj.status == 2) {
				aimgcomplete++;
				if(obj.getAttribute('thumbImg')) {
					thumbImg(obj);
				}
			}
			if(loadingcount >= 10) {
				break;
			}
		}
	}
	if(aimgcomplete < aimgs.length) {
		setTimeout(function () {
			attachimgshow(pid, onlyinpost);
		}, 100);
	}
}
```
