## Supplementary material for "Chromatin interaction analysis with updated ChIA-PET Tool (V3)": jquery.min.js

```
/*! jQuery v1.11.2 | (c) 2005, 2014 jQuery Foundation, Inc. | jquery.org/license */
!function(a,b){"object"==typeof module&&"object"==typeof module.exports?module.exports=a.document?b(a,!0):function(a){if(!a.document)throw new Error("jQuery requires a window with a document");return b(a)}:b(a)}("undefined"!=typeof window?window:this,function(a,b){var c=[],d=c.slice,e=c.concat,f=c.push,g=c.indexOf,h={},i=h.toString,j=h.hasOwnProperty,k={},l="1.11.2",m=function(a,b){return new m.fn.init(a,b)},n=/^[\s\uFEFF\xA0]+|[\s\uFEFF\xA0]+$/g,o=/^-ms-/,p=/-([\da-z])/gi,q=function(a,b){return b.toUpperCase()};m.fn=m.prototype={jquery:l,constructor:m,selector:"",length:0,toArray:function(){return d.call(this)},get:function(a){return null!=a?0>a?this[a+this.length]:this[a]:d.call(this)},pushStack:function(a){var b=m.merge(this.constructor(),a);return b.prevObject=this,b.context=this.context,b},each:function(a,b){return m.each(this,a,b)},map:function(a){return this.pushStack(m.map(this,function(b,c){return a.call(b,c,b)}))},slice:function(){return this.pushStack(d.apply(this,arguments))},first:function(){return this.eq(0)},last:function(){return this.eq(-1)},eq:function(a){var b=this.length,c=+a+(0>a?b:0);return this.pushStack(c>=0&&b>c?[this[c]]:[])},end:function(){return this.prevObject||this.constructor(null)},push:f,sort:c.sort,splice:c.splice},m.extend=m.fn.extend=function(){var a,b,c,d,e,f,g=arguments[0]||{},h=1,i=arguments.length,j=!1;for("boolean"==typeof g&&(j=g,g=arguments[h]||{},h++),"object"==typeof g||m.isFunction(g)||(g={}),h===i&&(g=this,h--);i>h;h++)if(null!=(e=arguments[h]))for(d in e)a=g[d],c=e[d],g!==c&&(j&&c&&(m.isPlainObject(c)||(b=m.isArray(c)))?(b?(b=!1,f=a&&m.isArray(a)?a:[]):f=a&&m.isPlainObject(a)?a:{},g[d]=m.extend(j,f,c)):void 0!==c&&(g[d]=c));return g},m.extend({expando:"jQuery"+(l+Math.random()).replace(/\D/g,""),isReady:!0,error:function(a){throw new Error(a)},noop:function(){},isFunction:function(a){return"function"===m.type(a)},isArray:Array.isArray||function(a){return"array"===m.type(a)},isWindow:function(a){return null!=a&&a==a.window},isNumeric:function(a){return!m.isArray(a)&&a-parseFloat(a)+1>=0},isEmptyObject:function(a){var b;for(b in a)return!1;return!0},isPlainObject:function(a){var b;if(!a||"object"!==m.type(a)||a.nodeType||m.isWindow(a))return!1;try{if(a.constructor&&!j.call(a,"constructor")&&!j.call(a.constructor.prototype,"isPrototypeOf"))return!1}catch(c){return!1}if(k.ownLast)for(b in a)return j.call(a,b);for(b in a);return void 0===b||j.call(a,b)},type:function(a){return null==a?a+"":"object"==typeof a||"function"==typeof a?h[i.call(a)]||"object":typeof a},globalEval:function(b){b&&m.trim(b)&&(a.execScript||function(b){a.eval.call(a,b)})(b)},camelCase:function(a){return a.replace(o,"ms-").replace(p,q)},nodeName:function(a,b){return a.nodeName&&a.nodeName.toLowerCase()===b.toLowerCase()},each:function(a,b,c){var d,e=0,f=a.length,g=r(a);if(c){if(g){for(;f>e;e++)if(d=b.apply(a[e],c),d===!1)break}else for(e in a)if(d=b.apply(a[e],c),d===!1)break}else if(g){for(;f>e;e++)if(d=b.call(a[e],e,a[e]),d===!1)break}else for(e in a)if(d=b.call(a[e],e,a[e]),d===!1)break;return a},trim:function(a){return null==a?"":(a+"").replace(n,"")},makeArray:function(a,b){var c=b||[];return null!=a&&(r(Object(a))?m.merge(c,"string"==typeof a?[a]:a):f.call(c,a)),c},inArray:function(a,b,c){var d;if(b){if(g)return g.call(b,a,c);for(d=b.length,c=c?0>c?Math.max(0,d+c):c:0;d>c;c++)if(c in b&&b[c]===a)return c}return-1},merge:function(a,b){var c=+b.length,d=0,e=a.length;while(c>d)a[e++]=b[d++];if(c!==c)while(void 0!==b[d])a[e++]=b[d++];return a.length=e,a},grep:function(a,b,c){for(var d,e=[],f=0,g=a.length,h=!c;g>f;f++)d=!b(a[f],f),d!==h&&e.push(a[f]);return e},map:function(a,b,c){var d,f=0,g=a.length,h=r(a),i=[];if(h)for(;g>f;f++)d=b(a[f],f,c),null!=d&&i.push(d);else for(f in a)d=b(a[f],f,c),null!=d&&i.push(d);return e.apply([],i)},guid:1,proxy:function(a,b){var c,e,f;return"string"==typeof b&&(f=a[b],b=a,a=f),m.isFunction(a)?(c=d.call(arguments,2),e=function(){return a.apply(b||this,c.concat(d.call(arguments)))},e.guid=a.guid=a.guid||m.guid++,e):void 0},now:function(){return+new Date},support:k}),m.each("Boolean Number String Function Array Date RegExp Object Error".split(" "),function(a,b){h["[object "+b+"]"]=b.toLowerCase()});function r(a){var b=a.length,c=m.type(a);return"function"===c||m.isWindow(a)?!1:1===a.nodeType&&b?!0:"array"===c||0===b||"number"==typeof b&&b>0&&b-1 in a}var s=function(a){var b,c,d,e,f,g,h,i,j,k,l,m,n,o,p,q,r,s,t,u="sizzle"+1*new Date,v=a.document,w=0,x=0,y=hb(),z=hb(),A=hb(),B=function(a,b){return a===b&&(l=!0),0},C=1<<31,D={}.hasOwnProperty,E=[],F=E.pop,G=E.push,H=E.push,I=E.slice,J=function(a,b){for(var c=0,d=a.length;d>c;c++)if(a[c]===b)return c;return-1},K="checked|selected|async|autofocus|autoplay|controls|defer|disabled|hidden|ismap|loop|multiple|open|readonly|required|scoped",L="[\\x20\\t\\r\\n\\f]",M="(?:\\\\.|[\\w-]|[^\\x00-\\xa0])+",N=M.replace("w","w#"),O="\\["+L+"*("+M+")(?:"+L+"*([*^$|!~]?=)"+L+"*(?:'((?:\\\\.|[^\\\\'])*)'|\"((?:\\\\.|[^\\\\\"])*)\"|("+N+"))|)"+L+"*\\]",P=":("+M+")(?:\\((('((?:\\\\.|[^\\\\'])*)'|\"((?:\\\\.|[^\\\\\"])*)\")|((?:\\\\.|[^\\\\()[\\]]|"+O+")*)|.*)\\)|)",Q=new RegExp(L+"+","g"),R=new RegExp("^"+L+"+|((?:^|[^\\\\])(?:\\\\.)*)"+L+"+$","g"),S=new RegExp("^"+L+"*,"+L+"*"),T=new RegExp("^"+L+"*([>+~]|"+L+")"+L+"*"),U=new RegExp("="+L+"*([^\\]'\"]*?)"+L+"*\\]","g"),V=new RegExp(P),W=new RegExp("^"+N+"$"),X={ID:new RegExp("^#("+M+")"),CLASS:new RegExp("^\\.("+M+")"),TAG:new RegExp("^("+M.replace("w","w*")+")"),ATTR:new RegExp("^"+O),PSEUDO:new RegExp("^"+P),CHILD:new RegExp("^:(only|first|last|nth|nth-last)-(child|of-type)(?:\\("+L+"*(even|odd|(([+-]|)(\\d*)n|)"+L+"*(?:([+-]|)"+L+"*(\\d+)|))"+L+"*\\)|)","i"),bool:new RegExp("^(?:"+K+")$","i"),needsContext:new RegExp("^"+L+"*[>+~]|:(even|odd|eq|gt|lt|nth|first|last)(?:\\("+L+"*((?:-\\d)?\\d*)"+L+"*\\)|)(?=[^-]|$)","i")},Y=/^(?:input|select|textarea|button)$/i,Z=/^h\d$/i,$=/^[^{]+\{\s*\[native \w/,_=/^(?:#([\w-]+)|(\w+)|\.([\w-]+))$/,ab=/[+~]/,bb=/'|\\/g,cb=new RegExp("\\\\([\\da-f]{1,6}"+L+"?|("+L+")|.)","ig"),db=function(a,b,c){var d="0x"+b-65536;return d!==d||c?b:0>d?String.fromCharCode(d+65536):String.fromCharCode(d>>10|55296,1023&d|56320)},eb=function(){m()};try{H.apply(E=I.call(v.childNodes),v.childNodes),E[v.childNodes.length].nodeType}catch(fb){H={apply:E.length?function(a,b){G.apply(a,I.call(b))}:function(a,b){var c=a.length,d=0;while(a[c++]=b[d++]);a.length=c-1}}}function gb(a,b,d,e){var f,h,j,k,l,o,r,s,w,x;if((b?b.ownerDocument||b:v)!==n&&m(b),b=b||n,d=d||[],k=b.nodeType,"string"!=typeof a||!a||1!==k&&9!==k&&11!==k)return d;if(!e&&p){if(11!==k&&(f=_.exec(a)))if(j=f[1]){if(9===k){if(h=b.getElementById(j),!h||!h.parentNode)return d;if(h.id===j)return d.push(h),d}else if(b.ownerDocument&&(h=b.ownerDocument.getElementById(j))&&t(b,h)&&h.id===j)return d.push(h),d}else{if(f[2])return H.apply(d,b.getElementsByTagName(a)),d;if((j=f[3])&&c.getElementsByClassName)return H.apply(d,b.getElementsByClassName(j)),d}if(c.qsa&&(!q||!q.test(a))){if(s=r=u,w=b,x=1!==k&&a,1===k&&"object"!==b.nodeName.toLowerCase()){o=g(a),(r=b.getAttribute("id"))?s=r.replace(bb,"\\$&"):b.setAttribute("id",s),s="[id='"+s+"'] ",l=o.length;while(l--)o[l]=s+rb(o[l]);w=ab.test(a)&&pb(b.parentNode)||b,x=o.join(",")}if(x)try{return H.apply(d,w.querySelectorAll(x)),d}catch(y){}finally{r||b.removeAttribute("id")}}}return i(a.replace(R,"$1"),b,d,e)}function hb(){var a=[];function b(c,e){return a.push(c+" ")>d.cacheLength&&delete b[a.shift()],b[c+" "]=e}return b}function ib(a){return a[u]=!0,a}function jb(a){var b=n.createElement("div");try{return!!a(b)}catch(c){return!1}finally{b.parentNode&&b.parentNode.removeChild(b),b=null}}function kb(a,b){var c=a.split("|"),e=a.length;while(e--)d.attrHandle[c[e]]=b}function lb(a,b){var c=b&&a,d=c&&1===a.nodeType&&1===b.nodeType&&(~b.sourceIndex||C)-(~a.sourceIndex||C);if(d)return d;if(c)while(c=c.nextSibling)if(c===b)return-1;return a?1:-1}function mb(a){return function(b){var c=b.nodeName.toLowerCase();return"input"===c&&b.type===a}}function nb(a){return function(b){var c=b.nodeName.toLowerCase();return("input"===c||"button"===c)&&b.type===a}}function ob(a){return ib(function(b){return b=+b,ib(function(c,d){var e,f=a([],c.length,b),g=f.length;while(g--)c[e=f[g]]&&(c[e]=!(d[e]=c[e]))})})}function pb(a){return a&&"undefined"!=typeof a.getElementsByTagName&&a}c=gb.support={},f=gb.isXML=function(a){var b=a&&(a.ownerDocument||a).documentElement;return b?"HTML"!==b.nodeName:!1},m=gb.setDocument=function(a){var b,e,g=a?a.ownerDocument||a:v;return g!==n&&9===g.nodeType&&g.documentElement?(n=g,o=g.documentElement,e=g.defaultView,e&&e!==e.top&&(e.addEventListener?e.addEventListener("unload",eb,!1):e.attachEvent&&e.attachEvent("onunload",eb)),p=!f(g),c.attributes=jb(function(a){return a.className="i",!a.getAttribute("className")}),c.getElementsByTagName=jb(function(a){return a.appendChild(g.createComment("")),!a.getElementsByTagName("*").length}),c.getElementsByClassName=$.test(g.getElementsByClassName),c.getById=jb(function(a){return o.appendChild(a).id=u,!g.getElementsByName||!g.getElementsByName(u).length}),c.getById?(d.find.ID=function(a,b){if("undefined"!=typeof b.getElementById&&p){var c=b.getElementById(a);return c&&c.parentNode?[c]:[]}},d.filter.ID=function(a){var b=a.replace(cb,db);return function(a){return a.getAttribute("id")===b}}):(delete d.find.ID,d.filter.ID=function(a){var b=a.replace(cb,db);return function(a){var c="undefined"!=typeof a.getAttributeNode&&a.getAttributeNode("id");return c&&c.value===b}}),d.find.TAG=c.getElementsByTagName?function(a,b){return"undefined"!=typeof b.getElementsByTagName?b.getElementsByTagName(a):c.qsa?b.querySelectorAll(a):void 0}:function(a,b){var c,d=[],e=0,f=b.getElementsByTagName(a);if("*"===a){while(c=f[e++])1===c.nodeType&&d.push(c);return d}return f},d.find.CLASS=c.getElementsByClassName&&function(a,b){return p?b.getElementsByClassName(a):void 0},r=[],q=[],(c.qsa=$.test(g.querySelectorAll))&&(jb(function(a){o.appendChild(a).innerHTML="<a id='"+u+"'></a><select id='"+u+"-\f]' msallowcapture=''><option selected=''></option></select>",a.querySelectorAll("[msallowcapture^='']").length&&q.push("[*^$]="+L+"*(?:''|\"\")"),a.querySelectorAll("[selected]").length||q.push("\\["+L+"*(?:value|"+K+")"),a.querySelectorAll("[id~="+u+"-]").length||q.push("~="),a.querySelectorAll(":checked").length||q.push(":checked"),a.querySelectorAll("a#"+u+"+*").length||q.push(".#.+[+~]")}),jb(function(a){var b=g.createElement("input");b.setAttribute("type","hidden"),a.appendChild(b).setAttribute("name","D"),a.querySelectorAll("[name=d]").length&&q.push("name"+L+"*[*^$|!~]?="),a.querySelectorAll(":enabled").length||q.push(":enabled",":disabled"),a.querySelectorAll("*,:x"),q.push(",.*:")})),(c.matchesSelector=$.test(s=o.matches||o.webkitMatchesSelector||o.mozMatchesSelector||o.oMatchesSelector||o.msMatchesSelector))&&jb(function(a){c.disconnectedMatch=s.call(a,"div"),s.call(a,"[s!='']:x"),r.push("!=",P)}),q=q.length&&new RegExp(q.join("|")),r=r.length&&new RegExp(r.join("|")),b=$.test(o.compareDocumentPosition),t=b||$.test(o.contains)?function(a,b){var c=9===a.nodeType?a.documentElement:a,d=b&&b.parentNode;return a===d||!(!d||1!==d.nodeType||!(c.contains?c.contains(d):a.compareDocumentPosition&&16&a.compareDocumentPosition(d)))}:function(a,b){if(b)while(b=b.parentNode)if(b===a)return!0;return!1},B=b?function(a,b){if(a===b)return l=!0,0;var d=!a.compareDocumentPosition-!b.compareDocumentPosition;return d?d:(d=(a.ownerDocument||a)===(b.ownerDocument||b)?a.compareDocumentPosition(b):1,1&d||!c.sortDetached&&b.compareDocumentPosition(a)===d?a===g||a.ownerDocument===v&&t(v,a)?-1:b===g||b.ownerDocument===v&&t(v,b)?1:k?J(k,a)-J(k,b):0:4&d?-1:1)}:function(a,b){if(a===b)return l=!0,0;var c,d=0,e=a.parentNode,f=b.parentNode,h=[a],i=[b];if(!e||!f)return a===g?-1:b===g?1:e?-1:f?1:k?J(k,a)-J(k,b):0;if(e===f)return lb(a,b);c=a;while(c=c.parentNode)h.unshift(c);c=b;while(c=c.parentNode)i.unshift(c);while(h[d]===i[d])d++;return d?lb(h[d],i[d]):h[d]===v?-1:i[d]===v?1:0},g):n},gb.matches=function(a,b){return gb(a,null,null,b)},gb.matchesSelector=function(a,b){if((a.ownerDocument||a)!==n&&m(a),b=b.replace(U,"='$1']"),!(!c.matchesSelector||!p||r&&r.test(b)||q&&q.test(b)))try{var d=s.call(a,b);if(d||c.disconnectedMatch||a.document&&11!==a.document.nodeType)return d}catch(e){}return gb(b,n,null,[a]).length>0},gb.contains=function(a,b){return(a.ownerDocument||a)!==n&&m(a),t(a,b)},gb.attr=function(a,b){(a.ownerDocument||a)!==n&&m(a);var e=d.attrHandle[b.toLowerCase()],f=e&&D.call(d.attrHandle,b.toLowerCase())?e(a,b,!p):void 0;return void 0!==f?f:c.attributes||!p?a.getAttribute(b):(f=a.getAttributeNode(b))&&f.specified?f.value:null},gb.error=function(a){throw new Error("Syntax error, unrecognized expression: "+a)},gb.uniqueSort=function(a){var b,d=[],e=0,f=0;if(l=!c.detectDuplicates,k=!c.sortStable&&a.slice(0),a.sort(B),l){while(b=a[f++])b===a[f]&&(e=d.push(f));while(e--)a.splice(d[e],1)}return k=null,a},e=gb.getText=function(a){var b,c="",d=0,f=a.nodeType;if(f){if(1===f||9===f||11===f){if("string"==typeof a.textContent)return a.textContent;for(a=a.firstChild;a;a=a.nextSibling)c+=e(a)}else if(3===f||4===f)return a.nodeValue}else while(b=a[d++])c+=e(b);return c},d=gb.selectors={cacheLength:50,createPseudo:ib,match:X,attrHandle:{},find:{},relative:{">":{dir:"parentNode",first:!0}," ":{dir:"parentNode"},"+":{dir:"previousSibling",first:!0},"~":{dir:"previousSibling"}},preFilter:{ATTR:function(a){return a[1]=a[1].replace(cb,db),a[3]=(a[3]||a[4]||a[5]||"").replace(cb,db),"~="===a[2]&&(a[3]=" "+a[3]+" "),a.slice(0,4)},CHILD:function(a){return a[1]=a[1].toLowerCase(),"nth"===a[1].slice(0,3)?(a[3]||gb.error(a[0]),a[4]=+(a[4]?a[5]+(a[6]||1):2*("even"===a[3]||"odd"===a[3])),a[5]=+(a[7]+a[8]||"odd"===a[3])):a[3]&&gb.error(a[0]),a},PSEUDO:function(a){var b,c=!a[6]&&a[2];return X.CHILD.test(a[0])?null:(a[3]?a[2]=a[4]||a[5]||"":c&&V.test(c)&&(b=g(c,!0))&&(b=c.indexOf(")",c.length-b)-c.length)&&(a[0]=a[0].slice(0,b),a[2]=c.slice(0,b)),a.slice(0,3))}},filter:{TAG:function(a){var b=a.replace(cb,db).toLowerCase();return"*"===a?function(){return!0}:function(a){return a.nodeName&&a.nodeName.toLowerCase()===b}},CLASS:function(a){var b=y[a+" "];return b||(b=new RegExp("(^|"+L+")"+a+"("+L+"|$)"))&&y(a,function(a){return b.test("string"==typeof a.className&&a.className||"undefined"!=typeof a.getAttribute&&a.getAttribute("class")||"")})},ATTR:function(a,b,c){return function(d){var e=gb.attr(d,a);return null==e?"!="===b:b?(e+="","="===b?e===c:"!="===b?e!==c:"^="===b?c&&0===e.indexOf(c):"*="===b?c&&e.indexOf(c)>-1:"$="===b?c&&e.slice(-c.length)===c:"~="===b?(" "+e.replace(Q," ")+" ").indexOf(c)>-1:"|="===b?e===c||e.slice(0,c.length+1)===c+"-":!1):!0}},CHILD:function(a,b,c,d,e){var f="nth"!==a.slice(0,3),g="last"!==a.slice(-4),h="of-type"===b;return 1===d&&0===e?function(a){return!!a.parentNode}:function(b,c,i){var j,k,l,m,n,o,p=f!==g?"nextSibling":"previousSibling",q=b.parentNode,r=h&&b.nodeName.toLowerCase(),s=!i&&!h;if(q){if(f){while(p){l=b;while(l=l[p])if(h?l.nodeName.toLowerCase()===r:1===l.nodeType)return!1;o=p="only"===a&&!o&&"nextSibling"}return!0}if(o=[g?q.firstChild:q.lastChild],g&&s){k=q[u]||(q[u]={}),j=k[a]||[],n=j[0]===w&&j[1],m=j[0]===w&&j[2],l=n&&q.childNodes[n];while(l=++n&&l&&l[p]||(m=n=0)||o.pop())if(1===l.nodeType&&++m&&l===b){k[a]=[w,n,m];break}}else if(s&&(j=(b[u]||(b[u]={}))[a])&&j[0]===w)m=j[1];else while(l=++n&&l&&l[p]||(m=n=0)||o.pop())if((h?l.nodeName.toLowerCase()===r:1===l.nodeType)&&++m&&(s&&((l[u]||(l[u]={}))[a]=[w,m]),l===b))break;return m-=e,m===d||m%d===0&&m/d>=0}}},PSEUDO:function(a,b){var c,e=d.pseudos[a]||d.setFilters[a.toLowerCase()]||gb.error("unsupported pseudo: "+a);return e[u]?e(b):e.length>1?(c=[a,a,"",b],d.setFilters.hasOwnProperty(a.toLowerCase())?ib(function(a,c){var d,f=e(a,b),g=f.length;while(g--)d=J(a,f[g]),a[d]=!(c[d]=f[g])}):function(a){return e(a,0,c)}):e}},pseudos:{not:ib(function(a){var b=[],c=[],d=h(a.replace(R,"$1"));return d[u]?ib(function(a,b,c,e){var f,g=d(a,null,e,[]),h=a.length;while(h--)(f=g[h])&&(a[h]=!(b[h]=f))}):function(a,e,f){return b[0]=a,d(b,null,f,c),b[0]=null,!c.pop()}}),has:ib(function(a){return function(b){return gb(a,b).length>0}}),contains:ib(function(a){return a=a.replace(cb,db),function(b){return(b.textContent||b.innerText||e(b)).indexOf(a)>-1}}),lang:ib(function(a){return W.test(a||"")||gb.error("unsupported lang: "+a),a=a.replace(cb,db).toLowerCase(),function(b){var c;do if(c=p?b.lang:b.getAttribute("xml:lang")||b.getAttribute("lang"))return c=c.toLowerCase(),c===a||0===c.indexOf(a+"-");while((b=b.parentNode)&&1===b.nodeType);return!1}}),target:function(b){var c=a.location&&a.location.hash;return c&&c.slice(1)===b.id},root:function(a){return a===o},focus:function(a){return a===n.activeElement&&(!n.hasFocus||n.hasFocus())&&!!(a.type||a.href||~a.tabIndex)},enabled:function(a){return a.disabled===!1},disabled:function(a){return a.disabled===!0},checked:function(a){var b=a.nodeName.toLowerCase();return"input"===b&&!!a.checked||"option"===b&&!!a.selected},selected:function(a){return a.parentNode&&a.parentNode.selectedIndex,a.selected===!0},empty:function(a){for(a=a.firstChild;a;a=a.nextSibling)if(a.nodeType<6)return!1;return!0},parent:function(a){return!d.pseudos.empty(a)},header:function(a){return Z.test(a.nodeName)},input:function(a){return Y.test(a.nodeName)},button:function(a){var b=a.nodeName.toLowerCase();return"input"===b&&"button"===a.type||"button"===b},text:function(a){var b;return"input"===a.nodeName.toLowerCase()&&"text"===a.type&&(null==(b=a.getAttribute("type"))||"text"===b.toLowerCase())},first:ob(function(){return[0]}),last:ob(function(a,b){return[b-1]}),eq:ob(function(a,b,c){return[0>c?c+b:c]}),even:ob(function(a,b){for(var c=0;b>c;c+=2)a.push(c);return a}),odd:ob(function(a,b){for(var c=1;b>c;c+=2)a.push(c);return a}),lt:ob(function(a,b,c){for(var d=0>c?c+b:c;--d>=0;)a.push(d);return a}),gt:ob(function(a,b,c){for(var d=0>c?c+b:c;++d<b;)a.push(d);return a})}},d.pseudos.nth=d.pseudos.eq;for(b in{radio:!0,checkbox:!0,file:!0,password:!0,image:!0})d.pseudos[b]=mb(b);for(b in{submit:!0,reset:!0})d.pseudos[b]=nb(b);function qb(){}qb.prototype=d.filters=d.pseudos,d.setFilters=new qb,g=gb.tokenize=function(a,b){var c,e,f,g,h,i,j,k=z[a+" "];if(k)return b?0:k.slice(0);h=a,i=[],j=d.preFilter;while(h){(!c||(e=S.exec(h)))&&(e&&(h=h.slice(e[0].length)||h),i.push(f=[])),c=!1,(e=T.exec(h))&&(c=e.shift(),f.push({value:c,type:e[0].replace(R," ")}),h=h.slice(c.length));for(g in d.filter)!(e=X[g].exec(h))||j[g]&&!(e=j[g](e))||(c=e.shift(),f.push({value:c,type:g,matches:e}),h=h.slice(c.length));if(!c)break}return b?h.length:h?gb.error(a):z(a,i).slice(0)};function rb(a){for(var b=0,c=a.length,d="";c>b;b++)d+=a[b].value;return d}function sb(a,b,c){var d=b.dir,e=c&&"parentNode"===d,f=x++;return b.first?function(b,c,f){while(b=b[d])if(1===b.nodeType||e)return a(b,c,f)}:function(b,c,g){var h,i,j=[w,f];if(g){while(b=b[d])if((1===b.nodeType||e)&&a(b,c,g))return!0}else while(b=b[d])if(1===b.nodeType||e){if(i=b[u]||(b[u]={}),(h=i[d])&&h[0]===w&&h[1]===f)return j[2]=h[2];if(i[d]=j,j[2]=a(b,c,g))return!0}}}function tb(a){return a.length>1?function(b,c,d){var e=a.length;while(e--)if(!a[e](b,c,d))return!1;return!0}:a[0]}function ub(a,b,c){for(var d=0,e=b.length;e>d;d++)gb(a,b[d],c);return c}function vb(a,b,c,d,e){for(var f,g=[],h=0,i=a.length,j=null!=b;i>h;h++)(f=a[h])&&(!c||c(f,d,e))&&(g.push(f),j&&b.push(h));return g}function wb(a,b,c,d,e,f){return d&&!d[u]&&(d=wb(d)),e&&!e[u]&&(e=wb(e,f)),ib(function(f,g,h,i){var j,k,l,m=[],n=[],o=g.length,p=f||ub(b||"*",h.nodeType?[h]:h,[]),q=!a||!f&&b?p:vb(p,m,a,h,i),r=c?e||(f?a:o||d)?[]:g:q;if(c&&c(q,r,h,i),d){j=vb(r,n),d(j,[],h,i),k=j.length;while(k--)(l=j[k])&&(r[n[k]]=!(q[n[k]]=l))}if(f){if(e||a){if(e){j=[],k=r.length;while(k--)(l=r[k])&&j.push(q[k]=l);e(null,r=[],j,i)}k=r.length;while(k--)(l=r[k])&&(j=e?J(f,l):m[k])>-1&&(f[j]=!(g[j]=l))}}else r=vb(r===g?r.splice(o,r.length):r),e?e(null,g,r,i):H.apply(g,r)})}function xb(a){for(var b,c,e,f=a.length,g=d.relative[a[0].type],h=g||d.relative[" "],i=g?1:0,k=sb(function(a){return a===b},h,!0),l=sb(function(a){return J(b,a)>-1},h,!0),m=[function(a,c,d){var e=!g&&(d||c!==j)||((b=c).nodeType?k(a,c,d):l(a,c,d));return b=null,e}];f>i;i++)if(c=d.relative[a[i].type])m=[sb(tb(m),c)];else{if(c=d.filter[a[i].type].apply(null,a[i].matches),c[u]){for(e=++i;f>e;e++)if(d.relative[a[e].type])break;return wb(i>1&&tb(m),i>1&&rb(a.slice(0,i-1).concat({value:" "===a[i-2].type?"*":""})).replace(R,"$1"),c,e>i&&xb(a.slice(i,e)),f>e&&xb(a=a.slice(e)),f>e&&rb(a))}m.push(c)}return tb(m)}function yb(a,b){var c=b.length>0,e=a.length>0,f=function(f,g,h,i,k){var l,m,o,p=0,q="0",r=f&&[],s=[],t=j,u=f||e&&d.find.TAG("*",k),v=w+=null==t?1:Math.random()||.1,x=u.length;for(k&&(j=g!==n&&g);q!==x&&null!=(l=u[q]);q++){if(e&&l){m=0;while(o=a[m++])if(o(l,g,h)){i.push(l);break}k&&(w=v)}c&&((l=!o&&l)&&p--,f&&r.push(l))}if(p+=q,c&&q!==p){m=0;while(o=b[m++])o(r,s,g,h);if(f){if(p>0)while(q--)r[q]||s[q]||(s[q]=F.call(i));s=vb(s)}H.apply(i,s),k&&!f&&s.length>0&&p+b.length>1&&gb.uniqueSort(i)}return k&&(w=v,j=t),r};return c?ib(f):f}return h=gb.compile=function(a,b){var c,d=[],e=[],f=A[a+" "];if(!f){b||(b=g(a)),c=b.length;while(c--)f=xb(b[c]),f[u]?d.push(f):e.push(f);f=A(a,yb(e,d)),f.selector=a}return f},i=gb.select=function(a,b,e,f){var i,j,k,l,m,n="function"==typeof a&&a,o=!f&&g(a=n.selector||a);if(e=e||[],1===o.length){if(j=o[0]=o[0].slice(0),j.length>2&&"ID"===(k=j[0]).type&&c.getById&&9===b.nodeType&&p&&d.relative[j[1].type]){if(b=(d.find.ID(k.matches[0].replace(cb,db),b)||[])[0],!b)return e;n&&(b=b.parentNode),a=a.slice(j.shift().value.length)}i=X.needsContext.test(a)?0:j.length;while(i--){if(k=j[i],d.relative[l=k.type])break;if((m=d.find[l])&&(f=m(k.matches[0].replace(cb,db),ab.test(j[0].type)&&pb(b.parentNode)||b))){if(j.splice(i,1),a=f.length&&rb(j),!a)return H.apply(e,f),e;break}}}return(n||h(a,o))(f,b,!p,e,ab.test(a)&&pb(b.parentNode)||b),e},c.sortStable=u.split("").sort(B).join("")===u,c.detectDuplicates=!!l,m(),c.sortDetached=jb(function(a){return 1&a.compareDocumentPosition(n.createElement("div"))}),jb(function(a){return a.innerHTML="<a href='#'></a>","#"===a.firstChild.getAttribute("href")})||kb("type|href|height|width",function(a,b,c){return c?void 0:a.getAttribute(b,"type"===b.toLowerCase()?1:2)}),c.attributes&&jb(function(a){return a.innerHTML="<input/>",a.firstChild.setAttribute("value",""),""===a.firstChild.getAttribute("value")})||kb("value",function(a,b,c){return c||"input"!==a.nodeName.toLowerCase()?void 0:a.defaultValue}),jb(function(a){return null==a.getAttribute("disabled")})||kb(K,function(a,b,c){var d;return c?void 0:a[b]===!0?b.toLowerCase():(d=a.getAttributeNode(b))&&d.specified?d.value:null}),gb}(a);m.find=s,m.expr=s.selectors,m.expr[":"]=m.expr.pseudos,m.unique=s.uniqueSort,m.text=s.getText,m.isXMLDoc=s.isXML,m.contains=s.contains;var t=m.expr.match.needsContext,u=/^<(\w+)\s*\/?>(?:<\/\1>|)$/,v=/^.[^:#\[\.,]*$/;function w(a,b,c){if(m.isFunction(b))return m.grep(a,function(a,d){return!!b.call(a,d,a)!==c});if(b.nodeType)return m.grep(a,function(a){return a===b!==c});if("string"==typeof b){if(v.test(b))return m.filter(b,a,c);b=m.filter(b,a)}return m.grep(a,function(a){return m.inArray(a,b)>=0!==c})}m.filter=function(a,b,c){var d=b[0];return c&&(a=":not("+a+")"),1===b.length&&1===d.nodeType?m.find.matchesSelector(d,a)?[d]:[]:m.find.matches(a,m.grep(b,function(a){return 1===a.nodeType}))},m.fn.extend({find:function(a){var b,c=[],d=this,e=d.length;if("string"!=typeof a)return this.pushStack(m(a).filter(function(){for(b=0;e>b;b++)if(m.contains(d[b],this))return!0}));for(b=0;e>b;b++)m.find(a,d[b],c);return c=this.pushStack(e>1?m.unique(c):c),c.selector=this.selector?this.selector+" "+a:a,c},filter:function(a){return this.pushStack(w(this,a||[],!1))},not:function(a){return this.pushStack(w(this,a||[],!0))},is:function(a){return!!w(this,"string"==typeof a&&t.test(a)?m(a):a||[],!1).length}});var x,y=a.document,z=/^(?:\s*(<[\w\W]+>)[^>]*|#([\w-]*))$/,A=m.fn.init=function(a,b){var c,d;if(!a)return this;if("string"==typeof a){if(c="<"===a.charAt(0)&&">"===a.charAt(a.length-1)&&a.length>=3?[null,a,null]:z.exec(a),!c||!c[1]&&b)return!b||b.jquery?(b||x).find(a):this.constructor(b).find(a);if(c[1]){if(b=b instanceof m?b[0]:b,m.merge(this,m.parseHTML(c[1],b&&b.nodeType?b.ownerDocument||b:y,!0)),u.test(c[1])&&m.isPlainObject(b))for(c in b)m.isFunction(this[c])?this[c](b[c]):this.attr(c,b[c]);return this}if(d=y.getElementById(c[2]),d&&d.parentNode){if(d.id!==c[2])return x.find(a);this.length=1,this[0]=d}return this.context=y,this.selector=a,this}return a.nodeType?(this.context=this[0]=a,this.length=1,this):m.isFunction(a)?"undefined"!=typeof x.ready?x.ready(a):a(m):(void 0!==a.selector&&(this.selector=a.selector,this.context=a.context),m.makeArray(a,this))};A.prototype=m.fn,x=m(y);var B=/^(?:parents|prev(?:Until|All))/,C={children:!0,contents:!0,next:!0,prev:!0};m.extend({dir:function(a,b,c){var d=[],e=a[b];while(e&&9!==e.nodeType&&(void 0===c||1!==e.nodeType||!m(e).is(c)))1===e.nodeType&&d.push(e),e=e[b];return d},sibling:function(a,b){for(var c=[];a;a=a.nextSibling)1===a.nodeType&&a!==b&&c.push(a);return c}}),m.fn.extend({has:function(a){var b,c=m(a,this),d=c.length;return this.filter(function(){for(b=0;d>b;b++)if(m.contains(this,c[b]))return!0})},closest:function(a,b){for(var c,d=0,e=this.length,f=[],g=t.test(a)||"string"!=typeof a?m(a,b||this.context):0;e>d;d++)for(c=this[d];c&&c!==b;c=c.parentNode)if(c.nodeType<11&&(g?g.index(c)>-1:1===c.nodeType&&m.find.matchesSelector(c,a))){f.push(c);break}return this.pushStack(f.length>1?m.unique(f):f)},index:function(a){return a?"string"==typeof a?m.inArray(this[0],m(a)):m.inArray(a.jquery?a[0]:a,this):this[0]&&this[0].parentNode?this.first().prevAll().length:-1},add:function(a,b){return this.pushStack(m.unique(m.merge(this.get(),m(a,b))))},addBack:function(a){return this.add(null==a?this.prevObject:this.prevObject.filter(a))}});function D(a,b){do a=a[b];while(a&&1!==a.nodeType);return a}m.each({parent:function(a){var b=a.parentNode;return b&&11!==b.nodeType?b:null},parents:function(a){return m.dir(a,"parentNode")},parentsUntil:function(a,b,c){return m.dir(a,"parentNode",c)},next:function(a){return D(a,"nextSibling")},prev:function(a){return D(a,"previousSibling")},nextAll:function(a){return m.dir(a,"nextSibling")},prevAll:function(a){return m.dir(a,"previousSibling")},nextUntil:function(a,b,c){return m.dir(a,"nextSibling",c)},prevUntil:function(a,b,c){return m.dir(a,"previousSibling",c)},siblings:function(a){return m.sibling((a.parentNode||{}).firstChild,a)},children:function(a){return m.sibling(a.firstChild)},contents:function(a){return m.nodeName(a,"iframe")?a.contentDocument||a.contentWindow.document:m.merge([],a.childNodes)}},function(a,b){m.fn[a]=function(c,d){var e=m.map(this,b,c);return"Until"!==a.slice(-5)&&(d=c),d&&"string"==typeof d&&(e=m.filter(d,e)),this.length>1&&(C[a]||(e=m.unique(e)),B.test(a)&&(e=e.reverse())),this.pushStack(e)}});var E=/\S+/g,F={};function G(a){var b=F[a]={};return m.each(a.match(E)||[],function(a,c){b[c]=!0}),b}m.Callbacks=function(a){a="string"==typeof a?F[a]||G(a):m.extend({},a);var b,c,d,e,f,g,h=[],i=!a.once&&[],j=function(l){for(c=a.memory&&l,d=!0,f=g||0,g=0,e=h.length,b=!0;h&&e>f;f++)if(h[f].apply(l[0],l[1])===!1&&a.stopOnFalse){c=!1;break}b=!1,h&&(i?i.length&&j(i.shift()):c?h=[]:k.disable())},k={add:function(){if(h){var d=h.length;!function f(b){m.each(b,function(b,c){var d=m.type(c);"function"===d?a.unique&&k.has(c)||h.push(c):c&&c.length&&"string"!==d&&f(c)})}(arguments),b?e=h.length:c&&(g=d,j(c))}return this},remove:function(){return h&&m.each(arguments,function(a,c){var d;while((d=m.inArray(c,h,d))>-1)h.splice(d,1),b&&(e>=d&&e--,f>=d&&f--)}),this},has:function(a){return a?m.inArray(a,h)>-1:!(!h||!h.length)},empty:function(){return h=[],e=0,this},disable:function(){return h=i=c=void 0,this},disabled:function(){return!h},lock:function(){return i=void 0,c||k.disable(),this},locked:function(){return!i},fireWith:function(a,c){return!h||d&&!i||(c=c||[],c=[a,c.slice?c.slice():c],b?i.push(c):j(c)),this},fire:function(){return k.fireWith(this,arguments),this},fired:function(){return!!d}};return k},m.extend({Deferred:function(a){var b=[["resolve","done",m.Callbacks("once memory"),"resolved"],["reject","fail",m.Callbacks("once memory"),"rejected"],["notify","progress",m.Callbacks("memory")]],c="pending",d={state:function(){return c},always:function(){return e.done(arguments).fail(arguments),this},then:function(){var a=arguments;return m.Deferred(function(c){m.each(b,function(b,f){var g=m.isFunction(a[b])&&a[b];e[f[1]](function(){var a=g&&g.apply(this,arguments);a&&m.isFunction(a.promise)?a.promise().done(c.resolve).fail(c.reject).progress(c.notify):c[f[0]+"With"](this===d?c.promise():this,g?[a]:arguments)})}),a=null}).promise()},promise:function(a){return null!=a?m.extend(a,d):d}},e={};return d.pipe=d.then,m.each(b,function(a,f){var g=f[2],h=f[3];d[f[1]]=g.add,h&&g.add(function(){c=h},b[1^a][2].disable,b[2][2].lock),e[f[0]]=function(){return e[f[0]+"With"](this===e?d:this,arguments),this},e[f[0]+"With"]=g.fireWith}),d.promise(e),a&&a.call(e,e),e},when:function(a){var b=0,c=d.call(arguments),e=c.length,f=1!==e||a&&m.isFunction(a.promise)?e:0,g=1===f?a:m.Deferred(),h=function(a,b,c){return function(e){b[a]=this,c[a]=arguments.length>1?d.call(arguments):e,c===i?g.notifyWith(b,c):--f||g.resolveWith(b,c)}},i,j,k;if(e>1)for(i=new Array(e),j=new Array(e),k=new Array(e);e>b;b++)c[b]&&m.isFunction(c[b].promise)?c[b].promise().done(h(b,k,c)).fail(g.reject).progress(h(b,j,i)):--f;return f||g.resolveWith(k,c),g.promise()}});var H;m.fn.ready=function(a){return m.ready.promise().done(a),this},m.extend({isReady:!1,readyWait:1,holdReady:function(a){a?m.readyWait++:m.ready(!0)},ready:function(a){if(a===!0?!--m.readyWait:!m.isReady){if(!y.body)return setTimeout(m.ready);m.isReady=!0,a!==!0&&--m.readyWait>0||(H.resolveWith(y,[m]),m.fn.triggerHandler&&(m(y).triggerHandler("ready"),m(y).off("ready")))}}});function I(){y.addEventListener?(y.removeEventListener("DOMContentLoaded",J,!1),a.removeEventListener("load",J,!1)):(y.detachEvent("onreadystatechange",J),a.detachEvent("onload",J))}function J(){(y.addEventListener||"load"===event.type||"complete"===y.readyState)&&(I(),m.ready())}m.ready.promise=function(b){if(!H)if(H=m.Deferred(),"complete"===y.readyState)setTimeout(m.ready);else if(y.addEventListener)y.addEventListener("DOMContentLoaded",J,!1),a.addEventListener("load",J,!1);else{y.attachEvent("onreadystatechange",J),a.attachEvent("onload",J);var c=!1;try{c=null==a.frameElement&&y.documentElement}catch(d){}c&&c.doScroll&&!function e(){if(!m.isReady){try{c.doScroll("left")}catch(a){return setTimeout(e,50)}I(),m.ready()}}()}return H.promise(b)};var K="undefined",L;for(L in m(k))break;k.ownLast="0"!==L,k.inlineBlockNeedsLayout=!1,m(function(){var a,b,c,d;c=y.getElementsByTagName("body")[0],c&&c.style&&(b=y.createElement("div"),d=y.createElement("div"),d.style.cssText="position:absolute;border:0;width:0;height:0;top:0;left:-9999px",c.appendChild(d).appendChild(b),typeof b.style.zoom!==K&&(b.style.cssText="display:inline;margin:0;border:0;padding:1px;width:1px;zoom:1",k.inlineBlockNeedsLayout=a=3===b.offsetWidth,a&&(c.style.zoom=1)),c.removeChild(d))}),function(){var a=y.createElement("div");if(null==k.deleteExpando){k.deleteExpando=!0;try{delete a.test}catch(b){k.deleteExpando=!1}}a=null}(),m.acceptData=function(a){var b=m.noData[(a.nodeName+" ").toLowerCase()],c=+a.nodeType||1;return 1!==c&&9!==c?!1:!b||b!==!0&&a.getAttribute("classid")===b};var M=/^(?:\{[\w\W]*\}|\[[\w\W]*\])$/,N=/([A-Z])/g;function O(a,b,c){if(void 0===c&&1===a.nodeType){var d="data-"+b.replace(N,"-$1").toLowerCase();if(c=a.getAttribute(d),"string"==typeof c){try{c="true"===c?!0:"false"===c?!1:"null"===c?null:+c+""===c?+c:M.test(c)?m.parseJSON(c):c}catch(e){}m.data(a,b,c)}else c=void 0}return c}function P(a){var b;for(b in a)if(("data"!==b||!m.isEmptyObject(a[b]))&&"toJSON"!==b)return!1;
return!0}function Q(a,b,d,e){if(m.acceptData(a)){var f,g,h=m.expando,i=a.nodeType,j=i?m.cache:a,k=i?a[h]:a[h]&&h;if(k&&j[k]&&(e||j[k].data)||void 0!==d||"string"!=typeof b)return k||(k=i?a[h]=c.pop()||m.guid++:h),j[k]||(j[k]=i?{}:{toJSON:m.noop}),("object"==typeof b||"function"==typeof b)&&(e?j[k]=m.extend(j[k],b):j[k].data=m.extend(j[k].data,b)),g=j[k],e||(g.data||(g.data={}),g=g.data),void 0!==d&&(g[m.camelCase(b)]=d),"string"==typeof b?(f=g[b],null==f&&(f=g[m.camelCase(b)])):f=g,f}}function R(a,b,c){if(m.acceptData(a)){var d,e,f=a.nodeType,g=f?m.cache:a,h=f?a[m.expando]:m.expando;if(g[h]){if(b&&(d=c?g[h]:g[h].data)){m.isArray(b)?b=b.concat(m.map(b,m.camelCase)):b in d?b=[b]:(b=m.camelCase(b),b=b in d?[b]:b.split(" ")),e=b.length;while(e--)delete d[b[e]];if(c?!P(d):!m.isEmptyObject(d))return}(c||(delete g[h].data,P(g[h])))&&(f?m.cleanData([a],!0):k.deleteExpando||g!=g.window?delete g[h]:g[h]=null)}}}m.extend({cache:{},noData:{"applet ":!0,"embed ":!0,"object ":"clsid:D27CDB6E-AE6D-11cf-96B8-444553540000"},hasData:function(a){return a=a.nodeType?m.cache[a[m.expando]]:a[m.expando],!!a&&!P(a)},data:function(a,b,c){return Q(a,b,c)},removeData:function(a,b){return R(a,b)},_data:function(a,b,c){return Q(a,b,c,!0)},_removeData:function(a,b){return R(a,b,!0)}}),m.fn.extend({data:function(a,b){var c,d,e,f=this[0],g=f&&f.attributes;if(void 0===a){if(this.length&&(e=m.data(f),1===f.nodeType&&!m._data(f,"parsedAttrs"))){c=g.length;while(c--)g[c]&&(d=g[c].name,0===d.indexOf("data-")&&(d=m.camelCase(d.slice(5)),O(f,d,e[d])));m._data(f,"parsedAttrs",!0)}return e}return"object"==typeof a?this.each(function(){m.data(this,a)}):arguments.length>1?this.each(function(){m.data(this,a,b)}):f?O(f,a,m.data(f,a)):void 0},removeData:function(a){return this.each(function(){m.removeData(this,a)})}}),m.extend({queue:function(a,b,c){var d;return a?(b=(b||"fx")+"queue",d=m._data(a,b),c&&(!d||m.isArray(c)?d=m._data(a,b,m.makeArray(c)):d.push(c)),d||[]):void 0},dequeue:function(a,b){b=b||"fx";var c=m.queue(a,b),d=c.length,e=c.shift(),f=m._queueHooks(a,b),g=function(){m.dequeue(a,b)};"inprogress"===e&&(e=c.shift(),d--),e&&("fx"===b&&c.unshift("inprogress"),delete f.stop,e.call(a,g,f)),!d&&f&&f.empty.fire()},_queueHooks:function(a,b){var c=b+"queueHooks";return m._data(a,c)||m._data(a,c,{empty:m.Callbacks("once memory").add(function(){m._removeData(a,b+"queue"),m._removeData(a,c)})})}}),m.fn.extend({queue:function(a,b){var c=2;return"string"!=typeof a&&(b=a,a="fx",c--),arguments.length<c?m.queue(this[0],a):void 0===b?this:this.each(function(){var c=m.queue(this,a,b);m._queueHooks(this,a),"fx"===a&&"inprogress"!==c[0]&&m.dequeue(this,a)})},dequeue:function(a){return this.each(function(){m.dequeue(this,a)})},clearQueue:function(a){return this.queue(a||"fx",[])},promise:function(a,b){var c,d=1,e=m.Deferred(),f=this,g=this.length,h=function(){--d||e.resolveWith(f,[f])};"string"!=typeof a&&(b=a,a=void 0),a=a||"fx";while(g--)c=m._data(f[g],a+"queueHooks"),c&&c.empty&&(d++,c.empty.add(h));return h(),e.promise(b)}});var S=/[+-]?(?:\d*\.|)\d+(?:[eE][+-]?\d+|)/.source,T=["Top","Right","Bottom","Left"],U=function(a,b){return a=b||a,"none"===m.css(a,"display")||!m.contains(a.ownerDocument,a)},V=m.access=function(a,b,c,d,e,f,g){var h=0,i=a.length,j=null==c;if("object"===m.type(c)){e=!0;for(h in c)m.access(a,b,h,c[h],!0,f,g)}else if(void 0!==d&&(e=!0,m.isFunction(d)||(g=!0),j&&(g?(b.call(a,d),b=null):(j=b,b=function(a,b,c){return j.call(m(a),c)})),b))for(;i>h;h++)b(a[h],c,g?d:d.call(a[h],h,b(a[h],c)));return e?a:j?b.call(a):i?b(a[0],c):f},W=/^(?:checkbox|radio)$/i;!function(){var a=y.createElement("input"),b=y.createElement("div"),c=y.createDocumentFragment();if(b.innerHTML="  <link/><table></table><a href='/a'>a</a><input type='checkbox'/>",k.leadingWhitespace=3===b.firstChild.nodeType,k.tbody=!b.getElementsByTagName("tbody").length,k.htmlSerialize=!!b.getElementsByTagName("link").length,k.html5Clone="<:nav></:nav>"!==y.createElement("nav").cloneNode(!0).outerHTML,a.type="checkbox",a.checked=!0,c.appendChild(a),k.appendChecked=a.checked,b.innerHTML="<textarea>x</textarea>",k.noCloneChecked=!!b.cloneNode(!0).lastChild.defaultValue,c.appendChild(b),b.innerHTML="<input type='radio' checked='checked' name='t'/>",k.checkClone=b.cloneNode(!0).cloneNode(!0).lastChild.checked,k.noCloneEvent=!0,b.attachEvent&&(b.attachEvent("onclick",function(){k.noCloneEvent=!1}),b.cloneNode(!0).click()),null==k.deleteExpando){k.deleteExpando=!0;try{delete b.test}catch(d){k.deleteExpando=!1}}}(),function(){var b,c,d=y.createElement("div");for(b in{submit:!0,change:!0,focusin:!0})c="on"+b,(k[b+"Bubbles"]=c in a)||(d.setAttribute(c,"t"),k[b+"Bubbles"]=d.attributes[c].expando===!1);d=null}();var X=/^(?:input|select|textarea)$/i,Y=/^key/,Z=/^(?:mouse|pointer|contextmenu)|click/,$=/^(?:focusinfocus|focusoutblur)$/,_=/^([^.]*)(?:\.(.+)|)$/;function ab(){return!0}function bb(){return!1}function cb(){try{return y.activeElement}catch(a){}}m.event={global:{},add:function(a,b,c,d,e){var f,g,h,i,j,k,l,n,o,p,q,r=m._data(a);if(r){c.handler&&(i=c,c=i.handler,e=i.selector),c.guid||(c.guid=m.guid++),(g=r.events)||(g=r.events={}),(k=r.handle)||(k=r.handle=function(a){return typeof m===K||a&&m.event.triggered===a.type?void 0:m.event.dispatch.apply(k.elem,arguments)},k.elem=a),b=(b||"").match(E)||[""],h=b.length;while(h--)f=_.exec(b[h])||[],o=q=f[1],p=(f[2]||"").split(".").sort(),o&&(j=m.event.special[o]||{},o=(e?j.delegateType:j.bindType)||o,j=m.event.special[o]||{},l=m.extend({type:o,origType:q,data:d,handler:c,guid:c.guid,selector:e,needsContext:e&&m.expr.match.needsContext.test(e),namespace:p.join(".")},i),(n=g[o])||(n=g[o]=[],n.delegateCount=0,j.setup&&j.setup.call(a,d,p,k)!==!1||(a.addEventListener?a.addEventListener(o,k,!1):a.attachEvent&&a.attachEvent("on"+o,k))),j.add&&(j.add.call(a,l),l.handler.guid||(l.handler.guid=c.guid)),e?n.splice(n.delegateCount++,0,l):n.push(l),m.event.global[o]=!0);a=null}},remove:function(a,b,c,d,e){var f,g,h,i,j,k,l,n,o,p,q,r=m.hasData(a)&&m._data(a);if(r&&(k=r.events)){b=(b||"").match(E)||[""],j=b.length;while(j--)if(h=_.exec(b[j])||[],o=q=h[1],p=(h[2]||"").split(".").sort(),o){l=m.event.special[o]||{},o=(d?l.delegateType:l.bindType)||o,n=k[o]||[],h=h[2]&&new RegExp("(^|\\.)"+p.join("\\.(?:.*\\.|)")+"(\\.|$)"),i=f=n.length;while(f--)g=n[f],!e&&q!==g.origType||c&&c.guid!==g.guid||h&&!h.test(g.namespace)||d&&d!==g.selector&&("**"!==d||!g.selector)||(n.splice(f,1),g.selector&&n.delegateCount--,l.remove&&l.remove.call(a,g));i&&!n.length&&(l.teardown&&l.teardown.call(a,p,r.handle)!==!1||m.removeEvent(a,o,r.handle),delete k[o])}else for(o in k)m.event.remove(a,o+b[j],c,d,!0);m.isEmptyObject(k)&&(delete r.handle,m._removeData(a,"events"))}},trigger:function(b,c,d,e){var f,g,h,i,k,l,n,o=[d||y],p=j.call(b,"type")?b.type:b,q=j.call(b,"namespace")?b.namespace.split("."):[];if(h=l=d=d||y,3!==d.nodeType&&8!==d.nodeType&&!$.test(p+m.event.triggered)&&(p.indexOf(".")>=0&&(q=p.split("."),p=q.shift(),q.sort()),g=p.indexOf(":")<0&&"on"+p,b=b[m.expando]?b:new m.Event(p,"object"==typeof b&&b),b.isTrigger=e?2:3,b.namespace=q.join("."),b.namespace_re=b.namespace?new RegExp("(^|\\.)"+q.join("\\.(?:.*\\.|)")+"(\\.|$)"):null,b.result=void 0,b.target||(b.target=d),c=null==c?[b]:m.makeArray(c,[b]),k=m.event.special[p]||{},e||!k.trigger||k.trigger.apply(d,c)!==!1)){if(!e&&!k.noBubble&&!m.isWindow(d)){for(i=k.delegateType||p,$.test(i+p)||(h=h.parentNode);h;h=h.parentNode)o.push(h),l=h;l===(d.ownerDocument||y)&&o.push(l.defaultView||l.parentWindow||a)}n=0;while((h=o[n++])&&!b.isPropagationStopped())b.type=n>1?i:k.bindType||p,f=(m._data(h,"events")||{})[b.type]&&m._data(h,"handle"),f&&f.apply(h,c),f=g&&h[g],f&&f.apply&&m.acceptData(h)&&(b.result=f.apply(h,c),b.result===!1&&b.preventDefault());if(b.type=p,!e&&!b.isDefaultPrevented()&&(!k._default||k._default.apply(o.pop(),c)===!1)&&m.acceptData(d)&&g&&d[p]&&!m.isWindow(d)){l=d[g],l&&(d[g]=null),m.event.triggered=p;try{d[p]()}catch(r){}m.event.triggered=void 0,l&&(d[g]=l)}return b.result}},dispatch:function(a){a=m.event.fix(a);var b,c,e,f,g,h=[],i=d.call(arguments),j=(m._data(this,"events")||{})[a.type]||[],k=m.event.special[a.type]||{};if(i[0]=a,a.delegateTarget=this,!k.preDispatch||k.preDispatch.call(this,a)!==!1){h=m.event.handlers.call(this,a,j),b=0;while((f=h[b++])&&!a.isPropagationStopped()){a.currentTarget=f.elem,g=0;while((e=f.handlers[g++])&&!a.isImmediatePropagationStopped())(!a.namespace_re||a.namespace_re.test(e.namespace))&&(a.handleObj=e,a.data=e.data,c=((m.event.special[e.origType]||{}).handle||e.handler).apply(f.elem,i),void 0!==c&&(a.result=c)===!1&&(a.preventDefault(),a.stopPropagation()))}return k.postDispatch&&k.postDispatch.call(this,a),a.result}},handlers:function(a,b){var c,d,e,f,g=[],h=b.delegateCount,i=a.target;if(h&&i.nodeType&&(!a.button||"click"!==a.type))for(;i!=this;i=i.parentNode||this)if(1===i.nodeType&&(i.disabled!==!0||"click"!==a.type)){for(e=[],f=0;h>f;f++)d=b[f],c=d.selector+" ",void 0===e[c]&&(e[c]=d.needsContext?m(c,this).index(i)>=0:m.find(c,this,null,[i]).length),e[c]&&e.push(d);e.length&&g.push({elem:i,handlers:e})}return h<b.length&&g.push({elem:this,handlers:b.slice(h)}),g},fix:function(a){if(a[m.expando])return a;var b,c,d,e=a.type,f=a,g=this.fixHooks[e];g||(this.fixHooks[e]=g=Z.test(e)?this.mouseHooks:Y.test(e)?this.keyHooks:{}),d=g.props?this.props.concat(g.props):this.props,a=new m.Event(f),b=d.length;while(b--)c=d[b],a[c]=f[c];return a.target||(a.target=f.srcElement||y),3===a.target.nodeType&&(a.target=a.target.parentNode),a.metaKey=!!a.metaKey,g.filter?g.filter(a,f):a},props:"altKey bubbles cancelable ctrlKey currentTarget eventPhase metaKey relatedTarget shiftKey target timeStamp view which".split(" "),fixHooks:{},keyHooks:{props:"char charCode key keyCode".split(" "),filter:function(a,b){return null==a.which&&(a.which=null!=b.charCode?b.charCode:b.keyCode),a}},mouseHooks:{props:"button buttons clientX clientY fromElement offsetX offsetY pageX pageY screenX screenY toElement".split(" "),filter:function(a,b){var c,d,e,f=b.button,g=b.fromElement;return null==a.pageX&&null!=b.clientX&&(d=a.target.ownerDocument||y,e=d.documentElement,c=d.body,a.pageX=b.clientX+(e&&e.scrollLeft||c&&c.scrollLeft||0)-(e&&e.clientLeft||c&&c.clientLeft||0),a.pageY=b.clientY+(e&&e.scrollTop||c&&c.scrollTop||0)-(e&&e.clientTop||c&&c.clientTop||0)),!a.relatedTarget&&g&&(a.relatedTarget=g===a.target?b.toElement:g),a.which||void 0===f||(a.which=1&f?1:2&f?3:4&f?2:0),a}},special:{load:{noBubble:!0},focus:{trigger:function(){if(this!==cb()&&this.focus)try{return this.focus(),!1}catch(a){}},delegateType:"focusin"},blur:{trigger:function(){return this===cb()&&this.blur?(this.blur(),!1):void 0},delegateType:"focusout"},click:{trigger:function(){return m.nodeName(this,"input")&&"checkbox"===this.type&&this.click?(this.click(),!1):void 0},_default:function(a){return m.nodeName(a.target,"a")}},beforeunload:{postDispatch:function(a){void 0!==a.result&&a.originalEvent&&(a.originalEvent.returnValue=a.result)}}},simulate:function(a,b,c,d){var e=m.extend(new m.Event,c,{type:a,isSimulated:!0,originalEvent:{}});d?m.event.trigger(e,null,b):m.event.dispatch.call(b,e),e.isDefaultPrevented()&&c.preventDefault()}},m.removeEvent=y.removeEventListener?function(a,b,c){a.removeEventListener&&a.removeEventListener(b,c,!1)}:function(a,b,c){var d="on"+b;a.detachEvent&&(typeof a[d]===K&&(a[d]=null),a.detachEvent(d,c))},m.Event=function(a,b){return this instanceof m.Event?(a&&a.type?(this.originalEvent=a,this.type=a.type,this.isDefaultPrevented=a.defaultPrevented||void 0===a.defaultPrevented&&a.returnValue===!1?ab:bb):this.type=a,b&&m.extend(this,b),this.timeStamp=a&&a.timeStamp||m.now(),void(this[m.expando]=!0)):new m.Event(a,b)},m.Event.prototype={isDefaultPrevented:bb,isPropagationStopped:bb,isImmediatePropagationStopped:bb,preventDefault:function(){var a=this.originalEvent;this.isDefaultPrevented=ab,a&&(a.preventDefault?a.preventDefault():a.returnValue=!1)},stopPropagation:function(){var a=this.originalEvent;this.isPropagationStopped=ab,a&&(a.stopPropagation&&a.stopPropagation(),a.cancelBubble=!0)},stopImmediatePropagation:function(){var a=this.originalEvent;this.isImmediatePropagationStopped=ab,a&&a.stopImmediatePropagation&&a.stopImmediatePropagation(),this.stopPropagation()}},m.each({mouseenter:"mouseover",mouseleave:"mouseout",pointerenter:"pointerover",pointerleave:"pointerout"},function(a,b){m.event.special[a]={delegateType:b,bindType:b,handle:function(a){var c,d=this,e=a.relatedTarget,f=a.handleObj;return(!e||e!==d&&!m.contains(d,e))&&(a.type=f.origType,c=f.handler.apply(this,arguments),a.type=b),c}}}),k.submitBubbles||(m.event.special.submit={setup:function(){return m.nodeName(this,"form")?!1:void m.event.add(this,"click._submit keypress._submit",function(a){var b=a.target,c=m.nodeName(b,"input")||m.nodeName(b,"button")?b.form:void 0;c&&!m._data(c,"submitBubbles")&&(m.event.add(c,"submit._submit",function(a){a._submit_bubble=!0}),m._data(c,"submitBubbles",!0))})},postDispatch:function(a){a._submit_bubble&&(delete a._submit_bubble,this.parentNode&&!a.isTrigger&&m.event.simulate("submit",this.parentNode,a,!0))},teardown:function(){return m.nodeName(this,"form")?!1:void m.event.remove(this,"._submit")}}),k.changeBubbles||(m.event.special.change={setup:function(){return X.test(this.nodeName)?(("checkbox"===this.type||"radio"===this.type)&&(m.event.add(this,"propertychange._change",function(a){"checked"===a.originalEvent.propertyName&&(this._just_changed=!0)}),m.event.add(this,"click._change",function(a){this._just_changed&&!a.isTrigger&&(this._just_changed=!1),m.event.simulate("change",this,a,!0)})),!1):void m.event.add(this,"beforeactivate._change",function(a){var b=a.target;X.test(b.nodeName)&&!m._data(b,"changeBubbles")&&(m.event.add(b,"change._change",function(a){!this.parentNode||a.isSimulated||a.isTrigger||m.event.simulate("change",this.parentNode,a,!0)}),m._data(b,"changeBubbles",!0))})},handle:function(a){var b=a.target;return this!==b||a.isSimulated||a.isTrigger||"radio"!==b.type&&"checkbox"!==b.type?a.handleObj.handler.apply(this,arguments):void 0},teardown:function(){return m.event.remove(this,"._change"),!X.test(this.nodeName)}}),k.focusinBubbles||m.each({focus:"focusin",blur:"focusout"},function(a,b){var c=function(a){m.event.simulate(b,a.target,m.event.fix(a),!0)};m.event.special[b]={setup:function(){var d=this.ownerDocument||this,e=m._data(d,b);e||d.addEventListener(a,c,!0),m._data(d,b,(e||0)+1)},teardown:function(){var d=this.ownerDocument||this,e=m._data(d,b)-1;e?m._data(d,b,e):(d.removeEventListener(a,c,!0),m._removeData(d,b))}}}),m.fn.extend({on:function(a,b,c,d,e){var f,g;if("object"==typeof a){"string"!=typeof b&&(c=c||b,b=void 0);for(f in a)this.on(f,b,c,a[f],e);return this}if(null==c&&null==d?(d=b,c=b=void 0):null==d&&("string"==typeof b?(d=c,c=void 0):(d=c,c=b,b=void 0)),d===!1)d=bb;else if(!d)return this;return 1===e&&(g=d,d=function(a){return m().off(a),g.apply(this,arguments)},d.guid=g.guid||(g.guid=m.guid++)),this.each(function(){m.event.add(this,a,d,c,b)})},one:function(a,b,c,d){return this.on(a,b,c,d,1)},off:function(a,b,c){var d,e;if(a&&a.preventDefault&&a.handleObj)return d=a.handleObj,m(a.delegateTarget).off(d.namespace?d.origType+"."+d.namespace:d.origType,d.selector,d.handler),this;if("object"==typeof a){for(e in a)this.off(e,b,a[e]);return this}return(b===!1||"function"==typeof b)&&(c=b,b=void 0),c===!1&&(c=bb),this.each(function(){m.event.remove(this,a,c,b)})},trigger:function(a,b){return this.each(function(){m.event.trigger(a,b,this)})},triggerHandler:function(a,b){var c=this[0];return c?m.event.trigger(a,b,c,!0):void 0}});function db(a){var b=eb.split("|"),c=a.createDocumentFragment();if(c.createElement)while(b.length)c.createElement(b.pop());return c}var eb="abbr|article|aside|audio|bdi|canvas|data|datalist|details|figcaption|figure|footer|header|hgroup|mark|meter|nav|output|progress|section|summary|time|video",fb=/ jQuery\d+="(?:null|\d+)"/g,gb=new RegExp("<(?:"+eb+")[\\s/>]","i"),hb=/^\s+/,ib=/<(?!area|br|col|embed|hr|img|input|link|meta|param)(([\w:]+)[^>]*)\/>/gi,jb=/<([\w:]+)/,kb=/<tbody/i,lb=/<|&#?\w+;/,mb=/<(?:script|style|link)/i,nb=/checked\s*(?:[^=]|=\s*.checked.)/i,ob=/^$|\/(?:java|ecma)script/i,pb=/^true\/(.*)/,qb=/^\s*<!(?:\[CDATA\[|--)|(?:\]\]|--)>\s*$/g,rb={option:[1,"<select multiple='multiple'>","</select>"],legend:[1,"<fieldset>","</fieldset>"],area:[1,"<map>","</map>"],param:[1,"<object>","</object>"],thead:[1,"<table>","</table>"],tr:[2,"<table><tbody>","</tbody></table>"],col:[2,"<table><tbody></tbody><colgroup>","</colgroup></table>"],td:[3,"<table><tbody><tr>","</tr></tbody></table>"],_default:k.htmlSerialize?[0,"",""]:[1,"X<div>","</div>"]},sb=db(y),tb=sb.appendChild(y.createElement("div"));rb.optgroup=rb.option,rb.tbody=rb.tfoot=rb.colgroup=rb.caption=rb.thead,rb.th=rb.td;function ub(a,b){var c,d,e=0,f=typeof a.getElementsByTagName!==K?a.getElementsByTagName(b||"*"):typeof a.querySelectorAll!==K?a.querySelectorAll(b||"*"):void 0;if(!f)for(f=[],c=a.childNodes||a;null!=(d=c[e]);e++)!b||m.nodeName(d,b)?f.push(d):m.merge(f,ub(d,b));return void 0===b||b&&m.nodeName(a,b)?m.merge([a],f):f}function vb(a){W.test(a.type)&&(a.defaultChecked=a.checked)}function wb(a,b){return m.nodeName(a,"table")&&m.nodeName(11!==b.nodeType?b:b.firstChild,"tr")?a.getElementsByTagName("tbody")[0]||a.appendChild(a.ownerDocument.createElement("tbody")):a}function xb(a){return a.type=(null!==m.find.attr(a,"type"))+"/"+a.type,a}function yb(a){var b=pb.exec(a.type);return b?a.type=b[1]:a.removeAttribute("type"),a}function zb(a,b){for(var c,d=0;null!=(c=a[d]);d++)m._data(c,"globalEval",!b||m._data(b[d],"globalEval"))}function Ab(a,b){if(1===b.nodeType&&m.hasData(a)){var c,d,e,f=m._data(a),g=m._data(b,f),h=f.events;if(h){delete g.handle,g.events={};for(c in h)for(d=0,e=h[c].length;e>d;d++)m.event.add(b,c,h[c][d])}g.data&&(g.data=m.extend({},g.data))}}function Bb(a,b){var c,d,e;if(1===b.nodeType){if(c=b.nodeName.toLowerCase(),!k.noCloneEvent&&b[m.expando]){e=m._data(b);for(d in e.events)m.removeEvent(b,d,e.handle);b.removeAttribute(m.expando)}"script"===c&&b.text!==a.text?(xb(b).text=a.text,yb(b)):"object"===c?(b.parentNode&&(b.outerHTML=a.outerHTML),k.html5Clone&&a.innerHTML&&!m.trim(b.innerHTML)&&(b.innerHTML=a.innerHTML)):"input"===c&&W.test(a.type)?(b.defaultChecked=b.checked=a.checked,b.value!==a.value&&(b.value=a.value)):"option"===c?b.defaultSelected=b.selected=a.defaultSelected:("input"===c||"textarea"===c)&&(b.defaultValue=a.defaultValue)}}m.extend({clone:function(a,b,c){var d,e,f,g,h,i=m.contains(a.ownerDocument,a);if(k.html5Clone||m.isXMLDoc(a)||!gb.test("<"+a.nodeName+">")?f=a.cloneNode(!0):(tb.innerHTML=a.outerHTML,tb.removeChild(f=tb.firstChild)),!(k.noCloneEvent&&k.noCloneChecked||1!==a.nodeType&&11!==a.nodeType||m.isXMLDoc(a)))for(d=ub(f),h=ub(a),g=0;null!=(e=h[g]);++g)d[g]&&Bb(e,d[g]);if(b)if(c)for(h=h||ub(a),d=d||ub(f),g=0;null!=(e=h[g]);g++)Ab(e,d[g]);else Ab(a,f);return d=ub(f,"script"),d.length>0&&zb(d,!i&&ub(a,"script")),d=h=e=null,f},buildFragment:function(a,b,c,d){for(var e,f,g,h,i,j,l,n=a.length,o=db(b),p=[],q=0;n>q;q++)if(f=a[q],f||0===f)if("object"===m.type(f))m.merge(p,f.nodeType?[f]:f);else if(lb.test(f)){h=h||o.appendChild(b.createElement("div")),i=(jb.exec(f)||["",""])[1].toLowerCase(),l=rb[i]||rb._default,h.innerHTML=l[1]+f.replace(ib,"<$1></$2>")+l[2],e=l[0];while(e--)h=h.lastChild;if(!k.leadingWhitespace&&hb.test(f)&&p.push(b.createTextNode(hb.exec(f)[0])),!k.tbody){f="table"!==i||kb.test(f)?"<table>"!==l[1]||kb.test(f)?0:h:h.firstChild,e=f&&f.childNodes.length;while(e--)m.nodeName(j=f.childNodes[e],"tbody")&&!j.childNodes.length&&f.removeChild(j)}m.merge(p,h.childNodes),h.textContent="";while(h.firstChild)h.removeChild(h.firstChild);h=o.lastChild}else p.push(b.createTextNode(f));h&&o.removeChild(h),k.appendChecked||m.grep(ub(p,"input"),vb),q=0;while(f=p[q++])if((!d||-1===m.inArray(f,d))&&(g=m.contains(f.ownerDocument,f),h=ub(o.appendChild(f),"script"),g&&zb(h),c)){e=0;while(f=h[e++])ob.test(f.type||"")&&c.push(f)}return h=null,o},cleanData:function(a,b){for(var d,e,f,g,h=0,i=m.expando,j=m.cache,l=k.deleteExpando,n=m.event.special;null!=(d=a[h]);h++)if((b||m.acceptData(d))&&(f=d[i],g=f&&j[f])){if(g.events)for(e in g.events)n[e]?m.event.remove(d,e):m.removeEvent(d,e,g.handle);j[f]&&(delete j[f],l?delete d[i]:typeof d.removeAttribute!==K?d.removeAttribute(i):d[i]=null,c.push(f))}}}),m.fn.extend({text:function(a){return V(this,function(a){return void 0===a?m.text(this):this.empty().append((this[0]&&this[0].ownerDocument||y).createTextNode(a))},null,a,arguments.length)},append:function(){return this.domManip(arguments,function(a){if(1===this.nodeType||11===this.nodeType||9===this.nodeType){var b=wb(this,a);b.appendChild(a)}})},prepend:function(){return this.domManip(arguments,function(a){if(1===this.nodeType||11===this.nodeType||9===this.nodeType){var b=wb(this,a);b.insertBefore(a,b.firstChild)}})},before:function(){return this.domManip(arguments,function(a){this.parentNode&&this.parentNode.insertBefore(a,this)})},after:function(){return this.domManip(arguments,function(a){this.parentNode&&this.parentNode.insertBefore(a,this.nextSibling)})},remove:function(a,b){for(var c,d=a?m.filter(a,this):this,e=0;null!=(c=d[e]);e++)b||1!==c.nodeType||m.cleanData(ub(c)),c.parentNode&&(b&&m.contains(c.ownerDocument,c)&&zb(ub(c,"script")),c.parentNode.removeChild(c));return this},empty:function(){for(var a,b=0;null!=(a=this[b]);b++){1===a.nodeType&&m.cleanData(ub(a,!1));while(a.firstChild)a.removeChild(a.firstChild);a.options&&m.nodeName(a,"select")&&(a.options.length=0)}return this},clone:function(a,b){return a=null==a?!1:a,b=null==b?a:b,this.map(function(){return m.clone(this,a,b)})},html:function(a){return V(this,function(a){var b=this[0]||{},c=0,d=this.length;if(void 0===a)return 1===b.nodeType?b.innerHTML.replace(fb,""):void 0;if(!("string"!=typeof a||mb.test(a)||!k.htmlSerialize&&gb.test(a)||!k.leadingWhitespace&&hb.test(a)||rb[(jb.exec(a)||["",""])[1].toLowerCase()])){a=a.replace(ib,"<$1></$2>");try{for(;d>c;c++)b=this[c]||{},1===b.nodeType&&(m.cleanData(ub(b,!1)),b.innerHTML=a);b=0}catch(e){}}b&&this.empty().append(a)},null,a,arguments.length)},replaceWith:function(){var a=arguments[0];return this.domManip(arguments,function(b){a=this.parentNode,m.cleanData(ub(this)),a&&a.replaceChild(b,this)}),a&&(a.length||a.nodeType)?this:this.remove()},detach:function(a){return this.remove(a,!0)},domManip:function(a,b){a=e.apply([],a);var c,d,f,g,h,i,j=0,l=this.length,n=this,o=l-1,p=a[0],q=m.isFunction(p);if(q||l>1&&"string"==typeof p&&!k.checkClone&&nb.test(p))return this.each(function(c){var d=n.eq(c);q&&(a[0]=p.call(this,c,d.html())),d.domManip(a,b)});if(l&&(i=m.buildFragment(a,this[0].ownerDocument,!1,this),c=i.firstChild,1===i.childNodes.length&&(i=c),c)){for(g=m.map(ub(i,"script"),xb),f=g.length;l>j;j++)d=i,j!==o&&(d=m.clone(d,!0,!0),f&&m.merge(g,ub(d,"script"))),b.call(this[j],d,j);if(f)for(h=g[g.length-1].ownerDocument,m.map(g,yb),j=0;f>j;j++)d=g[j],ob.test(d.type||"")&&!m._data(d,"globalEval")&&m.contains(h,d)&&(d.src?m._evalUrl&&m._evalUrl(d.src):m.globalEval((d.text||d.textContent||d.innerHTML||"").replace(qb,"")));i=c=null}return this}}),m.each({appendTo:"append",prependTo:"prepend",insertBefore:"before",insertAfter:"after",replaceAll:"replaceWith"},function(a,b){m.fn[a]=function(a){for(var c,d=0,e=[],g=m(a),h=g.length-1;h>=d;d++)c=d===h?this:this.clone(!0),m(g[d])[b](c),f.apply(e,c.get());return this.pushStack(e)}});var Cb,Db={};function Eb(b,c){var d,e=m(c.createElement(b)).appendTo(c.body),f=a.getDefaultComputedStyle&&(d=a.getDefaultComputedStyle(e[0]))?d.display:m.css(e[0],"display");return e.detach(),f}function Fb(a){var b=y,c=Db[a];return c||(c=Eb(a,b),"none"!==c&&c||(Cb=(Cb||m("<iframe frameborder='0' width='0' height='0'/>")).appendTo(b.documentElement),b=(Cb[0].contentWindow||Cb[0].contentDocument).document,b.write(),b.close(),c=Eb(a,b),Cb.detach()),Db[a]=c),c}!function(){var a;k.shrinkWrapBlocks=function(){if(null!=a)return a;a=!1;var b,c,d;return c=y.getElementsByTagName("body")[0],c&&c.style?(b=y.createElement("div"),d=y.createElement("div"),d.style.cssText="position:absolute;border:0;width:0;height:0;top:0;left:-9999px",c.appendChild(d).appendChild(b),typeof b.style.zoom!==K&&(b.style.cssText="-webkit-box-sizing:content-box;-moz-box-sizing:content-box;box-sizing:content-box;display:block;margin:0;border:0;padding:1px;width:1px;zoom:1",b.appendChild(y.createElement("div")).style.width="5px",a=3!==b.offsetWidth),c.removeChild(d),a):void 0}}();var Gb=/^margin/,Hb=new RegExp("^("+S+")(?!px)[a-z%]+$","i"),Ib,Jb,Kb=/^(top|right|bottom|left)$/;a.getComputedStyle?(Ib=function(b){return b.ownerDocument.defaultView.opener?b.ownerDocument.defaultView.getComputedStyle(b,null):a.getComputedStyle(b,null)},Jb=function(a,b,c){var d,e,f,g,h=a.style;return c=c||Ib(a),g=c?c.getPropertyValue(b)||c[b]:void 0,c&&(""!==g||m.contains(a.ownerDocument,a)||(g=m.style(a,b)),Hb.test(g)&&Gb.test(b)&&(d=h.width,e=h.minWidth,f=h.maxWidth,h.minWidth=h.maxWidth=h.width=g,g=c.width,h.width=d,h.minWidth=e,h.maxWidth=f)),void 0===g?g:g+""}):y.documentElement.currentStyle&&(Ib=function(a){return a.currentStyle},Jb=function(a,b,c){var d,e,f,g,h=a.style;return c=c||Ib(a),g=c?c[b]:void 0,null==g&&h&&h[b]&&(g=h[b]),Hb.test(g)&&!Kb.test(b)&&(d=h.left,e=a.runtimeStyle,f=e&&e.left,f&&(e.left=a.currentStyle.left),h.left="fontSize"===b?"1em":g,g=h.pixelLeft+"px",h.left=d,f&&(e.left=f)),void 0===g?g:g+""||"auto"});function Lb(a,b){return{get:function(){var c=a();if(null!=c)return c?void delete this.get:(this.get=b).apply(this,arguments)}}}!function(){var b,c,d,e,f,g,h;if(b=y.createElement("div"),b.innerHTML="  <link/><table></table><a href='/a'>a</a><input type='checkbox'/>",d=b.getElementsByTagName("a")[0],c=d&&d.style){c.cssText="float:left;opacity:.5",k.opacity="0.5"===c.opacity,k.cssFloat=!!c.cssFloat,b.style.backgroundClip="content-box",b.cloneNode(!0).style.backgroundClip="",k.clearCloneStyle="content-box"===b.style.backgroundClip,k.boxSizing=""===c.boxSizing||""===c.MozBoxSizing||""===c.WebkitBoxSizing,m.extend(k,{reliableHiddenOffsets:function(){return null==g&&i(),g},boxSizingReliable:function(){return null==f&&i(),f},pixelPosition:function(){return null==e&&i(),e},reliableMarginRight:function(){return null==h&&i(),h}});function i(){var b,c,d,i;c=y.getElementsByTagName("body")[0],c&&c.style&&(b=y.createElement("div"),d=y.createElement("div"),d.style.cssText="position:absolute;border:0;width:0;height:0;top:0;left:-9999px",c.appendChild(d).appendChild(b),b.style.cssText="-webkit-box-sizing:border-box;-moz-box-sizing:border-box;box-sizing:border-box;display:block;margin-top:1%;top:1%;border:1px;padding:1px;width:4px;position:absolute",e=f=!1,h=!0,a.getComputedStyle&&(e="1%"!==(a.getComputedStyle(b,null)||{}).top,f="4px"===(a.getComputedStyle(b,null)||{width:"4px"}).width,i=b.appendChild(y.createElement("div")),i.style.cssText=b.style.cssText="-webkit-box-sizing:content-box;-moz-box-sizing:content-box;box-sizing:content-box;display:block;margin:0;border:0;padding:0",i.style.marginRight=i.style.width="0",b.style.width="1px",h=!parseFloat((a.getComputedStyle(i,null)||{}).marginRight),b.removeChild(i)),b.innerHTML="<table><tr><td></td><td>t</td></tr></table>",i=b.getElementsByTagName("td"),i[0].style.cssText="margin:0;border:0;padding:0;display:none",g=0===i[0].offsetHeight,g&&(i[0].style.display="",i[1].style.display="none",g=0===i[0].offsetHeight),c.removeChild(d))}}}(),m.swap=function(a,b,c,d){var e,f,g={};for(f in b)g[f]=a.style[f],a.style[f]=b[f];e=c.apply(a,d||[]);for(f in b)a.style[f]=g[f];return e};var Mb=/alpha\([^)]*\)/i,Nb=/opacity\s*=\s*([^)]*)/,Ob=/^(none|table(?!-c[ea]).+)/,Pb=new RegExp("^("+S+")(.*)$","i"),Qb=new RegExp("^([+-])=("+S+")","i"),Rb={position:"absolute",visibility:"hidden",display:"block"},Sb={letterSpacing:"0",fontWeight:"400"},Tb=["Webkit","O","Moz","ms"];function Ub(a,b){if(b in a)return b;var c=b.charAt(0).toUpperCase()+b.slice(1),d=b,e=Tb.length;while(e--)if(b=Tb[e]+c,b in a)return b;return d}function Vb(a,b){for(var c,d,e,f=[],g=0,h=a.length;h>g;g++)d=a[g],d.style&&(f[g]=m._data(d,"olddisplay"),c=d.style.display,b?(f[g]||"none"!==c||(d.style.display=""),""===d.style.display&&U(d)&&(f[g]=m._data(d,"olddisplay",Fb(d.nodeName)))):(e=U(d),(c&&"none"!==c||!e)&&m._data(d,"olddisplay",e?c:m.css(d,"display"))));for(g=0;h>g;g++)d=a[g],d.style&&(b&&"none"!==d.style.display&&""!==d.style.display||(d.style.display=b?f[g]||"":"none"));return a}function Wb(a,b,c){var d=Pb.exec(b);return d?Math.max(0,d[1]-(c||0))+(d[2]||"px"):b}function Xb(a,b,c,d,e){for(var f=c===(d?"border":"content")?4:"width"===b?1:0,g=0;4>f;f+=2)"margin"===c&&(g+=m.css(a,c+T[f],!0,e)),d?("content"===c&&(g-=m.css(a,"padding"+T[f],!0,e)),"margin"!==c&&(g-=m.css(a,"border"+T[f]+"Width",!0,e))):(g+=m.css(a,"padding"+T[f],!0,e),"padding"!==c&&(g+=m.css(a,"border"+T[f]+"Width",!0,e)));return g}function Yb(a,b,c){var d=!0,e="width"===b?a.offsetWidth:a.offsetHeight,f=Ib(a),g=k.boxSizing&&"border-box"===m.css(a,"boxSizing",!1,f);if(0>=e||null==e){if(e=Jb(a,b,f),(0>e||null==e)&&(e=a.style[b]),Hb.test(e))return e;d=g&&(k.boxSizingReliable()||e===a.style[b]),e=parseFloat(e)||0}return e+Xb(a,b,c||(g?"border":"content"),d,f)+"px"}m.extend({cssHooks:{opacity:{get:function(a,b){if(b){var c=Jb(a,"opacity");return""===c?"1":c}}}},cssNumber:{columnCount:!0,fillOpacity:!0,flexGrow:!0,flexShrink:!0,fontWeight:!0,lineHeight:!0,opacity:!0,order:!0,orphans:!0,widows:!0,zIndex:!0,zoom:!0},cssProps:{"float":k.cssFloat?"cssFloat":"styleFloat"},style:function(a,b,c,d){if(a&&3!==a.nodeType&&8!==a.nodeType&&a.style){var e,f,g,h=m.camelCase(b),i=a.style;if(b=m.cssProps[h]||(m.cssProps[h]=Ub(i,h)),g=m.cssHooks[b]||m.cssHooks[h],void 0===c)return g&&"get"in g&&void 0!==(e=g.get(a,!1,d))?e:i[b];if(f=typeof c,"string"===f&&(e=Qb.exec(c))&&(c=(e[1]+1)*e[2]+parseFloat(m.css(a,b)),f="number"),null!=c&&c===c&&("number"!==f||m.cssNumber[h]||(c+="px"),k.clearCloneStyle||""!==c||0!==b.indexOf("background")||(i[b]="inherit"),!(g&&"set"in g&&void 0===(c=g.set(a,c,d)))))try{i[b]=c}catch(j){}}},css:function(a,b,c,d){var e,f,g,h=m.camelCase(b);return b=m.cssProps[h]||(m.cssProps[h]=Ub(a.style,h)),g=m.cssHooks[b]||m.cssHooks[h],g&&"get"in g&&(f=g.get(a,!0,c)),void 0===f&&(f=Jb(a,b,d)),"normal"===f&&b in Sb&&(f=Sb[b]),""===c||c?(e=parseFloat(f),c===!0||m.isNumeric(e)?e||0:f):f}}),m.each(["height","width"],function(a,b){m.cssHooks[b]={get:function(a,c,d){return c?Ob.test(m.css(a,"display"))&&0===a.offsetWidth?m.swap(a,Rb,function(){return Yb(a,b,d)}):Yb(a,b,d):void 0},set:function(a,c,d){var e=d&&Ib(a);return Wb(a,c,d?Xb(a,b,d,k.boxSizing&&"border-box"===m.css(a,"boxSizing",!1,e),e):0)}}}),k.opacity||(m.cssHooks.opacity={get:function(a,b){return Nb.test((b&&a.currentStyle?a.currentStyle.filter:a.style.filter)||"")?.01*parseFloat(RegExp.$1)+"":b?"1":""},set:function(a,b){var c=a.style,d=a.currentStyle,e=m.isNumeric(b)?"alpha(opacity="+100*b+")":"",f=d&&d.filter||c.filter||"";c.zoom=1,(b>=1||""===b)&&""===m.trim(f.replace(Mb,""))&&c.removeAttribute&&(c.removeAttribute("filter"),""===b||d&&!d.filter)||(c.filter=Mb.test(f)?f.replace(Mb,e):f+" "+e)}}),m.cssHooks.marginRight=Lb(k.reliableMarginRight,function(a,b){return b?m.swap(a,{display:"inline-block"},Jb,[a,"marginRight"]):void 0}),m.each({margin:"",padding:"",border:"Width"},function(a,b){m.cssHooks[a+b]={expand:function(c){for(var d=0,e={},f="string"==typeof c?c.split(" "):[c];4>d;d++)e[a+T[d]+b]=f[d]||f[d-2]||f[0];return e}},Gb.test(a)||(m.cssHooks[a+b].set=Wb)}),m.fn.extend({css:function(a,b){return V(this,function(a,b,c){var d,e,f={},g=0;if(m.isArray(b)){for(d=Ib(a),e=b.length;e>g;g++)f[b[g]]=m.css(a,b[g],!1,d);return f}return void 0!==c?m.style(a,b,c):m.css(a,b)},a,b,arguments.length>1)},show:function(){return Vb(this,!0)},hide:function(){return Vb(this)},toggle:function(a){return"boolean"==typeof a?a?this.show():this.hide():this.each(function(){U(this)?m(this).show():m(this).hide()})}});function Zb(a,b,c,d,e){return new Zb.prototype.init(a,b,c,d,e)
}m.Tween=Zb,Zb.prototype={constructor:Zb,init:function(a,b,c,d,e,f){this.elem=a,this.prop=c,this.easing=e||"swing",this.options=b,this.start=this.now=this.cur(),this.end=d,this.unit=f||(m.cssNumber[c]?"":"px")},cur:function(){var a=Zb.propHooks[this.prop];return a&&a.get?a.get(this):Zb.propHooks._default.get(this)},run:function(a){var b,c=Zb.propHooks[this.prop];return this.pos=b=this.options.duration?m.easing[this.easing](a,this.options.duration*a,0,1,this.options.duration):a,this.now=(this.end-this.start)*b+this.start,this.options.step&&this.options.step.call(this.elem,this.now,this),c&&c.set?c.set(this):Zb.propHooks._default.set(this),this}},Zb.prototype.init.prototype=Zb.prototype,Zb.propHooks={_default:{get:function(a){var b;return null==a.elem[a.prop]||a.elem.style&&null!=a.elem.style[a.prop]?(b=m.css(a.elem,a.prop,""),b&&"auto"!==b?b:0):a.elem[a.prop]},set:function(a){m.fx.step[a.prop]?m.fx.step[a.prop](a):a.elem.style&&(null!=a.elem.style[m.cssProps[a.prop]]||m.cssHooks[a.prop])?m.style(a.elem,a.prop,a.now+a.unit):a.elem[a.prop]=a.now}}},Zb.propHooks.scrollTop=Zb.propHooks.scrollLeft={set:function(a){a.elem.nodeType&&a.elem.parentNode&&(a.elem[a.prop]=a.now)}},m.easing={linear:function(a){return a},swing:function(a){return.5-Math.cos(a*Math.PI)/2}},m.fx=Zb.prototype.init,m.fx.step={};var $b,_b,ac=/^(?:toggle|show|hide)$/,bc=new RegExp("^(?:([+-])=|)("+S+")([a-z%]*)$","i"),cc=/queueHooks$/,dc=[ic],ec={"*":[function(a,b){var c=this.createTween(a,b),d=c.cur(),e=bc.exec(b),f=e&&e[3]||(m.cssNumber[a]?"":"px"),g=(m.cssNumber[a]||"px"!==f&&+d)&&bc.exec(m.css(c.elem,a)),h=1,i=20;if(g&&g[3]!==f){f=f||g[3],e=e||[],g=+d||1;do h=h||".5",g/=h,m.style(c.elem,a,g+f);while(h!==(h=c.cur()/d)&&1!==h&&--i)}return e&&(g=c.start=+g||+d||0,c.unit=f,c.end=e[1]?g+(e[1]+1)*e[2]:+e[2]),c}]};function fc(){return setTimeout(function(){$b=void 0}),$b=m.now()}function gc(a,b){var c,d={height:a},e=0;for(b=b?1:0;4>e;e+=2-b)c=T[e],d["margin"+c]=d["padding"+c]=a;return b&&(d.opacity=d.width=a),d}function hc(a,b,c){for(var d,e=(ec[b]||[]).concat(ec["*"]),f=0,g=e.length;g>f;f++)if(d=e[f].call(c,b,a))return d}function ic(a,b,c){var d,e,f,g,h,i,j,l,n=this,o={},p=a.style,q=a.nodeType&&U(a),r=m._data(a,"fxshow");c.queue||(h=m._queueHooks(a,"fx"),null==h.unqueued&&(h.unqueued=0,i=h.empty.fire,h.empty.fire=function(){h.unqueued||i()}),h.unqueued++,n.always(function(){n.always(function(){h.unqueued--,m.queue(a,"fx").length||h.empty.fire()})})),1===a.nodeType&&("height"in b||"width"in b)&&(c.overflow=[p.overflow,p.overflowX,p.overflowY],j=m.css(a,"display"),l="none"===j?m._data(a,"olddisplay")||Fb(a.nodeName):j,"inline"===l&&"none"===m.css(a,"float")&&(k.inlineBlockNeedsLayout&&"inline"!==Fb(a.nodeName)?p.zoom=1:p.display="inline-block")),c.overflow&&(p.overflow="hidden",k.shrinkWrapBlocks()||n.always(function(){p.overflow=c.overflow[0],p.overflowX=c.overflow[1],p.overflowY=c.overflow[2]}));for(d in b)if(e=b[d],ac.exec(e)){if(delete b[d],f=f||"toggle"===e,e===(q?"hide":"show")){if("show"!==e||!r||void 0===r[d])continue;q=!0}o[d]=r&&r[d]||m.style(a,d)}else j=void 0;if(m.isEmptyObject(o))"inline"===("none"===j?Fb(a.nodeName):j)&&(p.display=j);else{r?"hidden"in r&&(q=r.hidden):r=m._data(a,"fxshow",{}),f&&(r.hidden=!q),q?m(a).show():n.done(function(){m(a).hide()}),n.done(function(){var b;m._removeData(a,"fxshow");for(b in o)m.style(a,b,o[b])});for(d in o)g=hc(q?r[d]:0,d,n),d in r||(r[d]=g.start,q&&(g.end=g.start,g.start="width"===d||"height"===d?1:0))}}function jc(a,b){var c,d,e,f,g;for(c in a)if(d=m.camelCase(c),e=b[d],f=a[c],m.isArray(f)&&(e=f[1],f=a[c]=f[0]),c!==d&&(a[d]=f,delete a[c]),g=m.cssHooks[d],g&&"expand"in g){f=g.expand(f),delete a[d];for(c in f)c in a||(a[c]=f[c],b[c]=e)}else b[d]=e}function kc(a,b,c){var d,e,f=0,g=dc.length,h=m.Deferred().always(function(){delete i.elem}),i=function(){if(e)return!1;for(var b=$b||fc(),c=Math.max(0,j.startTime+j.duration-b),d=c/j.duration||0,f=1-d,g=0,i=j.tweens.length;i>g;g++)j.tweens[g].run(f);return h.notifyWith(a,[j,f,c]),1>f&&i?c:(h.resolveWith(a,[j]),!1)},j=h.promise({elem:a,props:m.extend({},b),opts:m.extend(!0,{specialEasing:{}},c),originalProperties:b,originalOptions:c,startTime:$b||fc(),duration:c.duration,tweens:[],createTween:function(b,c){var d=m.Tween(a,j.opts,b,c,j.opts.specialEasing[b]||j.opts.easing);return j.tweens.push(d),d},stop:function(b){var c=0,d=b?j.tweens.length:0;if(e)return this;for(e=!0;d>c;c++)j.tweens[c].run(1);return b?h.resolveWith(a,[j,b]):h.rejectWith(a,[j,b]),this}}),k=j.props;for(jc(k,j.opts.specialEasing);g>f;f++)if(d=dc[f].call(j,a,k,j.opts))return d;return m.map(k,hc,j),m.isFunction(j.opts.start)&&j.opts.start.call(a,j),m.fx.timer(m.extend(i,{elem:a,anim:j,queue:j.opts.queue})),j.progress(j.opts.progress).done(j.opts.done,j.opts.complete).fail(j.opts.fail).always(j.opts.always)}m.Animation=m.extend(kc,{tweener:function(a,b){m.isFunction(a)?(b=a,a=["*"]):a=a.split(" ");for(var c,d=0,e=a.length;e>d;d++)c=a[d],ec[c]=ec[c]||[],ec[c].unshift(b)},prefilter:function(a,b){b?dc.unshift(a):dc.push(a)}}),m.speed=function(a,b,c){var d=a&&"object"==typeof a?m.extend({},a):{complete:c||!c&&b||m.isFunction(a)&&a,duration:a,easing:c&&b||b&&!m.isFunction(b)&&b};return d.duration=m.fx.off?0:"number"==typeof d.duration?d.duration:d.duration in m.fx.speeds?m.fx.speeds[d.duration]:m.fx.speeds._default,(null==d.queue||d.queue===!0)&&(d.queue="fx"),d.old=d.complete,d.complete=function(){m.isFunction(d.old)&&d.old.call(this),d.queue&&m.dequeue(this,d.queue)},d},m.fn.extend({fadeTo:function(a,b,c,d){return this.filter(U).css("opacity",0).show().end().animate({opacity:b},a,c,d)},animate:function(a,b,c,d){var e=m.isEmptyObject(a),f=m.speed(b,c,d),g=function(){var b=kc(this,m.extend({},a),f);(e||m._data(this,"finish"))&&b.stop(!0)};return g.finish=g,e||f.queue===!1?this.each(g):this.queue(f.queue,g)},stop:function(a,b,c){var d=function(a){var b=a.stop;delete a.stop,b(c)};return"string"!=typeof a&&(c=b,b=a,a=void 0),b&&a!==!1&&this.queue(a||"fx",[]),this.each(function(){var b=!0,e=null!=a&&a+"queueHooks",f=m.timers,g=m._data(this);if(e)g[e]&&g[e].stop&&d(g[e]);else for(e in g)g[e]&&g[e].stop&&cc.test(e)&&d(g[e]);for(e=f.length;e--;)f[e].elem!==this||null!=a&&f[e].queue!==a||(f[e].anim.stop(c),b=!1,f.splice(e,1));(b||!c)&&m.dequeue(this,a)})},finish:function(a){return a!==!1&&(a=a||"fx"),this.each(function(){var b,c=m._data(this),d=c[a+"queue"],e=c[a+"queueHooks"],f=m.timers,g=d?d.length:0;for(c.finish=!0,m.queue(this,a,[]),e&&e.stop&&e.stop.call(this,!0),b=f.length;b--;)f[b].elem===this&&f[b].queue===a&&(f[b].anim.stop(!0),f.splice(b,1));for(b=0;g>b;b++)d[b]&&d[b].finish&&d[b].finish.call(this);delete c.finish})}}),m.each(["toggle","show","hide"],function(a,b){var c=m.fn[b];m.fn[b]=function(a,d,e){return null==a||"boolean"==typeof a?c.apply(this,arguments):this.animate(gc(b,!0),a,d,e)}}),m.each({slideDown:gc("show"),slideUp:gc("hide"),slideToggle:gc("toggle"),fadeIn:{opacity:"show"},fadeOut:{opacity:"hide"},fadeToggle:{opacity:"toggle"}},function(a,b){m.fn[a]=function(a,c,d){return this.animate(b,a,c,d)}}),m.timers=[],m.fx.tick=function(){var a,b=m.timers,c=0;for($b=m.now();c<b.length;c++)a=b[c],a()||b[c]!==a||b.splice(c--,1);b.length||m.fx.stop(),$b=void 0},m.fx.timer=function(a){m.timers.push(a),a()?m.fx.start():m.timers.pop()},m.fx.interval=13,m.fx.start=function(){_b||(_b=setInterval(m.fx.tick,m.fx.interval))},m.fx.stop=function(){clearInterval(_b),_b=null},m.fx.speeds={slow:600,fast:200,_default:400},m.fn.delay=function(a,b){return a=m.fx?m.fx.speeds[a]||a:a,b=b||"fx",this.queue(b,function(b,c){var d=setTimeout(b,a);c.stop=function(){clearTimeout(d)}})},function(){var a,b,c,d,e;b=y.createElement("div"),b.setAttribute("className","t"),b.innerHTML="  <link/><table></table><a href='/a'>a</a><input type='checkbox'/>",d=b.getElementsByTagName("a")[0],c=y.createElement("select"),e=c.appendChild(y.createElement("option")),a=b.getElementsByTagName("input")[0],d.style.cssText="top:1px",k.getSetAttribute="t"!==b.className,k.style=/top/.test(d.getAttribute("style")),k.hrefNormalized="/a"===d.getAttribute("href"),k.checkOn=!!a.value,k.optSelected=e.selected,k.enctype=!!y.createElement("form").enctype,c.disabled=!0,k.optDisabled=!e.disabled,a=y.createElement("input"),a.setAttribute("value",""),k.input=""===a.getAttribute("value"),a.value="t",a.setAttribute("type","radio"),k.radioValue="t"===a.value}();var lc=/\r/g;m.fn.extend({val:function(a){var b,c,d,e=this[0];{if(arguments.length)return d=m.isFunction(a),this.each(function(c){var e;1===this.nodeType&&(e=d?a.call(this,c,m(this).val()):a,null==e?e="":"number"==typeof e?e+="":m.isArray(e)&&(e=m.map(e,function(a){return null==a?"":a+""})),b=m.valHooks[this.type]||m.valHooks[this.nodeName.toLowerCase()],b&&"set"in b&&void 0!==b.set(this,e,"value")||(this.value=e))});if(e)return b=m.valHooks[e.type]||m.valHooks[e.nodeName.toLowerCase()],b&&"get"in b&&void 0!==(c=b.get(e,"value"))?c:(c=e.value,"string"==typeof c?c.replace(lc,""):null==c?"":c)}}}),m.extend({valHooks:{option:{get:function(a){var b=m.find.attr(a,"value");return null!=b?b:m.trim(m.text(a))}},select:{get:function(a){for(var b,c,d=a.options,e=a.selectedIndex,f="select-one"===a.type||0>e,g=f?null:[],h=f?e+1:d.length,i=0>e?h:f?e:0;h>i;i++)if(c=d[i],!(!c.selected&&i!==e||(k.optDisabled?c.disabled:null!==c.getAttribute("disabled"))||c.parentNode.disabled&&m.nodeName(c.parentNode,"optgroup"))){if(b=m(c).val(),f)return b;g.push(b)}return g},set:function(a,b){var c,d,e=a.options,f=m.makeArray(b),g=e.length;while(g--)if(d=e[g],m.inArray(m.valHooks.option.get(d),f)>=0)try{d.selected=c=!0}catch(h){d.scrollHeight}else d.selected=!1;return c||(a.selectedIndex=-1),e}}}}),m.each(["radio","checkbox"],function(){m.valHooks[this]={set:function(a,b){return m.isArray(b)?a.checked=m.inArray(m(a).val(),b)>=0:void 0}},k.checkOn||(m.valHooks[this].get=function(a){return null===a.getAttribute("value")?"on":a.value})});var mc,nc,oc=m.expr.attrHandle,pc=/^(?:checked|selected)$/i,qc=k.getSetAttribute,rc=k.input;m.fn.extend({attr:function(a,b){return V(this,m.attr,a,b,arguments.length>1)},removeAttr:function(a){return this.each(function(){m.removeAttr(this,a)})}}),m.extend({attr:function(a,b,c){var d,e,f=a.nodeType;if(a&&3!==f&&8!==f&&2!==f)return typeof a.getAttribute===K?m.prop(a,b,c):(1===f&&m.isXMLDoc(a)||(b=b.toLowerCase(),d=m.attrHooks[b]||(m.expr.match.bool.test(b)?nc:mc)),void 0===c?d&&"get"in d&&null!==(e=d.get(a,b))?e:(e=m.find.attr(a,b),null==e?void 0:e):null!==c?d&&"set"in d&&void 0!==(e=d.set(a,c,b))?e:(a.setAttribute(b,c+""),c):void m.removeAttr(a,b))},removeAttr:function(a,b){var c,d,e=0,f=b&&b.match(E);if(f&&1===a.nodeType)while(c=f[e++])d=m.propFix[c]||c,m.expr.match.bool.test(c)?rc&&qc||!pc.test(c)?a[d]=!1:a[m.camelCase("default-"+c)]=a[d]=!1:m.attr(a,c,""),a.removeAttribute(qc?c:d)},attrHooks:{type:{set:function(a,b){if(!k.radioValue&&"radio"===b&&m.nodeName(a,"input")){var c=a.value;return a.setAttribute("type",b),c&&(a.value=c),b}}}}}),nc={set:function(a,b,c){return b===!1?m.removeAttr(a,c):rc&&qc||!pc.test(c)?a.setAttribute(!qc&&m.propFix[c]||c,c):a[m.camelCase("default-"+c)]=a[c]=!0,c}},m.each(m.expr.match.bool.source.match(/\w+/g),function(a,b){var c=oc[b]||m.find.attr;oc[b]=rc&&qc||!pc.test(b)?function(a,b,d){var e,f;return d||(f=oc[b],oc[b]=e,e=null!=c(a,b,d)?b.toLowerCase():null,oc[b]=f),e}:function(a,b,c){return c?void 0:a[m.camelCase("default-"+b)]?b.toLowerCase():null}}),rc&&qc||(m.attrHooks.value={set:function(a,b,c){return m.nodeName(a,"input")?void(a.defaultValue=b):mc&&mc.set(a,b,c)}}),qc||(mc={set:function(a,b,c){var d=a.getAttributeNode(c);return d||a.setAttributeNode(d=a.ownerDocument.createAttribute(c)),d.value=b+="","value"===c||b===a.getAttribute(c)?b:void 0}},oc.id=oc.name=oc.coords=function(a,b,c){var d;return c?void 0:(d=a.getAttributeNode(b))&&""!==d.value?d.value:null},m.valHooks.button={get:function(a,b){var c=a.getAttributeNode(b);return c&&c.specified?c.value:void 0},set:mc.set},m.attrHooks.contenteditable={set:function(a,b,c){mc.set(a,""===b?!1:b,c)}},m.each(["width","height"],function(a,b){m.attrHooks[b]={set:function(a,c){return""===c?(a.setAttribute(b,"auto"),c):void 0}}})),k.style||(m.attrHooks.style={get:function(a){return a.style.cssText||void 0},set:function(a,b){return a.style.cssText=b+""}});var sc=/^(?:input|select|textarea|button|object)$/i,tc=/^(?:a|area)$/i;m.fn.extend({prop:function(a,b){return V(this,m.prop,a,b,arguments.length>1)},removeProp:function(a){return a=m.propFix[a]||a,this.each(function(){try{this[a]=void 0,delete this[a]}catch(b){}})}}),m.extend({propFix:{"for":"htmlFor","class":"className"},prop:function(a,b,c){var d,e,f,g=a.nodeType;if(a&&3!==g&&8!==g&&2!==g)return f=1!==g||!m.isXMLDoc(a),f&&(b=m.propFix[b]||b,e=m.propHooks[b]),void 0!==c?e&&"set"in e&&void 0!==(d=e.set(a,c,b))?d:a[b]=c:e&&"get"in e&&null!==(d=e.get(a,b))?d:a[b]},propHooks:{tabIndex:{get:function(a){var b=m.find.attr(a,"tabindex");return b?parseInt(b,10):sc.test(a.nodeName)||tc.test(a.nodeName)&&a.href?0:-1}}}}),k.hrefNormalized||m.each(["href","src"],function(a,b){m.propHooks[b]={get:function(a){return a.getAttribute(b,4)}}}),k.optSelected||(m.propHooks.selected={get:function(a){var b=a.parentNode;return b&&(b.selectedIndex,b.parentNode&&b.parentNode.selectedIndex),null}}),m.each(["tabIndex","readOnly","maxLength","cellSpacing","cellPadding","rowSpan","colSpan","useMap","frameBorder","contentEditable"],function(){m.propFix[this.toLowerCase()]=this}),k.enctype||(m.propFix.enctype="encoding");var uc=/[\t\r\n\f]/g;m.fn.extend({addClass:function(a){var b,c,d,e,f,g,h=0,i=this.length,j="string"==typeof a&&a;if(m.isFunction(a))return this.each(function(b){m(this).addClass(a.call(this,b,this.className))});if(j)for(b=(a||"").match(E)||[];i>h;h++)if(c=this[h],d=1===c.nodeType&&(c.className?(" "+c.className+" ").replace(uc," "):" ")){f=0;while(e=b[f++])d.indexOf(" "+e+" ")<0&&(d+=e+" ");g=m.trim(d),c.className!==g&&(c.className=g)}return this},removeClass:function(a){var b,c,d,e,f,g,h=0,i=this.length,j=0===arguments.length||"string"==typeof a&&a;if(m.isFunction(a))return this.each(function(b){m(this).removeClass(a.call(this,b,this.className))});if(j)for(b=(a||"").match(E)||[];i>h;h++)if(c=this[h],d=1===c.nodeType&&(c.className?(" "+c.className+" ").replace(uc," "):"")){f=0;while(e=b[f++])while(d.indexOf(" "+e+" ")>=0)d=d.replace(" "+e+" "," ");g=a?m.trim(d):"",c.className!==g&&(c.className=g)}return this},toggleClass:function(a,b){var c=typeof a;return"boolean"==typeof b&&"string"===c?b?this.addClass(a):this.removeClass(a):this.each(m.isFunction(a)?function(c){m(this).toggleClass(a.call(this,c,this.className,b),b)}:function(){if("string"===c){var b,d=0,e=m(this),f=a.match(E)||[];while(b=f[d++])e.hasClass(b)?e.removeClass(b):e.addClass(b)}else(c===K||"boolean"===c)&&(this.className&&m._data(this,"__className__",this.className),this.className=this.className||a===!1?"":m._data(this,"__className__")||"")})},hasClass:function(a){for(var b=" "+a+" ",c=0,d=this.length;d>c;c++)if(1===this[c].nodeType&&(" "+this[c].className+" ").replace(uc," ").indexOf(b)>=0)return!0;return!1}}),m.each("blur focus focusin focusout load resize scroll unload click dblclick mousedown mouseup mousemove mouseover mouseout mouseenter mouseleave change select submit keydown keypress keyup error contextmenu".split(" "),function(a,b){m.fn[b]=function(a,c){return arguments.length>0?this.on(b,null,a,c):this.trigger(b)}}),m.fn.extend({hover:function(a,b){return this.mouseenter(a).mouseleave(b||a)},bind:function(a,b,c){return this.on(a,null,b,c)},unbind:function(a,b){return this.off(a,null,b)},delegate:function(a,b,c,d){return this.on(b,a,c,d)},undelegate:function(a,b,c){return 1===arguments.length?this.off(a,"**"):this.off(b,a||"**",c)}});var vc=m.now(),wc=/\?/,xc=/(,)|(\[|{)|(}|])|"(?:[^"\\\r\n]|\\["\\\/bfnrt]|\\u[\da-fA-F]{4})*"\s*:?|true|false|null|-?(?!0\d)\d+(?:\.\d+|)(?:[eE][+-]?\d+|)/g;m.parseJSON=function(b){if(a.JSON&&a.JSON.parse)return a.JSON.parse(b+"");var c,d=null,e=m.trim(b+"");return e&&!m.trim(e.replace(xc,function(a,b,e,f){return c&&b&&(d=0),0===d?a:(c=e||b,d+=!f-!e,"")}))?Function("return "+e)():m.error("Invalid JSON: "+b)},m.parseXML=function(b){var c,d;if(!b||"string"!=typeof b)return null;try{a.DOMParser?(d=new DOMParser,c=d.parseFromString(b,"text/xml")):(c=new ActiveXObject("Microsoft.XMLDOM"),c.async="false",c.loadXML(b))}catch(e){c=void 0}return c&&c.documentElement&&!c.getElementsByTagName("parsererror").length||m.error("Invalid XML: "+b),c};var yc,zc,Ac=/#.*$/,Bc=/([?&])_=[^&]*/,Cc=/^(.*?):[ \t]*([^\r\n]*)\r?$/gm,Dc=/^(?:about|app|app-storage|.+-extension|file|res|widget):$/,Ec=/^(?:GET|HEAD)$/,Fc=/^\/\//,Gc=/^([\w.+-]+:)(?:\/\/(?:[^\/?#]*@|)([^\/?#:]*)(?::(\d+)|)|)/,Hc={},Ic={},Jc="*/".concat("*");try{zc=location.href}catch(Kc){zc=y.createElement("a"),zc.href="",zc=zc.href}yc=Gc.exec(zc.toLowerCase())||[];function Lc(a){return function(b,c){"string"!=typeof b&&(c=b,b="*");var d,e=0,f=b.toLowerCase().match(E)||[];if(m.isFunction(c))while(d=f[e++])"+"===d.charAt(0)?(d=d.slice(1)||"*",(a[d]=a[d]||[]).unshift(c)):(a[d]=a[d]||[]).push(c)}}function Mc(a,b,c,d){var e={},f=a===Ic;function g(h){var i;return e[h]=!0,m.each(a[h]||[],function(a,h){var j=h(b,c,d);return"string"!=typeof j||f||e[j]?f?!(i=j):void 0:(b.dataTypes.unshift(j),g(j),!1)}),i}return g(b.dataTypes[0])||!e["*"]&&g("*")}function Nc(a,b){var c,d,e=m.ajaxSettings.flatOptions||{};for(d in b)void 0!==b[d]&&((e[d]?a:c||(c={}))[d]=b[d]);return c&&m.extend(!0,a,c),a}function Oc(a,b,c){var d,e,f,g,h=a.contents,i=a.dataTypes;while("*"===i[0])i.shift(),void 0===e&&(e=a.mimeType||b.getResponseHeader("Content-Type"));if(e)for(g in h)if(h[g]&&h[g].test(e)){i.unshift(g);break}if(i[0]in c)f=i[0];else{for(g in c){if(!i[0]||a.converters[g+" "+i[0]]){f=g;break}d||(d=g)}f=f||d}return f?(f!==i[0]&&i.unshift(f),c[f]):void 0}function Pc(a,b,c,d){var e,f,g,h,i,j={},k=a.dataTypes.slice();if(k[1])for(g in a.converters)j[g.toLowerCase()]=a.converters[g];f=k.shift();while(f)if(a.responseFields[f]&&(c[a.responseFields[f]]=b),!i&&d&&a.dataFilter&&(b=a.dataFilter(b,a.dataType)),i=f,f=k.shift())if("*"===f)f=i;else if("*"!==i&&i!==f){if(g=j[i+" "+f]||j["* "+f],!g)for(e in j)if(h=e.split(" "),h[1]===f&&(g=j[i+" "+h[0]]||j["* "+h[0]])){g===!0?g=j[e]:j[e]!==!0&&(f=h[0],k.unshift(h[1]));break}if(g!==!0)if(g&&a["throws"])b=g(b);else try{b=g(b)}catch(l){return{state:"parsererror",error:g?l:"No conversion from "+i+" to "+f}}}return{state:"success",data:b}}m.extend({active:0,lastModified:{},etag:{},ajaxSettings:{url:zc,type:"GET",isLocal:Dc.test(yc[1]),global:!0,processData:!0,async:!0,contentType:"application/x-www-form-urlencoded; charset=UTF-8",accepts:{"*":Jc,text:"text/plain",html:"text/html",xml:"application/xml, text/xml",json:"application/json, text/javascript"},contents:{xml:/xml/,html:/html/,json:/json/},responseFields:{xml:"responseXML",text:"responseText",json:"responseJSON"},converters:{"* text":String,"text html":!0,"text json":m.parseJSON,"text xml":m.parseXML},flatOptions:{url:!0,context:!0}},ajaxSetup:function(a,b){return b?Nc(Nc(a,m.ajaxSettings),b):Nc(m.ajaxSettings,a)},ajaxPrefilter:Lc(Hc),ajaxTransport:Lc(Ic),ajax:function(a,b){"object"==typeof a&&(b=a,a=void 0),b=b||{};var c,d,e,f,g,h,i,j,k=m.ajaxSetup({},b),l=k.context||k,n=k.context&&(l.nodeType||l.jquery)?m(l):m.event,o=m.Deferred(),p=m.Callbacks("once memory"),q=k.statusCode||{},r={},s={},t=0,u="canceled",v={readyState:0,getResponseHeader:function(a){var b;if(2===t){if(!j){j={};while(b=Cc.exec(f))j[b[1].toLowerCase()]=b[2]}b=j[a.toLowerCase()]}return null==b?null:b},getAllResponseHeaders:function(){return 2===t?f:null},setRequestHeader:function(a,b){var c=a.toLowerCase();return t||(a=s[c]=s[c]||a,r[a]=b),this},overrideMimeType:function(a){return t||(k.mimeType=a),this},statusCode:function(a){var b;if(a)if(2>t)for(b in a)q[b]=[q[b],a[b]];else v.always(a[v.status]);return this},abort:function(a){var b=a||u;return i&&i.abort(b),x(0,b),this}};if(o.promise(v).complete=p.add,v.success=v.done,v.error=v.fail,k.url=((a||k.url||zc)+"").replace(Ac,"").replace(Fc,yc[1]+"//"),k.type=b.method||b.type||k.method||k.type,k.dataTypes=m.trim(k.dataType||"*").toLowerCase().match(E)||[""],null==k.crossDomain&&(c=Gc.exec(k.url.toLowerCase()),k.crossDomain=!(!c||c[1]===yc[1]&&c[2]===yc[2]&&(c[3]||("http:"===c[1]?"80":"443"))===(yc[3]||("http:"===yc[1]?"80":"443")))),k.data&&k.processData&&"string"!=typeof k.data&&(k.data=m.param(k.data,k.traditional)),Mc(Hc,k,b,v),2===t)return v;h=m.event&&k.global,h&&0===m.active++&&m.event.trigger("ajaxStart"),k.type=k.type.toUpperCase(),k.hasContent=!Ec.test(k.type),e=k.url,k.hasContent||(k.data&&(e=k.url+=(wc.test(e)?"&":"?")+k.data,delete k.data),k.cache===!1&&(k.url=Bc.test(e)?e.replace(Bc,"$1_="+vc++):e+(wc.test(e)?"&":"?")+"_="+vc++)),k.ifModified&&(m.lastModified[e]&&v.setRequestHeader("If-Modified-Since",m.lastModified[e]),m.etag[e]&&v.setRequestHeader("If-None-Match",m.etag[e])),(k.data&&k.hasContent&&k.contentType!==!1||b.contentType)&&v.setRequestHeader("Content-Type",k.contentType),v.setRequestHeader("Accept",k.dataTypes[0]&&k.accepts[k.dataTypes[0]]?k.accepts[k.dataTypes[0]]+("*"!==k.dataTypes[0]?", "+Jc+"; q=0.01":""):k.accepts["*"]);for(d in k.headers)v.setRequestHeader(d,k.headers[d]);if(k.beforeSend&&(k.beforeSend.call(l,v,k)===!1||2===t))return v.abort();u="abort";for(d in{success:1,error:1,complete:1})v[d](k[d]);if(i=Mc(Ic,k,b,v)){v.readyState=1,h&&n.trigger("ajaxSend",[v,k]),k.async&&k.timeout>0&&(g=setTimeout(function(){v.abort("timeout")},k.timeout));try{t=1,i.send(r,x)}catch(w){if(!(2>t))throw w;x(-1,w)}}else x(-1,"No Transport");function x(a,b,c,d){var j,r,s,u,w,x=b;2!==t&&(t=2,g&&clearTimeout(g),i=void 0,f=d||"",v.readyState=a>0?4:0,j=a>=200&&300>a||304===a,c&&(u=Oc(k,v,c)),u=Pc(k,u,v,j),j?(k.ifModified&&(w=v.getResponseHeader("Last-Modified"),w&&(m.lastModified[e]=w),w=v.getResponseHeader("etag"),w&&(m.etag[e]=w)),204===a||"HEAD"===k.type?x="nocontent":304===a?x="notmodified":(x=u.state,r=u.data,s=u.error,j=!s)):(s=x,(a||!x)&&(x="error",0>a&&(a=0))),v.status=a,v.statusText=(b||x)+"",j?o.resolveWith(l,[r,x,v]):o.rejectWith(l,[v,x,s]),v.statusCode(q),q=void 0,h&&n.trigger(j?"ajaxSuccess":"ajaxError",[v,k,j?r:s]),p.fireWith(l,[v,x]),h&&(n.trigger("ajaxComplete",[v,k]),--m.active||m.event.trigger("ajaxStop")))}return v},getJSON:function(a,b,c){return m.get(a,b,c,"json")},getScript:function(a,b){return m.get(a,void 0,b,"script")}}),m.each(["get","post"],function(a,b){m[b]=function(a,c,d,e){return m.isFunction(c)&&(e=e||d,d=c,c=void 0),m.ajax({url:a,type:b,dataType:e,data:c,success:d})}}),m._evalUrl=function(a){return m.ajax({url:a,type:"GET",dataType:"script",async:!1,global:!1,"throws":!0})},m.fn.extend({wrapAll:function(a){if(m.isFunction(a))return this.each(function(b){m(this).wrapAll(a.call(this,b))});if(this[0]){var b=m(a,this[0].ownerDocument).eq(0).clone(!0);this[0].parentNode&&b.insertBefore(this[0]),b.map(function(){var a=this;while(a.firstChild&&1===a.firstChild.nodeType)a=a.firstChild;return a}).append(this)}return this},wrapInner:function(a){return this.each(m.isFunction(a)?function(b){m(this).wrapInner(a.call(this,b))}:function(){var b=m(this),c=b.contents();c.length?c.wrapAll(a):b.append(a)})},wrap:function(a){var b=m.isFunction(a);return this.each(function(c){m(this).wrapAll(b?a.call(this,c):a)})},unwrap:function(){return this.parent().each(function(){m.nodeName(this,"body")||m(this).replaceWith(this.childNodes)}).end()}}),m.expr.filters.hidden=function(a){return a.offsetWidth<=0&&a.offsetHeight<=0||!k.reliableHiddenOffsets()&&"none"===(a.style&&a.style.display||m.css(a,"display"))},m.expr.filters.visible=function(a){return!m.expr.filters.hidden(a)};var Qc=/%20/g,Rc=/\[\]$/,Sc=/\r?\n/g,Tc=/^(?:submit|button|image|reset|file)$/i,Uc=/^(?:input|select|textarea|keygen)/i;function Vc(a,b,c,d){var e;if(m.isArray(b))m.each(b,function(b,e){c||Rc.test(a)?d(a,e):Vc(a+"["+("object"==typeof e?b:"")+"]",e,c,d)});else if(c||"object"!==m.type(b))d(a,b);else for(e in b)Vc(a+"["+e+"]",b[e],c,d)}m.param=function(a,b){var c,d=[],e=function(a,b){b=m.isFunction(b)?b():null==b?"":b,d[d.length]=encodeURIComponent(a)+"="+encodeURIComponent(b)};if(void 0===b&&(b=m.ajaxSettings&&m.ajaxSettings.traditional),m.isArray(a)||a.jquery&&!m.isPlainObject(a))m.each(a,function(){e(this.name,this.value)});else for(c in a)Vc(c,a[c],b,e);return d.join("&").replace(Qc,"+")},m.fn.extend({serialize:function(){return m.param(this.serializeArray())},serializeArray:function(){return this.map(function(){var a=m.prop(this,"elements");return a?m.makeArray(a):this}).filter(function(){var a=this.type;return this.name&&!m(this).is(":disabled")&&Uc.test(this.nodeName)&&!Tc.test(a)&&(this.checked||!W.test(a))}).map(function(a,b){var c=m(this).val();return null==c?null:m.isArray(c)?m.map(c,function(a){return{name:b.name,value:a.replace(Sc,"\r\n")}}):{name:b.name,value:c.replace(Sc,"\r\n")}}).get()}}),m.ajaxSettings.xhr=void 0!==a.ActiveXObject?function(){return!this.isLocal&&/^(get|post|head|put|delete|options)$/i.test(this.type)&&Zc()||$c()}:Zc;var Wc=0,Xc={},Yc=m.ajaxSettings.xhr();a.attachEvent&&a.attachEvent("onunload",function(){for(var a in Xc)Xc[a](void 0,!0)}),k.cors=!!Yc&&"withCredentials"in Yc,Yc=k.ajax=!!Yc,Yc&&m.ajaxTransport(function(a){if(!a.crossDomain||k.cors){var b;return{send:function(c,d){var e,f=a.xhr(),g=++Wc;if(f.open(a.type,a.url,a.async,a.username,a.password),a.xhrFields)for(e in a.xhrFields)f[e]=a.xhrFields[e];a.mimeType&&f.overrideMimeType&&f.overrideMimeType(a.mimeType),a.crossDomain||c["X-Requested-With"]||(c["X-Requested-With"]="XMLHttpRequest");for(e in c)void 0!==c[e]&&f.setRequestHeader(e,c[e]+"");f.send(a.hasContent&&a.data||null),b=function(c,e){var h,i,j;if(b&&(e||4===f.readyState))if(delete Xc[g],b=void 0,f.onreadystatechange=m.noop,e)4!==f.readyState&&f.abort();else{j={},h=f.status,"string"==typeof f.responseText&&(j.text=f.responseText);try{i=f.statusText}catch(k){i=""}h||!a.isLocal||a.crossDomain?1223===h&&(h=204):h=j.text?200:404}j&&d(h,i,j,f.getAllResponseHeaders())},a.async?4===f.readyState?setTimeout(b):f.onreadystatechange=Xc[g]=b:b()},abort:function(){b&&b(void 0,!0)}}}});function Zc(){try{return new a.XMLHttpRequest}catch(b){}}function $c(){try{return new a.ActiveXObject("Microsoft.XMLHTTP")}catch(b){}}m.ajaxSetup({accepts:{script:"text/javascript, application/javascript, application/ecmascript, application/x-ecmascript"},contents:{script:/(?:java|ecma)script/},converters:{"text script":function(a){return m.globalEval(a),a}}}),m.ajaxPrefilter("script",function(a){void 0===a.cache&&(a.cache=!1),a.crossDomain&&(a.type="GET",a.global=!1)}),m.ajaxTransport("script",function(a){if(a.crossDomain){var b,c=y.head||m("head")[0]||y.documentElement;return{send:function(d,e){b=y.createElement("script"),b.async=!0,a.scriptCharset&&(b.charset=a.scriptCharset),b.src=a.url,b.onload=b.onreadystatechange=function(a,c){(c||!b.readyState||/loaded|complete/.test(b.readyState))&&(b.onload=b.onreadystatechange=null,b.parentNode&&b.parentNode.removeChild(b),b=null,c||e(200,"success"))},c.insertBefore(b,c.firstChild)},abort:function(){b&&b.onload(void 0,!0)}}}});var _c=[],ad=/(=)\?(?=&|$)|\?\?/;m.ajaxSetup({jsonp:"callback",jsonpCallback:function(){var a=_c.pop()||m.expando+"_"+vc++;return this[a]=!0,a}}),m.ajaxPrefilter("json jsonp",function(b,c,d){var e,f,g,h=b.jsonp!==!1&&(ad.test(b.url)?"url":"string"==typeof b.data&&!(b.contentType||"").indexOf("application/x-www-form-urlencoded")&&ad.test(b.data)&&"data");return h||"jsonp"===b.dataTypes[0]?(e=b.jsonpCallback=m.isFunction(b.jsonpCallback)?b.jsonpCallback():b.jsonpCallback,h?b[h]=b[h].replace(ad,"$1"+e):b.jsonp!==!1&&(b.url+=(wc.test(b.url)?"&":"?")+b.jsonp+"="+e),b.converters["script json"]=function(){return g||m.error(e+" was not called"),g[0]},b.dataTypes[0]="json",f=a[e],a[e]=function(){g=arguments},d.always(function(){a[e]=f,b[e]&&(b.jsonpCallback=c.jsonpCallback,_c.push(e)),g&&m.isFunction(f)&&f(g[0]),g=f=void 0}),"script"):void 0}),m.parseHTML=function(a,b,c){if(!a||"string"!=typeof a)return null;"boolean"==typeof b&&(c=b,b=!1),b=b||y;var d=u.exec(a),e=!c&&[];return d?[b.createElement(d[1])]:(d=m.buildFragment([a],b,e),e&&e.length&&m(e).remove(),m.merge([],d.childNodes))};var bd=m.fn.load;m.fn.load=function(a,b,c){if("string"!=typeof a&&bd)return bd.apply(this,arguments);var d,e,f,g=this,h=a.indexOf(" ");return h>=0&&(d=m.trim(a.slice(h,a.length)),a=a.slice(0,h)),m.isFunction(b)?(c=b,b=void 0):b&&"object"==typeof b&&(f="POST"),g.length>0&&m.ajax({url:a,type:f,dataType:"html",data:b}).done(function(a){e=arguments,g.html(d?m("<div>").append(m.parseHTML(a)).find(d):a)}).complete(c&&function(a,b){g.each(c,e||[a.responseText,b,a])}),this},m.each(["ajaxStart","ajaxStop","ajaxComplete","ajaxError","ajaxSuccess","ajaxSend"],function(a,b){m.fn[b]=function(a){return this.on(b,a)}}),m.expr.filters.animated=function(a){return m.grep(m.timers,function(b){return a===b.elem}).length};var cd=a.document.documentElement;function dd(a){return m.isWindow(a)?a:9===a.nodeType?a.defaultView||a.parentWindow:!1}m.offset={setOffset:function(a,b,c){var d,e,f,g,h,i,j,k=m.css(a,"position"),l=m(a),n={};"static"===k&&(a.style.position="relative"),h=l.offset(),f=m.css(a,"top"),i=m.css(a,"left"),j=("absolute"===k||"fixed"===k)&&m.inArray("auto",[f,i])>-1,j?(d=l.position(),g=d.top,e=d.left):(g=parseFloat(f)||0,e=parseFloat(i)||0),m.isFunction(b)&&(b=b.call(a,c,h)),null!=b.top&&(n.top=b.top-h.top+g),null!=b.left&&(n.left=b.left-h.left+e),"using"in b?b.using.call(a,n):l.css(n)}},m.fn.extend({offset:function(a){if(arguments.length)return void 0===a?this:this.each(function(b){m.offset.setOffset(this,a,b)});var b,c,d={top:0,left:0},e=this[0],f=e&&e.ownerDocument;if(f)return b=f.documentElement,m.contains(b,e)?(typeof e.getBoundingClientRect!==K&&(d=e.getBoundingClientRect()),c=dd(f),{top:d.top+(c.pageYOffset||b.scrollTop)-(b.clientTop||0),left:d.left+(c.pageXOffset||b.scrollLeft)-(b.clientLeft||0)}):d},position:function(){if(this[0]){var a,b,c={top:0,left:0},d=this[0];return"fixed"===m.css(d,"position")?b=d.getBoundingClientRect():(a=this.offsetParent(),b=this.offset(),m.nodeName(a[0],"html")||(c=a.offset()),c.top+=m.css(a[0],"borderTopWidth",!0),c.left+=m.css(a[0],"borderLeftWidth",!0)),{top:b.top-c.top-m.css(d,"marginTop",!0),left:b.left-c.left-m.css(d,"marginLeft",!0)}}},offsetParent:function(){return this.map(function(){var a=this.offsetParent||cd;while(a&&!m.nodeName(a,"html")&&"static"===m.css(a,"position"))a=a.offsetParent;return a||cd})}}),m.each({scrollLeft:"pageXOffset",scrollTop:"pageYOffset"},function(a,b){var c=/Y/.test(b);m.fn[a]=function(d){return V(this,function(a,d,e){var f=dd(a);return void 0===e?f?b in f?f[b]:f.document.documentElement[d]:a[d]:void(f?f.scrollTo(c?m(f).scrollLeft():e,c?e:m(f).scrollTop()):a[d]=e)},a,d,arguments.length,null)}}),m.each(["top","left"],function(a,b){m.cssHooks[b]=Lb(k.pixelPosition,function(a,c){return c?(c=Jb(a,b),Hb.test(c)?m(a).position()[b]+"px":c):void 0})}),m.each({Height:"height",Width:"width"},function(a,b){m.each({padding:"inner"+a,content:b,"":"outer"+a},function(c,d){m.fn[d]=function(d,e){var f=arguments.length&&(c||"boolean"!=typeof d),g=c||(d===!0||e===!0?"margin":"border");return V(this,function(b,c,d){var e;return m.isWindow(b)?b.document.documentElement["client"+a]:9===b.nodeType?(e=b.documentElement,Math.max(b.body["scroll"+a],e["scroll"+a],b.body["offset"+a],e["offset"+a],e["client"+a])):void 0===d?m.css(b,c,g):m.style(b,c,d,g)},b,f?d:void 0,f,null)}})}),m.fn.size=function(){return this.length},m.fn.andSelf=m.fn.addBack,"function"==typeof define&&define.amd&&define("jquery",[],function(){return m});var ed=a.jQuery,fd=a.$;return m.noConflict=function(b){return a.$===m&&(a.$=fd),b&&a.jQuery===m&&(a.jQuery=ed),m},typeof b===K&&(a.jQuery=a.$=m),m});
//# sourceMappingURL=jquery.min.map
```
