## Supplementary material for "Chromatin interaction analysis with updated ChIA-PET Tool (V3)": script.js

```
( function( $ ) {
$( document ).ready(function() {
$(document).ready(function(){

$('#menu > ul > li ul').each(function(index, e){
  var count = $(e).find('li').length;
  var content = '<span class=\"cnt\">' + count + '</span>';
  $(e).closest('li').children('a').append(content);
});
$('#menu ul ul li:odd').addClass('odd');
$('#menu ul ul li:even').addClass('even');
$('#menu > ul > li > a').click(function() {
  $('#menu li').removeClass('active');
  $(this).closest('li').addClass('active');	
  var checkElement = $(this).next();
  if((checkElement.is('ul')) && (checkElement.is(':visible'))) {
    $(this).closest('li').removeClass('active');
    checkElement.slideUp('normal');
  }
  if((checkElement.is('ul')) && (!checkElement.is(':visible'))) {
    $('#menu ul ul:visible').slideUp('normal');
    checkElement.slideDown('normal');
  }
  if($(this).closest('li').find('ul').children().length == 0) {
    return true;
  } else {
    return false;	
  }		
});

});

});
} )( jQuery );
```
