## Supplementary figures and images for "Chromatin interaction analysis with updated ChIA-PET Tool (V3)"

### footer-line.png

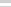

### imgzoom_tb.gif

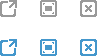

### logo.png

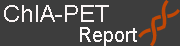

### noize.png

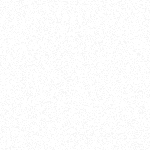

### pic-next.png

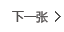

### pic-prev.png

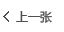

### Rplot4.png

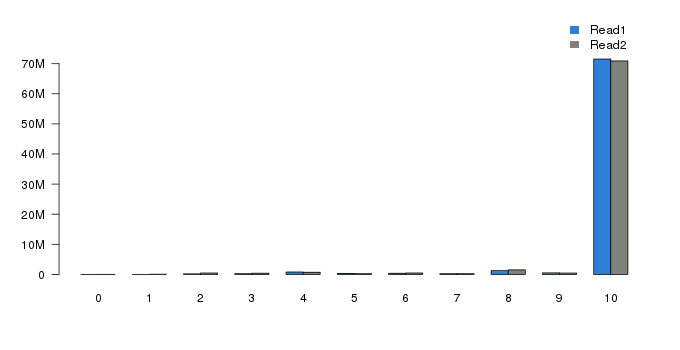

### Rplot5.png

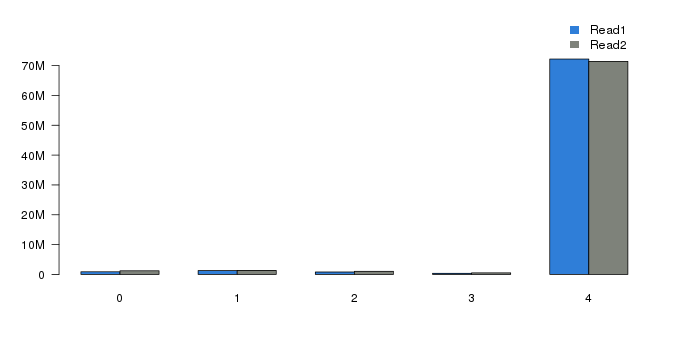

### Rplot6.png

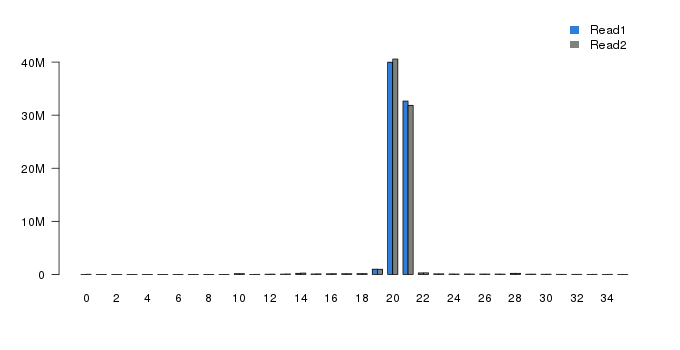

### Rplot8.png

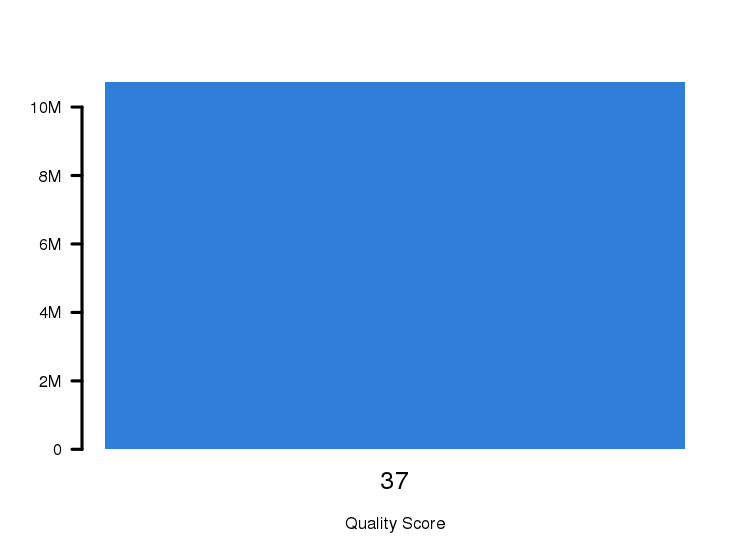

### Rplot9.png

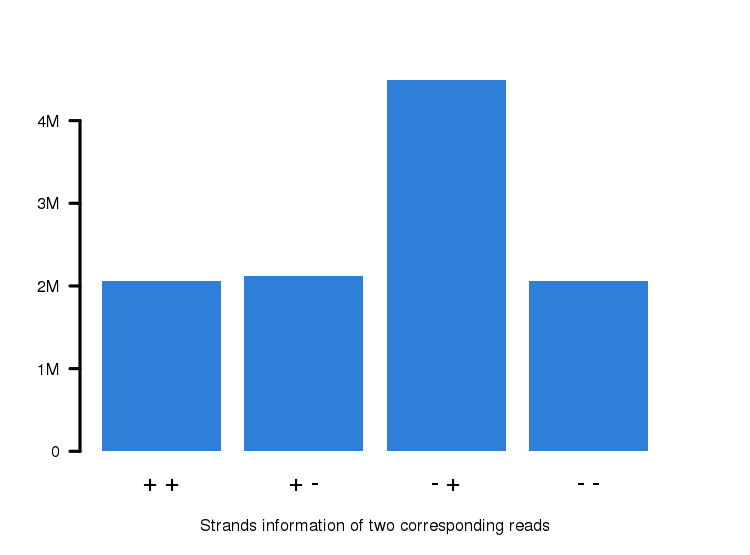

### Rplot10.png

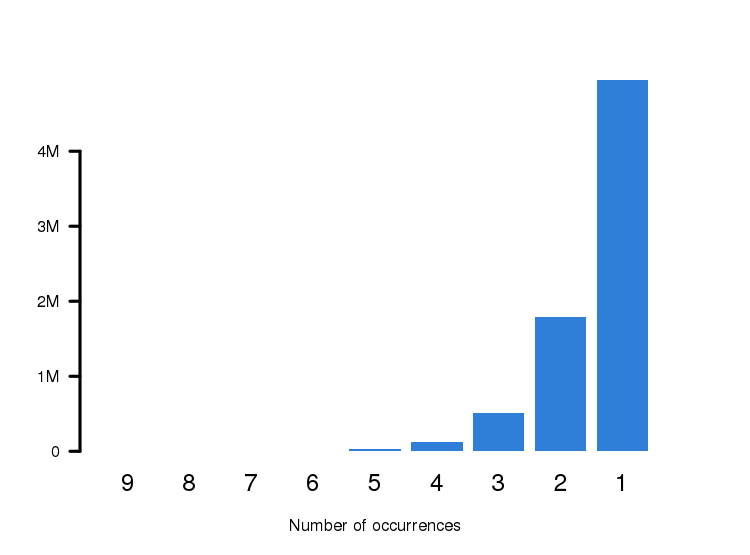

### Rplot11.png

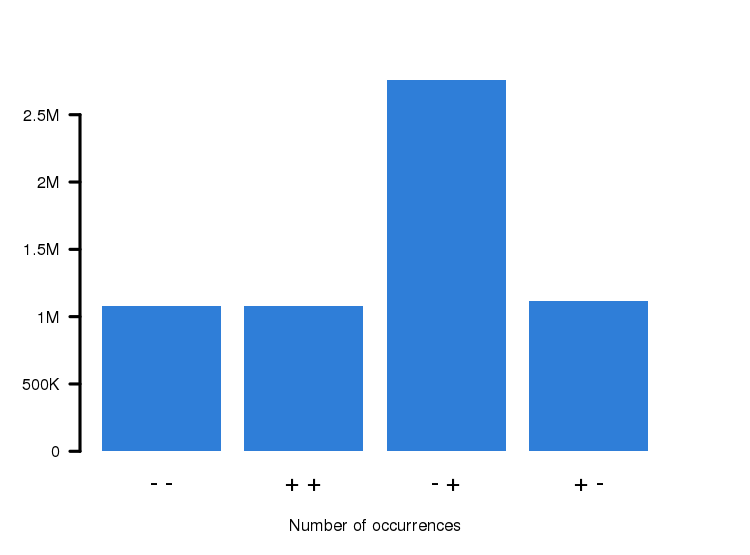

### Rplot12.png

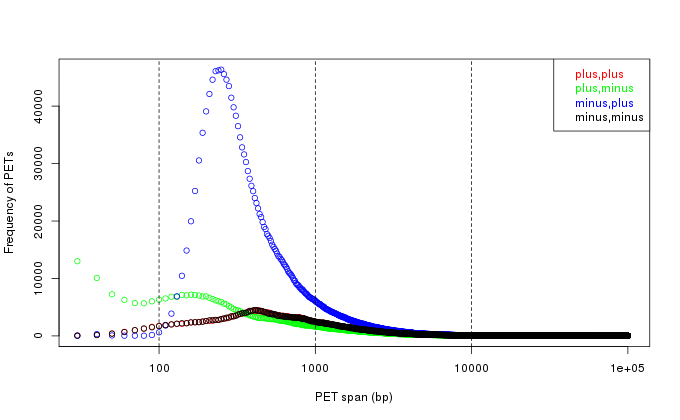

### Rplot14.png

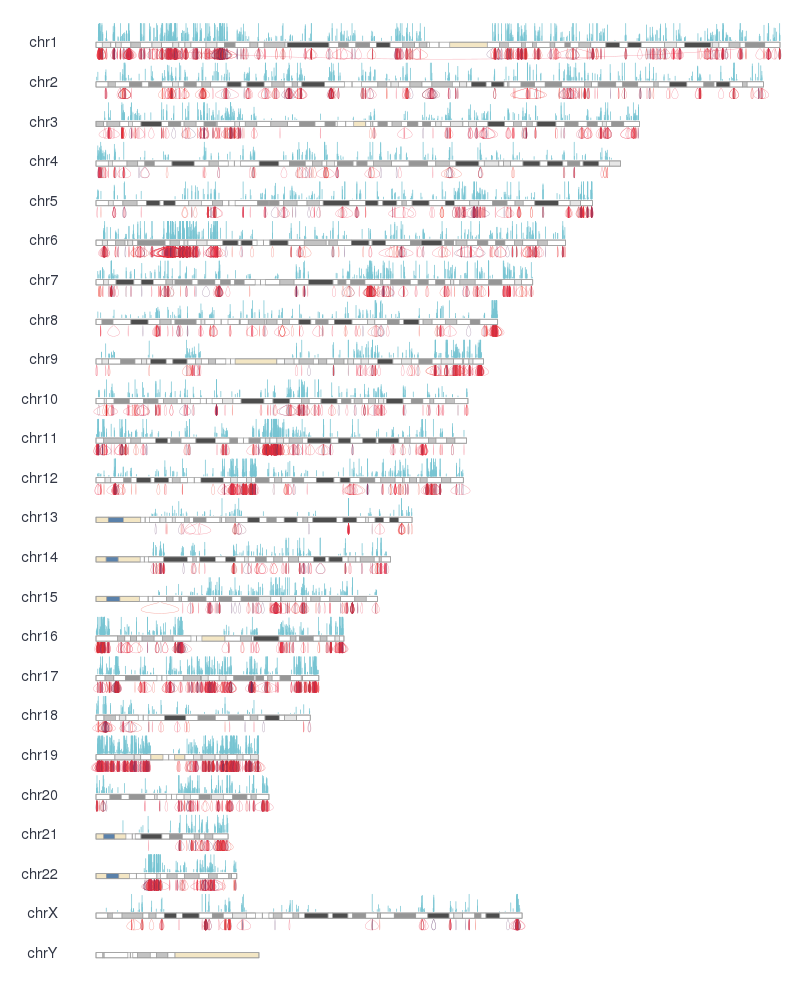

### Rplot15.png

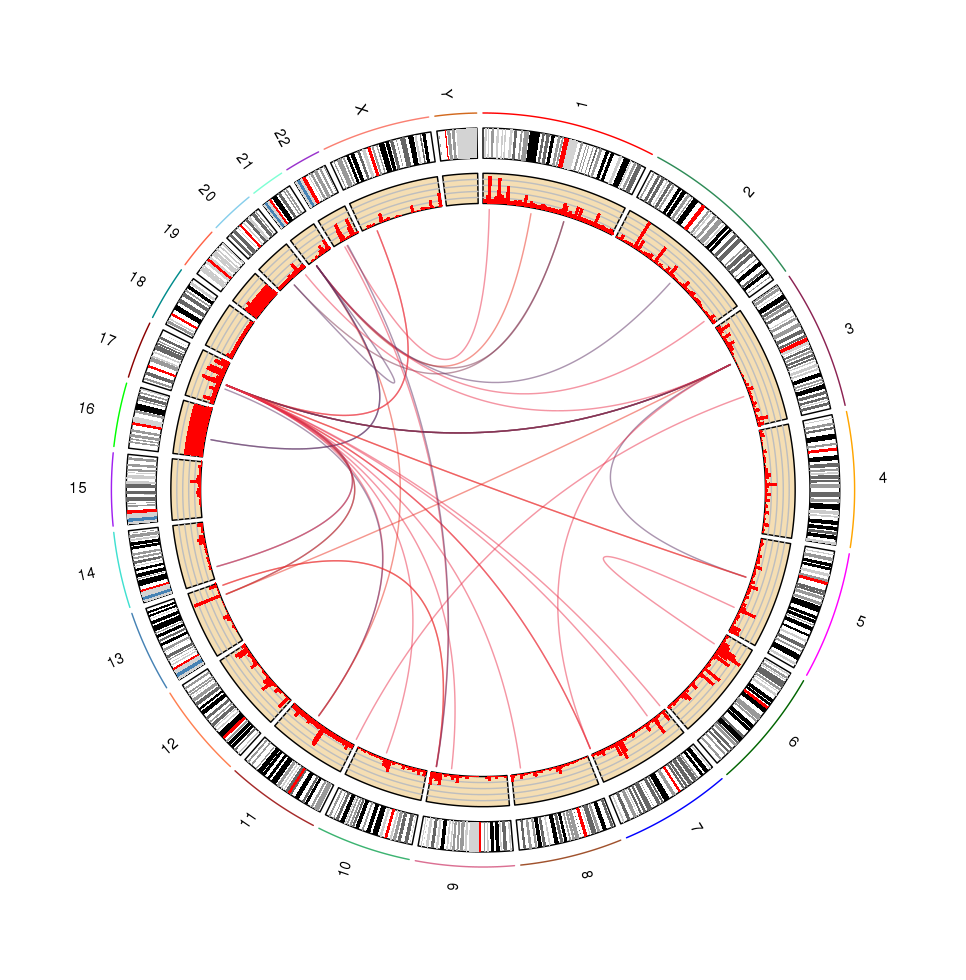

### Rplot16.png

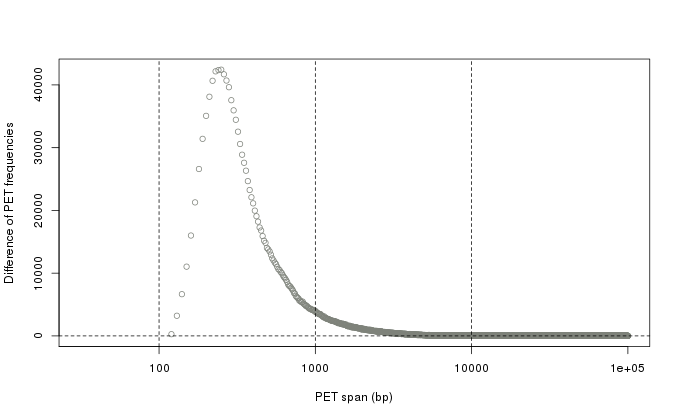

### Rplot24.png

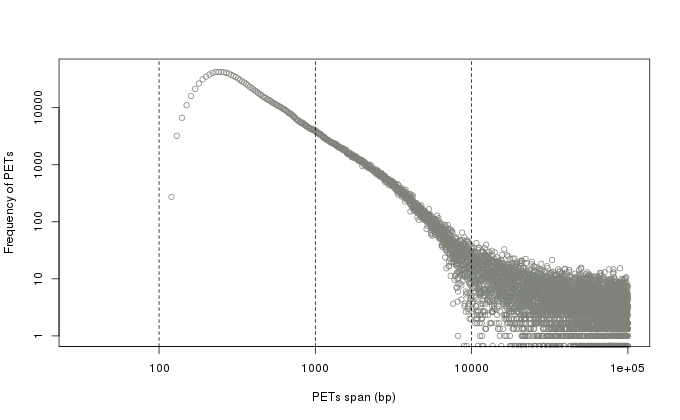
